# Characterizing dysbiosis of gut microbiome in PD: Evidence for overabundance of opportunistic pathogens

**DOI:** 10.1101/2020.01.13.905166

**Authors:** Zachary D Wallen, Mary Appah, Marissa N Dean, Cheryl L Sesler, Stewart A Factor, Eric Molho, Cyrus P Zabetian, David G Standaert, Haydeh Payami

## Abstract

In Parkinson’s disease (PD), gastrointestinal features are common and often precede the motor signs. Braak and colleagues proposed that PD may start in the gut, triggered by a pathogen, and spread to the brain. Numerous studies have examined the gut microbiome in PD, all found it to be altered, but found inconsistent results on associated microorganisms. Studies to date have been small (N=20 to 306) and are difficult to compare or combine due to varied methodology. We conducted a microbiome-wide association study (MWAS) with two large datasets for internal replication (N=333 and 507). We used uniform methodology when possible, interrogated confounders, and applied two statistical tests for concordance, followed by correlation network analysis to infer interactions. Fifteen genera were associated with PD at a microbiome-wide significance level, in both datasets, with both methods, with or without covariate adjustment. The associations were not independent, rather represented 3 clusters of co-occurring microorganisms. Cluster 1 was composed of opportunistic pathogens; all were elevated in PD. Cluster 2 were short-chain-fatty-acid producing bacteria; all were reduced in PD. Cluster 3 were carbohydrate-metabolizing probiotics; elevated in PD. Depletion of anti-inflammatory short-chain-fatty-acid producing bacteria and elevated levels of probiotics are confirmatory. Overabundance of opportunistic pathogens is a novel finding and their identity provides a lead to experimentally test their role in PD.

## Introduction

PD is a common, progressive and debilitating disease which currently cannot be prevented or cured. With the exception of rare genetic forms, the cause of PD is unknown. Many susceptibility loci^1^ and environmental risk factors^2^ have been identified, but each has a modest effect on risk, and none is sufficient to cause disease. Gene-environment interaction studies have not been able to identify a causative combination.^3-6^ The triggers that cause PD are unknown.

The emerging information about the importance of the gut microbiome in human health and disease,^7^ together with the well-established connection between PD and the gut including common and early occurrence of constipation,^8^ inflammation,^9^ and increased gut membrane permeability,^10^ have raised the possibility that microorganisms in the gut may play a role in PD pathogenesis and prompted a fast growing literature on studies conducted in humans and animal models.^11-30^ Every study that has compared the global composition of the gut microbiome in PD vs. controls found it to be significantly altered; in contrast, attempts to identify PD-associated microorganisms have produced inconsistent results.^31,32^ Low reproducibility has been attributed to small sample sizes (missing true associations due to low power), relaxed statistical thresholds (inflating false positive results), and publishing without a replication dataset (required for genomic studies). Differences in methods of DNA extraction, sequencing, bioinformatics and statistics can all contribute to inter-study variations. The choice of taxonomic resolution for analysis (PD has been tested at all levels from phylum to species) and the inconsistent taxonomic assignments and nomenclature used in various reference databases add to the confusion when comparing results. Last but not least, is confounding by heterogeneity in the populations that were studied: PD is heterogenous and so is the microbiome. PD subtypes cannot be readily identified thus patient populations are inevitably varied. A myriad of factors can affect the microbiome ranging from diet, health and medication to cultural habits, life-styles, race and geography.^33,34^

Identifying microorganisms involved in the dysbiosis of the microbiome is essential for understanding their role in disease. We conducted a hypothesis-free microbiome-wide association study (MWAS) modeled after and using the standards of rigors that are used in genome-wide association studies (GWAS), but with analytic methods that are appropriate for the high-dimensionality and compositionality of the microbiome data. We used two datasets to allow internal replication. The sample sizes in prior PD-microbiome studies have ranged from 10 to 197 PD cases and 10 to 130 controls.^32^ The largest published study (197 cases and 130 controls) is the dataset 1 in the present study, re-analyzed here with a more advanced bioinformatics pipeline than we previously published.^16^ In addition, we present an unpublished independent dataset with 323 cases of PD and 184 controls, analyzed in parallel to dataset 1. Two large data sets allowed for internal replication, and power to detect both rare and common signals. We standardized data collection and processing as much as possible across the two datasets, and for variations that could not be handled in study design, we used statistical techniques to make appropriate adjustments. We used two different statistical tests for MWAS and focused only on results that were reproducibly significant across methods and across datasets. We employed correlation network analysis to infer interactions among PD-associated microorganisms. We were able to confirm some of the previously reported associations with common taxa, and identified novel associations with rare microorganisms that are commensal, but can become opportunistic pathogens in immune-compromised hosts.

## Results

### Dramatic difference between datasets

We discovered a remarkable difference between the two datasets, despite efforts to standardize data collection and analysis (Figure 1). All subjects lived in the United States. Diagnosis, subject selection and data collection were performed by the NeuroGenetics Research Consortium (NGRC) investigators at the four NGRC-affiliated movement disorder clinics, using standardized methods. Dataset 1 (212 PD and 136 controls) was collected in Seattle, WA, Albany, NY, and Atlanta, GA in 2014. Dataset 2 (323 PD and 184 controls) was collected in Birmingham, AL during 2015-2018. Stool was collected using the same kit, DNA was extracted using the same chemistry, and the 16S rRNA gene V4 region was sequenced using the same primers, but in different laboratories, resulting in 10x greater sequence depth in dataset 2 than dataset 1. The same pipeline was used on the two datasets to process the sequences and assign taxonomic classification. Yet, principal component analysis (PCA)^35^ revealed the composition of the microbiome of the samples to be strikingly different in the two datasets (Figure 1), and the difference was statistically significant (P<1E-5, tested using permutational multivariate analysis of variance (PERMANOVA)). The separation of datasets was evident in cases, and in controls, in the same pattern. Greater sequence depth in dataset 2 was a significant contributor to this disparity, but not the sole explanation because the difference between datasets was still significant once sequence depth was adjusted for (PERMANOVA P<1E-5). For all statistical tests (global composition, MWAS, correlations and network analysis), the two datasets were analyzed separately, one, for independent validation, and two, to avoid confounding by mixing two clearly different datasets.

**Figure 1.**
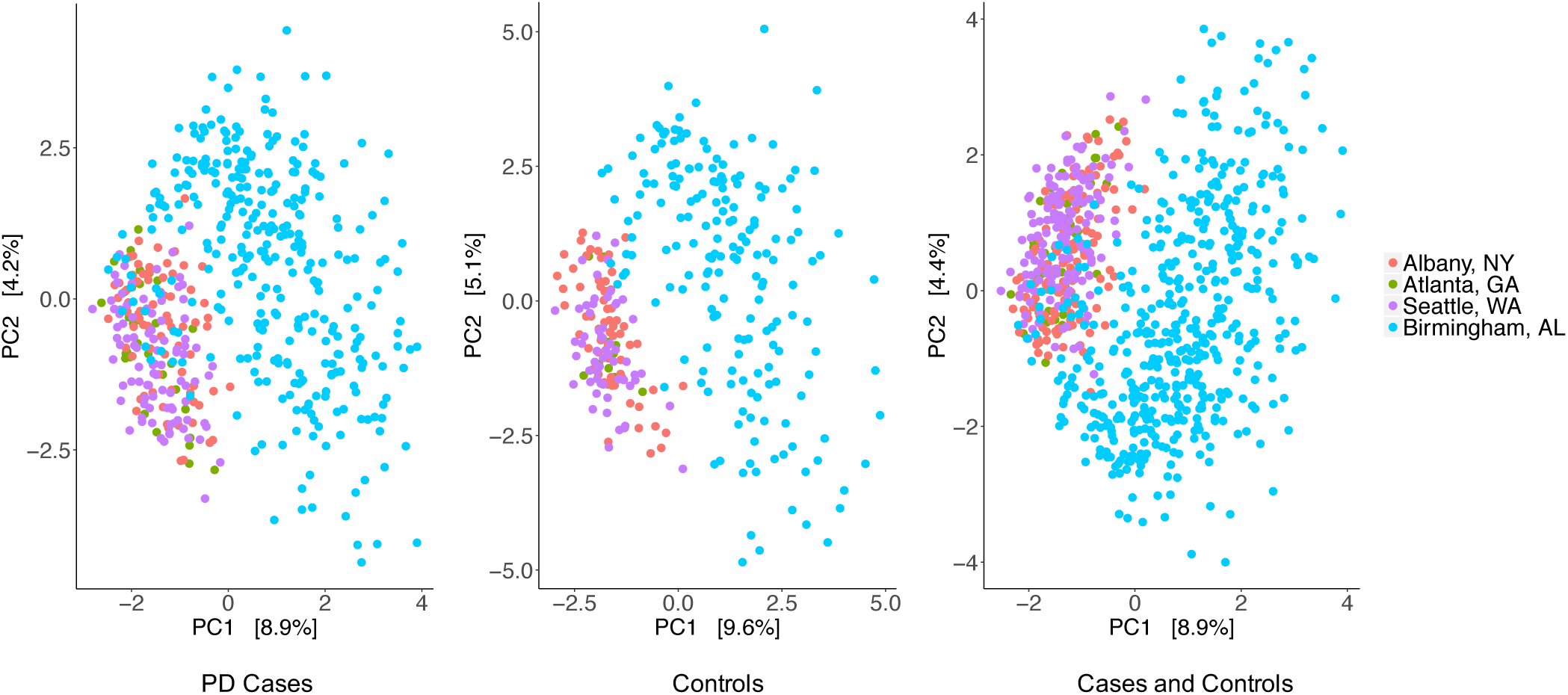
The gut microbiome compositions of the two dataset differed significantly. Principal Component (PC) Analysis was used to generate the graphs for PD cases (left), controls (middle), and cases and controls combined (right) where each point represents the composition of the gut microbiome of one individual and distances indicate degree of similarity to other individuals. Percentages on the x-axis and y-axis correspond to the percent variation in gut microbiome compositions explained by PC1 and PC2. The difference between dataset 1 and dataset 2 was formally tested using PERMANOVA and was significant (P<1E-5). Dataset 1: red (Albany, NY), purple (Seattle, WA) and green (Atlanta, GA). Dataset 2: blue (Birmingham, AL).

### Metadata and Confounders

Metadata were collected using two self-administered questionnaires and medical records (Supplementary Table 1). An Environmental and Family History Questionnaire^4,36^ was used to collect data relevant to PD. A Gut Microbiome Questionnaire^16^ was completed immediately after stool collection and gathered data relevant to the microbiome including diet, gastrointestinal problems, medical conditions, and use of medications. PD medications that subjects were taking at the time of stool collection were extracted from medical records by clinical investigators. The aim of this study was to identify reproducible signals of association between PD and microbiota, and to that end, metadata were used as potential confounders, not as research questions. For example, we did not set out to test the effects of constipation, levodopa or any of the 47 variables listed in Supplementary Table 1 on the microbiome, because, while of interest, that was not the primary aim of the study, and doing so would have reduced the power for the primary aim.

**Table 1.**
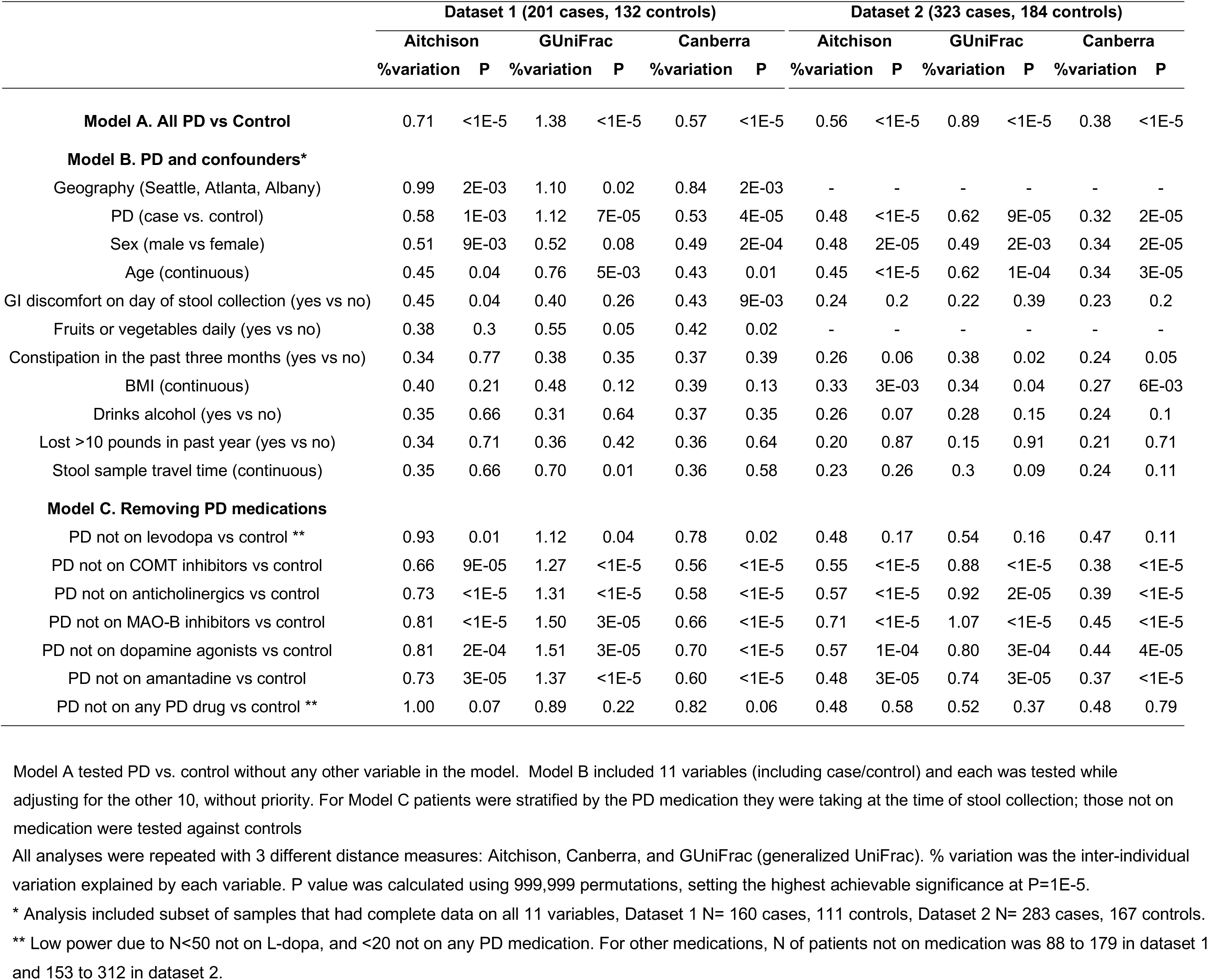
Effect of PD and other key variables on the global composition of microbiome.

To identify which of the variables might confound the study, we tested the distribution of each variable in cases vs. controls, and those that differed at a conservative uncorrected P<0.05 in at least one dataset were tagged as potential confounders (Supplementary Table 1). These included, most notably, constipation in the past 3 months (more common in PD, P=6E-16 dataset 1, P=6E-10 dataset 2) and gastrointestinal discomfort on the day of stool collection (more common in PD, P=2E-9 dataset 1, P=4E-6 dataset 2) as well as sex and age, body mass index (BMI), weight loss, fruits or vegetable intake, alcohol use, and stool sample travel time. These variables, and geographic site, were tested along with case-control status in PERMANOVA (global composition test), and those that were significant were used as covariates in ANCOM (differential abundance test for MWAS). Thus, the results on both the global composition test and PD-associated taxa in MWAS have been adjusted for known potential confounders, except PD medications which had to be handled differently because of collinearity with PD (see section on “Cause of disease or consequence of medication”).

### Global composition of microbiome

First we tested the difference between PD and controls in the global composition of the gut microbiome (β diversity, Table 1). Case vs. control status was tested once by itself, once with all potential confounders in the model in a marginal test where each variable was tested while being adjusted for all others in the model, and once stratified by PD medication (Table 1). To gauge the effect of distance metric on the results, all tests were repeated with Aitchison,^35^ generalized UniFrac (GUniFrac),^37^ and Canberra^38^ distances. Tests were conducted using PERMANOVA^39^ with 99,999 permutations limiting maximum achievable significance to P=1E-5.

PD microbiomes differed significantly from control microbiomes, in both datasets, with every distance metric measured (P<1E-5, Table 1). The PD effect was significant and independent of all analyzed confounders, including geography, constipation, gastrointestinal discomfort, sex, age, BMI, fruit or vegetable intake, alcohol use, and stool sample travel time.

Results were in agreement with population studies in detecting significant effects of sex, age, BMI, gastrointestinal issues and diet on the microbiome,^33,34^ and with other PD studies in detecting evidence for dysbiosis in PD.^11-30^

### Identification of PD-associated microorganisms

To identify PD-associated microorganisms, we conducted MWAS, testing differences between cases and controls in the relative abundances of genera. We conducted MWAS on each dataset separately to test if results replicate, and also to avoid confounding by the heterogeneity between datasets. Each data set was tested with two methods to test analytic concordance: once using analysis of composition of microbiomes (ANCOM)^40^ and again using Kruskal-Wallis rank sum test (KW).^41^ We chose ANCOM because among the numerous methods that have been proposed, ANCOM singularly met three key criteria: incorporates compositionality of the eco-system, allows covariate adjustment, and keeps false positive rate low while maintaining power.^40,42^ Differential abundance was tested hypothesis-free microbiome-wide: ANCOM included all 445 genera detected in dataset 1 and 561 genera in dataset 2; KW included 109 genera in dataset 1 and 163 in dataset 2 (excluding unassigned genera and genera present in <10% of samples). In ANCOM, dataset-specific covariates were included and adjusted for (see MWAS section in Methods). All tests were corrected for multiple testing.

We detected association signals for 15 genera that were microbiome-wide significant by both methods and reproduced robustly in the two datasets, with or without covariate adjustment (Table 2, Figure 2). Five genera had higher abundances in PD than controls: *Porphyromonas, Prevotella, Corynebacterium_1, Bifidobacterium* and *Lactobacillus.* Ten genera had lower abundances in PD than controls: *Faecalibacterium, Agathobacter, Blautia, Roseburia, Fusicatenibacter, Lachnospira, Butyricicoccus, Lachnospiraceae_ND3007_group, Lachnospiraceae_UCG-004*, and *Oscillospira.* Complete MWAS results are in Supplementary Tables 2-5.

**Table 2.**
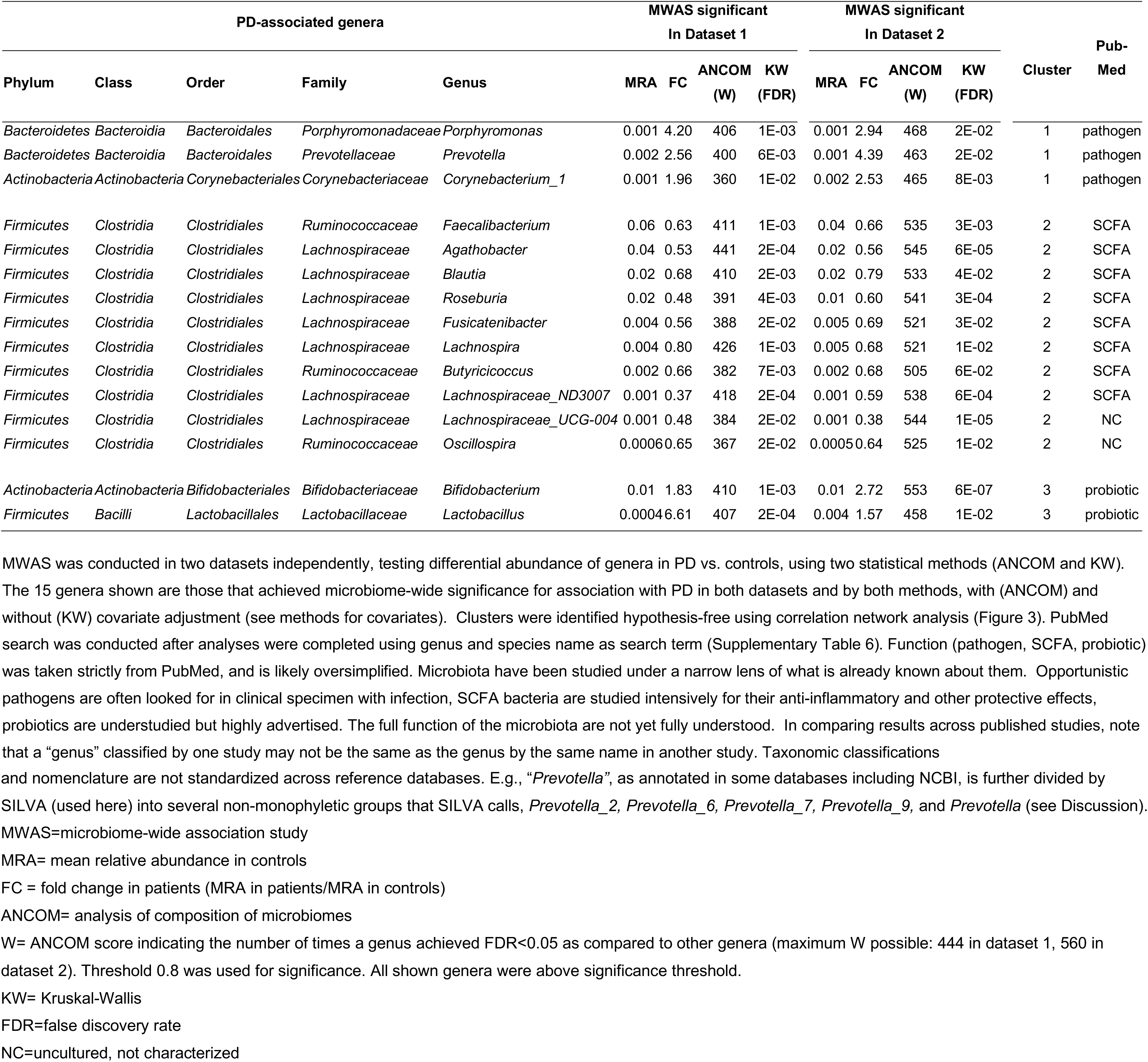
PD-associated genera.

**Figure 2.**
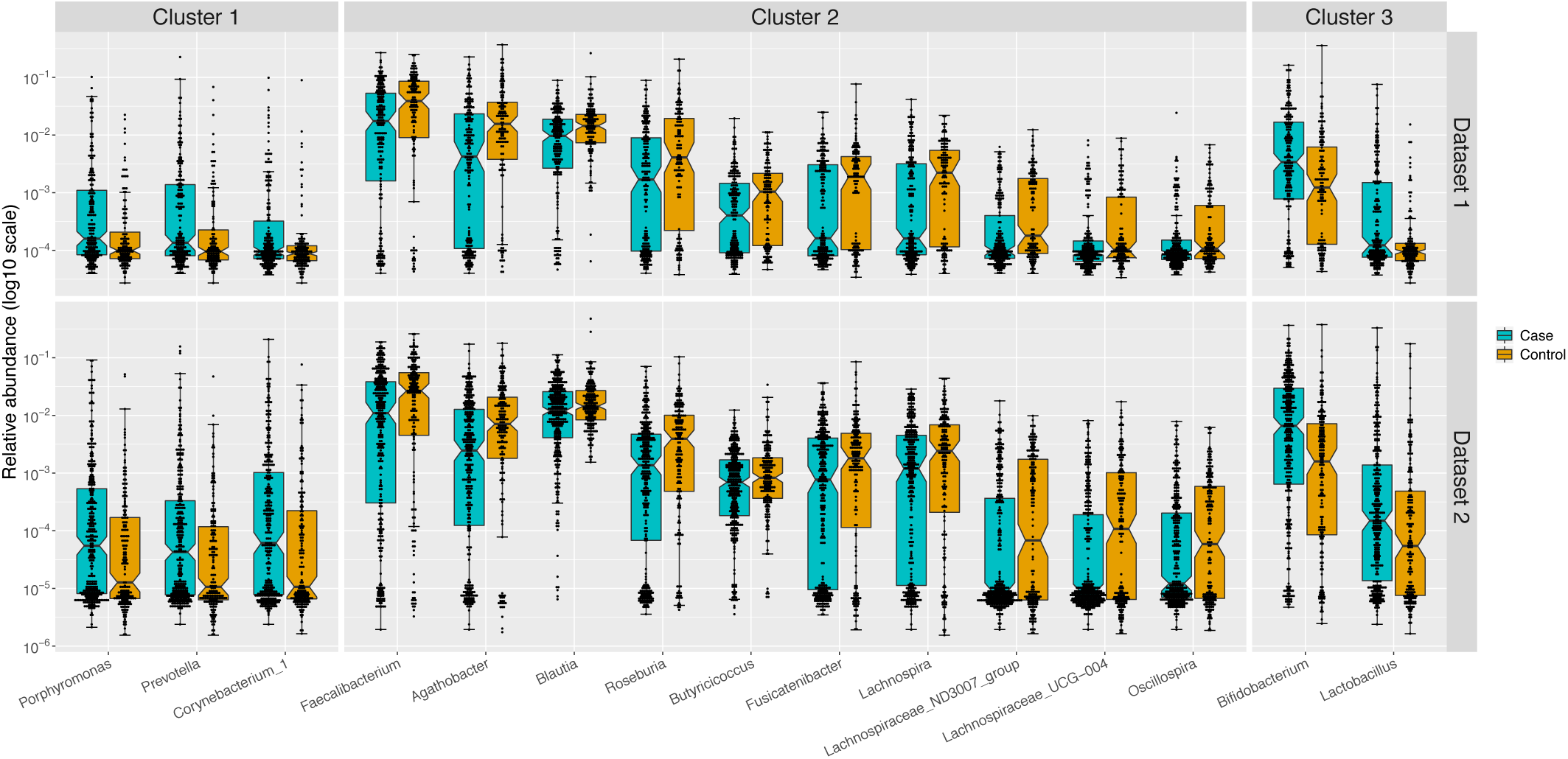
Differential abundances of 15 PD-associated genera replicated in two datasets. Relative abundances in PD cases (blue) and controls (orange) were plotted as log10 scale on the y-axis. Each dot represents a sample, plotted according to the relative abundance of the genus in the sample. The notch in each box indicates the confidence interval of the median. The bottom, middle, and top boundaries of each box represent the first, second (median), and third quartiles of the relative abundances. The whiskers (lines extending from the top and bottom of the box and ending in horizontal cap) extend to points within 1.5 times the interquartile range. The points extending above the whiskers are outliers.

### Correlation network analysis

We questioned if the 15 association signals were independent. We used hypothesis-free correlation network analysis^43^ to infer ecological networks of interacting organisms microbiome-wide (Figure 3, Supplementary Figure 1). The PD-associated genera mapped to three polymicrobial clusters. *Porphyromonas, Prevotella*, and *Corynebacterium_1*, which were elevated in PD, mapped to a community of highly correlated organisms, which we denoted as cluster 1. Cluster 1 was the most distinct cluster in the microbiome with correlations reaching r=0.82 (P<3E-4), the highest in the microbiome in our data. The 10 genera that were depleted in PD formed cluster 2, where eight of them clustered at r≥0.4 (P<3E-4), and remaining two (*Oscillospira* and *Lachnospiraceae_UCG-004)*, clustered with the others at r=0.25 (P<3E-4) and r=0.35 (P<3E-4). *Lactobacillus* and *Bifidobacterium*, both elevated in PD, were correlated with each other at r=0.33 (P<3E-4), which we denoted as cluster 3. Correlations within each cluster were all in the positive direction; i.e., members of clusters 1 tended to increase in abundance together, cluster 2 decreased together, and cluster 3 increased together.

**Figure 3.**
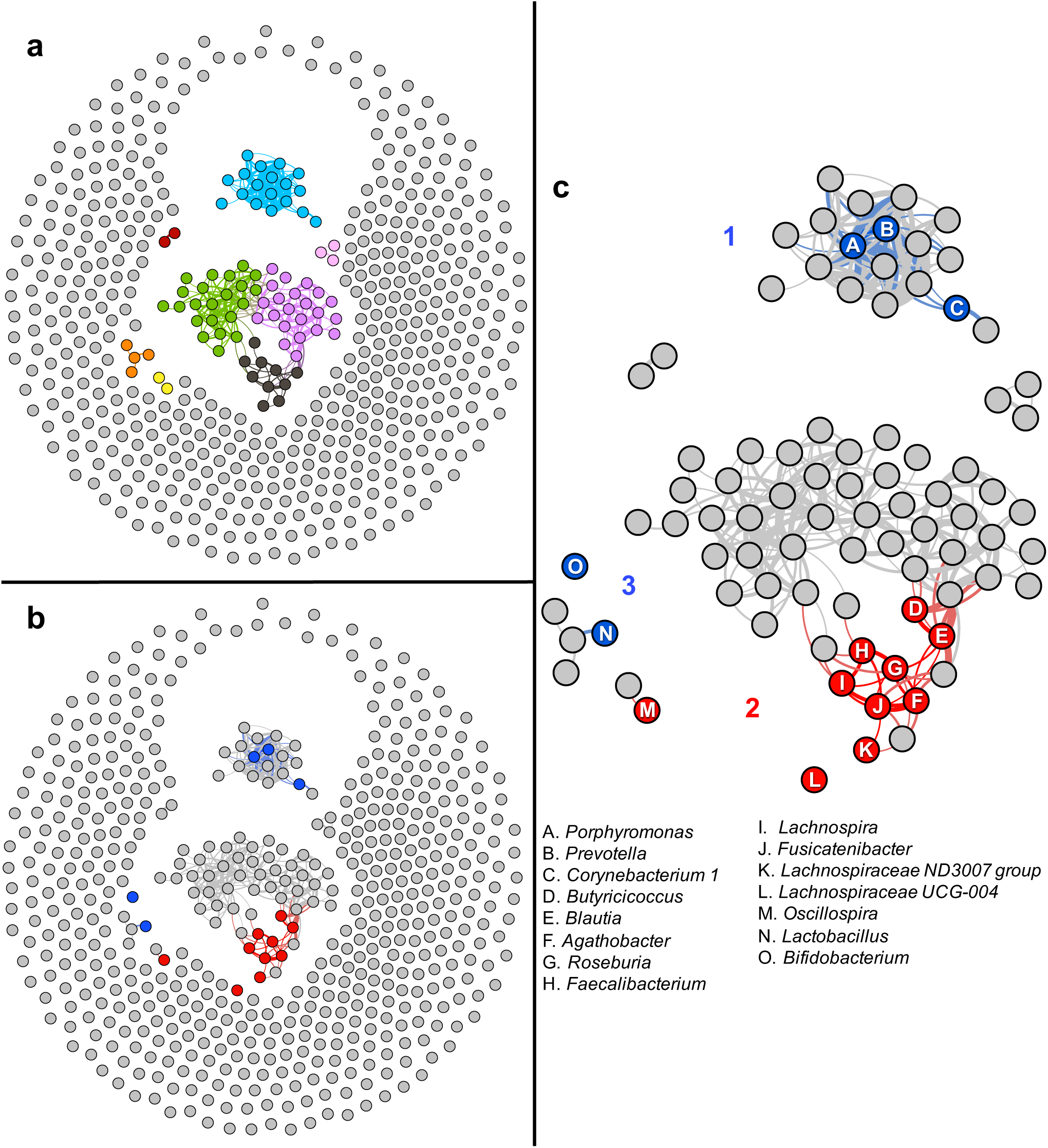
Correlation network analysis mapped PD-associated genera to three polymicrobial clusters. Pairwise correlations in relative abundances were calculated for all genera microbiome-wide and used to detect clusters of co-occurring microorganisms. To display, we used an arbitrary correlation coefficient threshold at r≥ |0.4| to connect the genera that were correlated. All correlations noted were significant at P<3E-4 (the limit for 3,000 permutations). Here we show the result for PD cases in dataset 2 because it had larger sample size and greater sequencing depth than dataset 1. (See Supplementary Figure 1 for cases and controls in dataset 1 and dataset 2). **(a)** Algorithm-detected clusters shown in different colors. **(b)** The algorithm-detected clusters, as in panel a but shown in grey, and PD-associated genera highlighted in blue (if increased in PD) or red (if decreased in PD). **(c)** Zoomed in version of panel b. The 15 PD-associated genera fell in 3 clusters. Cluster 1 was a tightly correlated cluster of microorganisms (r approaching 0.8) which included *Porphyromonas, Prevotella*, and *Corynebacterium_1* (all elevated in PD). Cluster 2 included the 10 genera that were reduced in PD, eight of which are shown connected at r≥0.4, and two are unconnected but correlated significantly (P=3E-4) with the others in the cluster at r=0.25 and r=0.35. *Lactobacillus* and *Bifidobacterium* (correlated at r=0.33 (P<3E-4)) were denoted cluster 3. For unconnected genera (r<0.4), the proximity between nodules does not imply relatedness, for example, *Oscillospira* (M) falls closer to *Lactobacillus* (N) than to *Roseburia* (G) but it is correlated significantly with *Roseburia* (r=0.25, P<3E-4) and not with *Lactobacillus* (r=0.04, P=0.44).

### Functional characteristics

Analyses so far were all hypothesis-free, data-driven, and blind to the functional relevance of the microorganisms. Having identified the associations and their corresponding clusters, we broke the blind by searching PubMed. PubMed results on functional characteristics converged on clusters defined by agnostic network analysis.

#### Genera in cluster 1

*Porphyromonas* and *Prevotella* are anaerobic, gram negative bacteria with lipopolysaccharides (endotoxins) in their outer membrane. They are commensal to the human gastrointestinal and urogenital tracts. *Corynebacterium* are aerobic, gram positive, and have a higher abundance in the skin microbiota than the gut. While commensal and often harmless, *Porphyromonas, Prevotella* and *Corynebacterium* are opportunistic pathogens capable of causing infections in immune-compromised individuals or if they gain access to sterile sites via compromised membranes, post-surgery, bites, or wounds.^44-46^

Many, but not all species of *Porphyromonas, Prevotella*, and *Corynebacterium* are pathogens. *Corynebacterium diphtheriae* is the leading cause of diphtheria. *Porphyromonas gingivalis* causes periodontal disease. We did not detect *C. diphtheriae*, and *P. gingivalis* was extremely rare in our samples. We were interested in knowing the species that made-up these three genera in our PD samples. The bioinformatic pipeline used in our study (DADA2 with SILVA as reference database) assigned the detected sequences (amplicon sequence variants, ASVs) to species if the sequences were 100% identical, otherwise, the ASV was unassigned to species. To confirm and expand on DADA2-SILVA assignments, we blasted all the ASVs that made up each of the three genera against the NCBI 16S rRNA database, focusing only on matches that were >99%-100% identical to a species with high statistical confidence. In PD patients, we found that 80% of *Corynebacterium_1* was composed of one unique ASV with 100% identity to *C. amycolatum* and *C. lactis;* 96% of *Porphyromonas* was composed of ASVs that matched *P. asaccharolytica, P. bennonis, P. somerae* or *P. uenonis* with >99%-100% identity; and 98% of *Prevotella* was composed of ASVs that matched *P. bivia, P. buccalis, P. disiens*, or *P. timonensis* with >99%-100% identity (83% of *Prevotella* matched *P. bivia, P. buccalis, P. disiens*, or *P. timonensis* at 100% identity). We conducted a PubMed search for each of these 10 species, using genus and species name as key word (ex. *Corynebacterium amycolatum*), with search filters: Humans, English, Title/Abstract. Excluding method papers, PubMed returned 104 articles that addressed function, characteristics or relevance to human health, and every article was about the microorganism (search term) as a pathogen in clinical specimens from various infections (Supplementary Table 6).

Clinical specimen from chronic wounds, infections and inflammations are often polymicrobial.^44-46^ *Porphyromonas, Prevotella, Corynebacterium* and other members of cluster 1 are often observed together in these polymicrobial infections.^44-46^ With the newly acquired knowledge on the potential biological significance of cluster 1, we questioned if this polymicrobial group as a whole may be relevant to PD. The co-occurring organisms in cluster 1 (defined by correlation r≥0.4) were *Anaerococcus, Campylobacter, Ezakiella, Finegoldia, Murdochiella, Peptoniphilus, Porphyromonas, Prevotella* and *Varibaculum* in dataset 1, and *Anaerococcus, Campylobacter, Corynebacterium_1, Ezakiella, Fastidiosipila, Finegoldia, Lawsonella, Mobiluncus, Mogibacterium, Murdochiella, Negativicoccus, Peptoniphilus, Porphyromonas, Prevotella, Prevotella_6, S5-A14a, Varibaculum*, and unclassified *Corynebacteriaceae* in dataset 2. Most of these organisms are rare and may have been missed in MWAS. We conducted another MWAS where we collapsed the non-significant members of cluster 1 into one group (partial cluster 1), leaving *Porphyromonas, Prevotella* and *Corynebacterium_1* as individual genera along with the rest of the genera in MWAS. As expected, we recaptured all 15 PD-associated genera, and in addition, we gained a new significant signal for the partial cluster 1 that was ANCOM and KW significant in both datasets (dataset 1: 2.9-fold increased abundance in PD, ANCOM W=392, KW FDR=0.03; dataset 2: 2.5-fold increased abundance in PD, ANCOM W=480, KW FDR=0.002).

#### Genera in cluster 2

Of the ten PD-associated genera in cluster 2, three (*Oscillospira, Lachnospiraceae_UCG-004* and *Lachnospiraceae_ND3007_group*) have been detected only by sequencing and not yet been cultured. The rest (*Agathobacter, Blautia, Butyricicoccus, Faecalibacterium, Fusicatenibacter, Lachnospira* and *Roseburia*) are all anaerobic, gram positive bacteria in the *Ruminococcaceae*, and *Lachnospiraceae* families. They are best known for producing short chain fatty acids (SCFA), mainly butyrate, which help maintain integrity of the gut membrane, and have anti-inflammatory properties.^47,48^

#### Genera in cluster 3

*Lactobacillus*^49^ and *Bifidobacteria*^50^ are anaerobic gram positive bacteria. They are among ubiquitous inhabitants of the human gastrointestinal microbiome. They metabolize carbohydrates in plants and dairy, and are considered probiotic for their health benefits,^51,52^ although they have also been implicated as cause of infection and excessive immune stimulation in susceptible individuals.^52,53^

### Cause of disease or consequence of medication

Human association studies are powerful tools for identifying disease-relevant leads and to generate hypotheses that can then be tested experimentally. Even if we find a strong candidate that blurs the line between association and causality, we cannot prove that it preceded PD because there are decades of preclinical and prodromal disease, and we do not know when it all begins. While cause cannot be proven in these studies, we can sometimes tease out consequence.

Medications have profound effects on the microbiome.^33^ Levodopa is the most commonly used PD medication (>85% of PD patients were on varying doses of levodopa). To gauge if the association of PD with any of the 15 genera was a consequence of levodopa treatment, we tested if the change in the differential abundance of the 15 genera correlated with increasing levodopa dose.

We found no significant evidence to suggest that the increasing abundance of *Porphyromonas, Prevotella*, or *Corynebacterium_1* (cluster 1) correlated with levodopa therapy. We did find significant evidence in dataset 2 to suggest that increasing doses of levodopa were correlated with decreasing levels of SCFA producing organisms (*Faecalibacterium* P=0.01, *Agathobacter* P=0.02, *Blautia* P=5E-4, *Roseburia* P=0.02, *Fusicatenibacter* P=0.01, *Lachnospira* P=5E-3, *Lachnospiraceae_ND3007_group* P=5E-3, *Lachnospiraceae_UCG-004* P=0.03). A similar pattern was present in dataset 1, albeit most did not reach statistical significance possibly due to the smaller sample size of dataset 1. We also detected significant correlation between increasing levodopa dose and increasing levels of *Bifidobacterium* (dataset 1 P=5E-3, dataset 2 P=2E-6) and *Lactobacillus* (dataset 2 P=4E-3). These data suggest that the increase in abundance of cluster 1 (opportunistic pathogens) is independent of levodopa, but that the reduction in cluster 2 (SCFA) and increase in cluster 3 (probiotics), if not solely a consequence of medication, worsen with increasing doses of levodopa.

## Discussion

### Summary

We confirmed that the gut microbiome is altered in PD and showed that the PD effect on the global composition of the gut microbiome is independent of the effects of sex, age, BMI, constipation, gastrointestinal discomfort, geography, and diet. Using hypothesis-free microbiome-wide association studies we identified 15 PD-associated genera that achieved microbiome-wide significance in both datasets, with two methods, and with or without covariate adjustment. The 15 association signals were robust to the dramatic population-specific differences in the composition of microbiomes of the two datasets. We used hypothesis-free correlation network analysis to infer interactions and to identify communities of co-occurring microorganisms. Using this agnostic approach, we learned that the 15 PD-associated genera represent three polymicrobial clusters. Review of the literature revealed that the clusters, as defined by agnostic network analysis, also share functional characteristics. Our results suggest the gut microbiomes of persons with PD can present with (1) an overabundance of a polymicrobial cluster of opportunistic pathogens, (2) reduced levels of SCFA producing bacteria, and/or (3) elevated levels of carbohydrate metabolizers commonly known as probiotics.

### Alignment with PD literature

Reduced levels of SCFA producing bacteria^12,14,16,18,19,21,26,27^ and elevated levels of probiotic bacteria in PD^14,16,18,21,25-27^ have been reported before, and thus are confirmatory. Overabundance of opportunistic pathogens was a novel finding. We suspect the reason we were able to detect these microorganisms is because they are rare (Figure 2) and we had a much larger sample size and power than prior studies. The microorganisms identified in prior PD studies were among the more abundant microorganisms in the gut. There have been two systematic reviews of PD-microbiome studies, which clearly show the vast disparity in the findings, but also reveal few findings that have emerged in more than one study.^31,32^ The most recent review highlighted 6 associations that were significant in more than one study: *Faecalibacterium, Roseburia, Bifidobacterium, Lactobacillus, Akkemansia* and *Prevotella*.^32^ We confirmed the reduction in *Faecalibacterium* and *Roseburia* (cluster 2), and the increase in *Bifidobacterium* and *Lactobacillus* (cluster 3). We also confirmed increased *Akkermansia* in both datasets but it was only significant in dataset 1. *Prevotella* results are interesting, with Scheperjans et al.^11^ and Petrov et al.^18^ reporting it decreased in PD while we find it elevated in both datasets. The apparent inconsistency may be simply because what is being referred to as “*Prevotella*” is not the same in these studies. We all used different taxonomic classification: Scheperjans et al. reported at the family level (*Prevotellaceae*), we at genus level (*Prevotella*), and Petrov et al. at species level (*Prevotella copri*). The SILVA database we used here, classified family *Prevotellaceae* into 11 genera. The more common genera in the *Prevotellaceae* family (*Paraprevotella, Prevotella_9* and *Prevotella_7*) did in fact have lower frequencies in PD than in controls, as Scheperjans et al. observed, but the difference was not significant in our datasets (FDR>0.6 in both datasets). Species *P. copri*, which Petrov et al. found reduced in PD, was the main species of the *Prevotella_9* genus, which was reduced in our PD samples as well but not significantly (FDR>0.8 in both datasets). We found instead elevated levels of the less common genus *Prevotella* (FDR=0.006 in dataset 1 and FDR=0.02 in dataset 2). These findings suggest family *Prevotellaceae* may be heterogenous in its association with PD. When comparing studies, another important consideration is the reference database: there are many and they have varied phylogenetic resolution and nomenclature. For example, genus *Corynebacterium* in NCBI is divided into 2 non-monophyletic genera in SILVA: *Corynebacterium_1* and *Corynebacterium*. Similarly, what is called genus *Prevotella* in NCBI, is divided into multiple non-monophyletic genera in SILVA (we detected *Prevotella, Prevotella_2, Prevotella_6, Prevotella_7*, and *Prevotella_9*). The varying resolution at which the tests are conducted, and the reference databases used, cause confusion in the literature.

### Opportunistic pathogens

Overabundance of opportunistic pathogens in PD gut microbiome was a novel and potentially the most exciting finding of this study. Braak and colleagues originally hypothesized that non-inherited forms of PD are caused by a pathogen that can pass through the mucosal barrier of the gastrointestinal tract and spread to the brain through the enteric nervous system.^54,55^ While many aspects of Braak’s hypothesis have gained support in recent years, there is no direct evidence that a pathogen is involved. Presence of α-synuclein in the gastrointestinal tract has been documented in persons with established Lewy body disease^56^ as well as those with rapid eye movement sleep behavior disorder which is considered prodromal PD.^57^ Epidemiological studies suggest that truncal vagotomy if conducted decades before onset of PD reduces risk of developing PD.^58,59^ In a mouse model, α-synuclein fibrils injected into the gut induced α-synuclein pathology which spread to the brain resulting in Parkinsonian neurodegeneration and behavioral phenotype; whereas truncal vagotomy and α-synuclein deficiency prevented the gut-to-brain spread and the associated neurodegeneration.^60^ Human studies unrelated to PD have shown that infection in the gut or the olfactory system induce α-synuclein expression, and the increased abundance of α-synuclein mobilizes the immune system to fight the pathogen.^61,62^ It was also shown in a genetic model of PD (*pink1* knock-out mice) that intestinal infection by pathogens elicits activation of cytotoxic T cells in the periphery and the brain and leads to deterioration of dopaminergic cells and motor impairment, suggesting that intestinal infection acts as a triggering event in PD.^63^ Despite the increasing evidence linking the gut, α-synuclein, and inflammation to PD, there is no direct evidence that a pathogen is responsible for the pathology. Here, we present the first evidence from human samples indicating an overabundance of opportunistic pathogens in the gut microbiome of persons with PD. The three genera that rose to significance (*Porphyromonas, Prevotella*, or *Corynebacterium_1)* represented a larger polymicrobial cluster of opportunistic pathogens that co-occur in controls as well as in patients (although at much lower abundances in healthy gut). Per literature, these opportunist pathogens are often harmless, but can grow and cause infections if the immune system is compromised or if they penetrate sterile sites through, for example, compromised membanes.^44-46^ The exciting question is whether these are Braak’s pathogens capable of triggering PD, or they are irrelevant to PD but are able to penetrate the gut and grow because the gut lining is compromised in PD. We re-emphasize that no claims can be made on function based solely on association. The knowledge on the function of microorganisms in the gut is currently limited. While there may be a large body of literature, each organism has been studied with a narrow lens. Organisms that are known to be opportunistic pathogens are being looked for in clinical specimen, whether they have other critical functions is not known. The identity of these microorganisms will enable experimental studies to determine if and how they play a role in PD.

### Anti-inflammatory SCFA producing bacteria

Our second main finding was a polymicrobial cluster of ten genera whose relative abundances were reduced in PD. All ten genera belong to the *Lachnospiraceae* and *Ruminococcaceae* families, well-known for producing SCFA. Several studies had found reduced levels of different SCFA producing bacteria in PD patients.^12,14,16,18,19,21,26,27^ Our finding is therefore confirmatory, and expands on the list of PD-associated genera in these two taxonomic families. We and others noted that the decreasing levels of *Lachnospiraceae* correlate with increasing daily dose of levodopa, disease duration,^12^ disease severity and motor impairment,^26^ which suggest SCFA producing microorganisms diminish as a consequence of medication and/or advancing disease. SCFA promote gastrointestinal motility, maintain integrity of the gut lining, and control inflammation in the gut and the brain,^47,48,64-66^ each of which are compromised in PD. It is important to note, however, that reduced levels of SCFA in the gut has been documented in many inflammatory disorders,^67-71^ and is not specific to PD.

### Probiotics

We also found elevated levels of *Bifidobacterium* and *Lactobacillus* in PD, which have been noted in some of the prior PD studies, albeit not consistently.^14,16,18,21,25-27^ Both are ubiquitous inhabitants of human gut and metabolize carbohydrates derived from plants and dairy.^49,50^ We found a significant correlation between increasing levodopa dose and increasing *Bifidobacterium* and *Lactobacillus* levels. *Lactobacillus* produce a bacterial enzyme that metabolizes levodopa into dopamine before it can reach the brain, reducing efficacy of the drug and requiring higher doses, which in feedback causes further growth of the bacteria.^72,73^ Ironically, *Bifidobacterium* and *Lactobacillus* are sold in stores as probiotics, and a clinical trial has reported fermented milk which contained *Bifidobacterium, Lactobacillus*, and fiber, among other active ingredients, improved constipation in PD.^74^ While generally believed to be safe, and possibly beneficial for the healthy population, they can act as opportunistic pathogens and cause infection and excessive immune stimulation in immune compromised individuals.^52,53^ It is important to understand why *Bifidobacterium* and *Lactobacillus* are elevated in PD and if they are beneficial (a compensatory mechanism to overcome the dysbiosis) or detrimental (feedback of levodopa).

### Conclusion

We uncovered robust and reproducible signals, which reaffirm (SCFA, probiotics) and generate new leads (opportunistic pathogens) for experimentation into cause and effect, disease progression, and therapeutic targets. This study was limited by its singular and precise focus and intentionally conservative analytic execution. There is more to be learned with larger sample sizes with greater power, longitudinal studies to track change from prodromal to advanced disease, and by next generation metagenome sequencing to broaden the scope from bacteria and archaea to include viruses and fungi, and improve the resolution to strain and gene level.

## Methods

### Subjects and Data Collection

#### Subjects

(Supplementary Table 1) The study was approved by institutional review boards for ethical conduct of human subject research at all participating institutions. All subjects provided informed consent for their participation. Subjects were enrolled by NGRC investigators, using standardized methods, at four NGRC affiliated movement disorder clinics in United States. Dataset 1 was collected in Seattle, WA, Albany, NY, and Atlanta, GA in 2014 and included 212 persons with PD and 136 controls.^16^ Dataset 2 was collected in Birmingham, AL during 2015-2018, and included 323 PD and 184 controls (unpublished). PD was diagnosed by a movement disorder specialist using UK Brain Bank criteria,^75^ and controls were self-reported free of neurological disease.

#### Metadata

(Supplementary Table 1) Data were collected using two self-administered questionnaires: an Environmental and Family History Questionnaire (EFQ) and Gut Microbiome Questionnaire (GMQ).^4,16,36^ EFQ covered sex, age, ancestry, and lifetime exposure data on PD-related risk factors. GMQ covered information pertinent to microbiome analysis and was filled out immediately after stool sample collection. PD medications that subjects were taking at the time of sample collection were extracted from medical records by clinical investigators.

#### Stool samples

Subjects collected stool samples at home using DNA/RNA-free sterile swabs (BD BBL CultureSwab Sterile/Media-free Swabs, Fisher Scientific, Pittsburgh, PA). The sample was collected from excreted stool (the kit is not a rectal swab), thus minimizing contamination by skin microbiota. The stool samples were shipped immediately via standard U.S. postal service at ambient temperature and stored at −20°C upon arrival. The collection kit chosen was the most reasonable option at the time (2014). Collection kits with stabilizing solutions (e.g., OMNIgene GUT by DNA Genotek) were first introduced in 2015-2016. Immediate freezing was not feasible because we could not collect stool from over 800 participants, most of whom suffer constipation, while in clinic, nor was it acceptable to the participants to place their stool in their home freezer before shipping. We tested the effect of stool sample travel time on the results as follows. Subjects recorded the collection date and we recorded when it was placed in −20°C freezer, the difference was calculated as the stool sample travel time. We tested the stool sample travel time in cases vs. controls (Supplementary Table 1). We adjusted the PERMANOVA and MWAS for stool sample travel time.

### DNA extraction and sequencing

DNA extraction and sequencing of datasets were done in different laboratories (the Knight Lab at University of California San Diego for dataset 1,^16^ and HudsonAlpha Institute for Biotechnology for dataset 2), keeping methods uniform as possible. Negative controls were included in both datasets. DNA was extracted using MoBio PowerMag Soil DNA Isolation Kit for dataset 1 and MoBio PowerSoil DNA Isolation Kit for dataset 2, both kits using equivalent chemistries (MoBio Industries, Carlsbad, CA).

Hypervariable region 4 (V4) of the bacterial/archaeal 16S rRNA gene was PCR amplified using primers 515F (5’-GTGCCAGCMGCCGCGGTAA-3’) and 806R (5’-GGACTACHVGGGTWTCTAAT-3’) and sequenced using Illumina MiSeq. For dataset 1, paired-end 150 bp was used and all samples were sequenced in one run. For dataset 2, paired-end 250 bp was used and samples were sequenced in 6 runs. Sequence files were de-multiplexed using QIIME2 (core distribution 2018.6)^76^ for dataset 1 and Illumina’s BCL2FASTQ software on BaseSpace for dataset 2. Fifteen samples in dataset 1 had low sequencing counts and were excluded for present analysis.

### Bioinformatics

#### Sequence QC

Forward and reverse primers were trimmed from the 5’ end of sequences using cutadapt v 1.16.^77^ After primer trimming, only sequences with lengths of 147–151 bp in dataset 1 and 230–233 bp in dataset 2 were retained. DADA2 R package v 1.8^78^ was used for the remaining bioinformatics with default parameters unless when specified. Sequences were quality trimmed and filtered using the filterAndTrim function: trimming 3’ ends to 147 bp (forward) and 147 bp (reverse) in dataset 1, and 228 bp (forward) and 203 bp (reverse) in dataset 2, and removing sequences if they exceeded a maximum of two expected errors.

#### Amplicon sequence variant (ASV) inference and ASV tables

For each sequencing run: (a) a model for sequencing error was constructed using the learnErrors function specifying that all bases in all sequences be used for constructing the model, (b) sequences were de-replicated to find unique sequences using the derepFastq function, (c) ASVs were inferred from de-replicated sequences using the dada function, (d) forward and reverse sequences were merged using the mergePairs function, and (e) sequences with <250 bp or >256 bp were removed. This resulted in 1 ASV table for dataset 1 and 6 ASV tables for dataset 2. The 6 ASV tables of dataset 2 were merged using the mergeSequenceTables function. Chimeras were detected and removed using the removeBimeraDenovo function.

#### Data transformation

The following procedures were used to account for variable sequence depth. Sequence counts were normalized to relative abundances (calculated by dividing the number of sequences that were assigned to a unique ASV or to a genus by the total sequence count in the sample) for PERMANOVA when using Canberra or GUniFrac distance, for MWAS when using KW, and for testing correlation with levodopa drug dose. Centered-log ratio (clr) transformation (using the transform function of the microbiome v 1.4.2 R package (http://microbiome.github.com/microbiome)) was used for PCA, and for PERMANOVA when using Aitchison distance. Log ratios (implemented internally in ANCOM and SparCC) were used when using ANCOM for MWAS, and for correlation network analysis. Earlier microbiome studies (including our first study conducted with dataset 1)^16^ often used rarefaction to normalize the sequence count. Although not as efficient as the other methods due to data loss,^79^ for added assurance, we rarefied the data, repeated the MWAS with ANCOM, and were able to recover all 15 significant PD-associated genera.

#### Taxonomic assignment

MWAS and correlation network analysis were conducted at genus level. To define genera, first each unique ASV was assigned to a genus using the assignTaxonomy function, which performs DADA2’s native implementation of the Ribosomal Database Project (RDP) naïve Bayesian classifier,^80^ using SILVA v 132 as reference and a bootstrap confidence of 80%. Then, each genus (including the unclassified genera) was formed by agglomerating all ASVs that were assigned to that genus using the tax_glom function in phyloseq.

Post MWAS, we explored PD-associated genera at the species level. DADA2 pipeline assigns ASVs to species only if the sequences match 100%. We used the addSpecies function in DADA2 with SILVA as reference and addMultiple=TRUE, first finding 100% matches, then filtering out those matches that did not correspond to the genus given by the assignTaxonomy function. To confirm and expand on DADA2-SILVA species assignments, we BLASTed ASVs against the NCBI 16S rRNA gene sequence database (downloaded on 12/3/2019), and extracted taxonomic designations with the most significant E-value. Nucleotide BLAST search was performed using the BLAST+ executables v 2.9.0 with default parameters^81^ (ftp://ftp.ncbi.nlm.nih.gov/blast/executables/blast+/).

#### Phylogenetic trees

A phylogenetic tree of ASVs was constructed for each dataset, as described by Callahan et al.^82^ Briefly, multiple sequence alignment of ASVs was performed using the AlignSeqs function from the DECIPHER R package v 2.8.1.^83^ Aligned ASVs were then used to build a phylogenetic tree using the phangorn R package v 2.5.3.^84^

#### Phyloseq Object

For each dataset, a phyloseq object was created for use in conducting statistical analyses. For each dataset, the ASV table, taxonomic assignments, phylogenetic tree and metadata were merged into a single file, using phyloseq function in phyloseq R package v 1.24.2.^85^

### Data Analysis and Statistics

#### Principal component analysis

PCA was performed on the clr transformed ASV data^35^ using the ordinate function in phyloseq. PC1 and PC2 were plotted using the plot_ordination function in phyloseq (Figure 1).

#### Confounders

We interrogated 47 variables (extracted from metadata) as potential confounders (Supplementary Table 1). In each dataset, we first tested the distribution of each variable in cases vs controls, using Fisher’s exact test (fisher.test function in R) for categorical variables, and Mann-Whitney-*U* (wilcox.test function in R) for quantitative variables. Variables that differed between cases and control at uncorrected P<0.05 were tagged as potential confounders, and were then included in PERMANOVA, along with case-control status, and tested for their effects on microbiome composition (Table 1). Since PERMANOVA was conducted using marginal effects model without rank (see below), simultaneous inclusion of case-control and other variables allowed testing the association of each variable with microbiome composition while adjusting for all other variables in the model. Thus PD effect on microbiome composition (ß diversity) was adjusted for variables that differed between cases and controls. Next, variables that were associated with microbiome composition at PERMANOVA P<0.05 were included as covariates in MWAS. Thus variables that could have led to spurious taxa-disease association because they differed between cases and controls and were also associated with microbiome, were adjusted for in MWAS.

PD medications were present only in PD cases and could not be included as covariates in PERMANOVA or MWAS. To gauge the effect of PD on ß diversity independent of each medication, we performed PERMANOVA using cases not on PD medication vs. controls (Table 1). The potential confounding effect of medication on differential abundance of genera was tested post-MWAS. For each genus whose relative abundance was associated with PD, we tested the correlation between relative abundance of the genus with daily dose of Levodopa (mg/day) using Spearman correlation implemented in the cor.test function in R.

#### Global composition of microbiome (ß diversity)

PERMANOVA was used to identify variables that had a significant effect on ß diversity (Table 1). Tests were conducted using adonis2 function in vegan v 2.5.3 (https://CRAN.R-project.org/package=vegan). P-values were generated by 99,999 permutations which caps at P<1E-5 as highest significance.

Three models were tested.

(Model A) PD vs. control: [Distance ∼ case/control]

(Model B) PD vs. control and all variables tagged as potential confounders:

Dataset 1: [Distance ∼ case/control + sex + age + geography + BMI + loss of 10lbs in past year + gastrointestinal discomfort on day of stool collection + constipation in past three months + alcohol use + fruits or vegetables daily + stool sample travel time]
Dataset 2: [Distance ∼ case/control + sex + age + BMI + loss of 10lbs in past year + gastrointestinal discomfort on day of stool collection + constipation in past three months + alcohol use+ stool sample travel time]

where distance (a measure of (dis)similarity between pairs of samples), age (in years), BMI (kg/m^2^), and stool sample travel time (in days) were continuous variables and the remaining variables were categorical. We tested marginal effects, so that each variable was tested while being adjusted for all others in the model, without priority.

(Model C) Subset of PD cases not on a given PD medication vs controls: [Distance ∼ case/control]

To gauge the effect of the distance measure on the results, all three models were tested using Aitchison,^35^ GUniFrac,^37^ and Canberra^38^ distances. Aitchison distances were calculated by first transforming the ASV data using clr, and then calculating the Euclidean distances using the vegdist function. To calculate GUniFrac distances, unrooted ASV phylogenetic trees were rooted using the root function in the ape v 5.3 R package^86^ specifying the unique ASV with the highest raw count as the root, then data were transformed to relative abundances and distances were calculated using the GUniFrac function in the R package GUniFrac v 1.1,^37^ specifying alpha to be 0.5. To calculate Canberra distances, data were transformed to relative abundances and distances were calculated using the vegdist function in vegan.

#### MWAS

We conducted MWAS to identify the genera whose abundances differed in cases vs. controls. We chose genus classification because it is the highest resolution attainable with high confidence from 16S sequencing.

For statistical analysis of MWAS, we used ANCOM (Table 2, and Supplementary Tables 2-3). We chose ANCOM because it incorporates compositionality of the microbiome data, has low false positive rate, and allows covariate adjustment.^40,42^ ANCOM was run using ANCOM.main function from the ANCOMv2 R code (https://sites.google.com/site/siddharthamandal1985/research). All genera that were detected in each dataset were included in ANCOM MWAS. Sequence counts were transformed to log ratios, as implemented in ANCOM. Case/control status was specified as the main variable. For each dataset, the variables that were significant at P<0.05 in PERMANOVA were included as covariates to be adjusted, as follows:

Dataset 1: [Genus ∼ case/control + sex + age + geography + gastrointestinal discomfort on day of stool collection + fruits or vegetables daily + stool sample travel time]
Dataset 2: [Genus ∼ case/control + sex + age + BMI + constipation in past three months]

where genus (ASV counts assigned to a genus, transformed to log ratios by ANCOM), age (in years), BMI (kg/m^2^), and stool sample travel time (in days) were continuous variables and the remaining variables were categorical. We used the taxa-wise FDR option (multcorr=2) and set significance level to FDR<0.05 to generate W statistics, and threshold of 0.8 for declaring an association as significant.

For comparison, we repeated the MWAS using KW as statistical test (Table 2, and Supplementary Tables 4-5). For KW, genera counts were transformed to genera relative abundances. Unclassified genera, and genera present in <10% of samples were excluded from KW MWAS. KW does not allow covariate adjustment. The kruskal.test function from the stats R package was used to test for significance. P-values were corrected for multiple testing using Benjamini-Hochberg FDR method implemented in the p.adjust function from stats package.

To visualize the distribution of genera that were significant in MWAS (Figure 2), boxplots were created using ggplot2 v 3.1.0 (https://ggplot2.tidyverse.org) with a pseudo-count of 1 added to counts before transforming to relative abundances to avoid taking the log of zero during plotting.

#### Correlation network analysis

(Figure 3, Supplementary Figure 1) For each dataset, and for cases and controls separately, pairwise correlations were calculated between all genera, microbiome-wide, using log-ratio transformed relative abundances as implemented in the SparCC^43^ (https://bitbucket.org/yonatanf/sparcc). Significance of each correlation was determined by pseudo P-values based on 3,000 permutations. Correlation networks were visualized by plotting all genera, microbiome-wide, and connecting correlated genera with an edge, using the program Gephi v 0.9.2.^87^ We chose a minimum correlation (r) of 0.4 to connect two genera with an edge to create the graphic. All correlations r≥0.4 were significant at P<3E-4, which is the maximum significance attainable with 3,000 permutations. To better visualize networks of connected genera, we first used the force-directed algorithm, Force Atlas 2,^88^ then a community detection algorithm^89^ as implemented in Gephi’s modularity function.

## Data availability

Data will be publicly available at NCBI Sequence Read Archive (SRA).

## Code availability

No custom codes were used. All software and packages, their versions, relevant specification and parameters are stated in the “Methods” section.

## Contributions

ZDW and HP were responsible for the design and execution of the study and wrote the first draft of the paper. All co-authors reviewed and critiqued the paper. SAF, EM, CPZ, DGS and MND were responsible for the clinical aspects of the study. ZDW and MA performed bioinformatics and statistical analysis. CLS assisted with blasting and literature searches.

## Acknowledgement

This work was supported by the National Institute of Neurological Disorders and Stroke grant R01036960 (to HP), The U.S. Army Medical Research Materiel Command endorsed by the U.S. Army through the Parkinson’s Research Program Investigator-Initiated Research Award under Award No. W81XWH1810508 (to HP); NIH Udall grant P50 NS062684 (to CPZ) and P50 NS108675 (to DGS), and NIH Training Grant T32NS095775 (to ZDW). Opinions, interpretations, conclusions and recommendations are those of the authors and are not necessarily endorsed by the U.S. Army or the NIH.

