## Supplementary material for "Characterizing dysbiosis of gut microbiome in PD: Evidence for overabundance of opportunistic pathogens"

**Supplementary Table 1. Descriptive Data**

| | | | Dataset 1 | | | Dataset 2 | | |
|---|---|---|---|---|---|---|---|---|
| | | | PD | Control | P | PD | Control | P |
| | Number of subjects | Enrolled with complete data | 212 | 136 | - | 323 | 184 | - |
| | | Passed sequence QC* | 201 | 132 | - | 323 | 184 | - |
| | | Passed sequence and metadata QC* | 199 | 132 | - | 323 | 184 | - |
| | Microbiome | Number of unique ASVs detected | 4,863 | 3,315 | - | 9,188 | 6,667 | - |
| | | Number of genera detected | 404 | 333 | - | 527 | 441 | - |
| 1 | | Stool sample travel time in days, mean (SD) | 3.3 (1.9) | 2.6 (1.5) | 2E-03 | 5.2 (3.3) | 5.0 (2.6) | ns |
| **2** | Age & Sex | **Age, mean (SD)** | 68.3 ( 9.2) | 70.2 (8.6) | 0.04 | 67.7 (9.0) | 66.4 (8.3) | 0.05 |
| **3** | | **Sex (male)** | 67% | 39% | 1E-06 | 64% | 30% | 2E-13 |
| **4** | Geography | **Seattle, WA** | 93 | 58 | - | 0 | 0 | - |
| | | **Albany, NY** | 75 | 62 | - | 0 | 0 | - |
| | | **Atlanta, GA** | 31 | 12 | - | 0 | 0 | - |
| | | **Birmingham, AL** | 0 | 0 | - | 323 | 184 | - |
| 5 | Ancestry | Race (%White) | 99% | 100% | ns | 99% | >99% | ns |
| 6 | | Jewish | - | - | - | 7% | 7% | ns |
| **7** | Weight | **BMI, mean (SD)** | 26.6 (5.5) | 28.3 (5.7) | 0.02 | 27.4 (5.0) | 27.9 (5.9) | ns |
| **8** | | **Lost >10 pounds in past year** | 23% | 12% | 0.01 | 25% | 12% | 3E-04 |
| 9 | | Gained >10 pounds in past year | 13% | 8% | ns | 15% | 11% | ns |
| **10** | Daily Diet | **Fruits or vegetables** | 78% | 89% | 0.02 | - | - | - |
| 11 | | Meat, fish, poultry | 57% | 63% | ns | - | - | - |
| 12 | | Nuts | 22% | 28% | ns | - | - | - |
| 13 | | Yogurt | 36% | 45% | ns | - | - | - |
| 14 | | Grains | 69% | 67% | ns | - | - | - |
| **15** | | **Alcohol** | 60% | 71% | 0.04 | 42% | 56% | 3E-03 |
| 16 | | Tobacco | 7% | 4% | ns | 4% | 7% | ns |
| 17 | | Caffeine | 71% | 76% | ns | 86% | 88% | ns |

| | | | | | | | | |
|---|---|---|---|---|---|---|---|---|
| 18 | GI Health | Constipation in ≥3 days prior to stool collection | 15% | 2% | 3E-05 | 18% | 5% | 2E-05 |
| 19 | | Diarrhea on the day of stool collection | 3% | 2% | ns | 4% | 3% | ns |
| 20 | | GI pain on the day of stool collection | 9% | 7% | ns | 9% | 2% | 1E-03 |
| 21 | | Gas on the day of stool collection | 14% | 2% | 9E-05 | 16% | 4% | 2E-04 |
| 22 | | Bloating on the day of stool collection | 10% | 2% | 7E-03 | 12% | 5% | 0.01 |
| **23** | | **GI discomfort on the day of stool collection (yes to any item 18-22)** | 57% | 22% | 2E-09 | 34% | 15% | 4E-06 |
| **24** | | **Constipation in the past 3 months** | 43% | 5% | 6E-16 | 44% | 17% | 6E-10 |
| 25 | | Diarrhea in the past 3 months | 17% | 22% | ns | 26% | 30% | ns |
| 26 | | Colitis | 5% | 2% | ns | 17% | 13% | ns |
| 27 | | IBS | 7% | 6% | ns | 5% | 8% | ns |
| 28 | | Crohn's disease | 2% | 1% | ns | 1% | 0 | ns |
| 29 | | IBD | 3% | 2% | ns | 3% | 2% | ns |
| 30 | | Ulcers | 9% | 7% | ns | 2% | 2% | ns |
| 31 | | SIBO | - | - | - | 0 | 0 | ns |
| 32 | | Celiac | - | - | - | 0 | 0 | ns |
| 33 | | GI cancer | - | - | - | <1% | <1% | ns |
| 34 | | Intestinal disease (yes to any item 26-33) | 20% | 15% | ns | 27% | 24% | ns |
| 35 | Medications | Currently taking digestive medication | 31% | 17% | 6E-03 | - | - | - |
| 36 | | Currently taking antibiotics | 4% | 2% | ns | 4% | 4% | ns |
| 37 | | Taken antibiotics in past 3 months | 13% | 17% | ns | 21% | 19% | ns |
| 38 | | Currently taking anti-inflammatory drugs | 41% | 44% | ns | - | - | - |
| 39 | | Currently taking probiotics | 23% | 26% | ns | - | - | - |
| 40 | Parkinson Medications | Patients on carbidopa/levodopa | 91% | - | - | 85% | - | - |
| 41 | | Levodopa dose, mean (SD) | 764 (574) | - | - | 563 (443) | - | - |
| 42 | | Patients on dopamine agonist | 53% | - | - | 51% | - | - |
| 43 | | Patients on MAO-B inhibitor | 38% | - | - | 27% | - | - |
| 44 | | Patients on amantadine | 26% | - | - | 19% | - | - |
| 45 | | Patients on COMT inhibitor | 20% | - | - | 4% | - | - |

| | | | | | | | |
|---|---|---|---|---|---|---|---|
| 46 | Patients on anticholinergics | 4% | - | - | 3% | - | - |
| 47 | Patients not on PD medication | 2% | - | - | 5% | - | - |

*15 samples were excluded from dataset 1 due to low sequence count, and additional 2 were excluded due to unreliable metadata.

Variables that were carried forward and adjusted as covariates in PERMANOVA for testing β diversity and ANCOM for MWAS are shown in bold. Constipation in ≥3 days prior to stool collection, GI pain on day of stool collection, gas on day of stool collection, and bloating on day of stool collection were captured by GI discomfort on day of stool collection. Currently taking digestive medication was not selected as covariate because it was no longer significant when adjusted for GI discomfort.

**Supplementary Table 2. MWAS of dataset 1 conducted using ANCOM**

W= ANCOM score indicating the number of times a genus achieved FDR<0.05 as compared to other genera (maximum W possible: 444 in dataset 1, 560 in dataset 2).

0.8= Threshold at which results were considered significant (TRUE).

| W | 0.8 | Kingdom | Phylum | Class | Order | Family | Genus |
|---|---|---|---|---|---|---|---|
| 441 | TRUE | Bacteria | Firmicutes | Clostridia | Clostridiales | Lachnospiraceae | Agathobacter |
| 426 | TRUE | Bacteria | Firmicutes | Clostridia | Clostridiales | Lachnospiraceae | Lachnospira |
| 418 | TRUE | Bacteria | Firmicutes | Clostridia | Clostridiales | Lachnospiraceae | Lachnospiraceae_ND3007_group |
| 411 | TRUE | Bacteria | Firmicutes | Clostridia | Clostridiales | Ruminococcaceae | Faecalibacterium |
| 410 | TRUE | Bacteria | Actinobacteria | Actinobacteria | Bifidobacteriales | Bifidobacteriaceae | Bifidobacterium |
| 410 | TRUE | Bacteria | Firmicutes | Clostridia | Clostridiales | Lachnospiraceae | Blautia |
| 407 | TRUE | Bacteria | Firmicutes | Bacilli | Lactobacillales | Lactobacillaceae | Lactobacillus |
| 406 | TRUE | Bacteria | Bacteroidetes | Bacteroidia | Bacteroidales | Porphyromonadaceae | Porphyromonas |
| 400 | TRUE | Bacteria | Bacteroidetes | Bacteroidia | Bacteroidales | Prevotellaceae | Prevotella |
| 393 | TRUE | Bacteria | Firmicutes | Clostridia | Clostridiales | Lachnospiraceae | Hungatella |
| 391 | TRUE | Bacteria | Firmicutes | Clostridia | Clostridiales | Lachnospiraceae | Roseburia |
| 388 | TRUE | Bacteria | Firmicutes | Clostridia | Clostridiales | Lachnospiraceae | Fusicatenibacter |
| 384 | TRUE | Bacteria | Firmicutes | Clostridia | Clostridiales | Lachnospiraceae | Lachnospiraceae_UCG-004 |
| 382 | TRUE | Bacteria | Firmicutes | Clostridia | Clostridiales | Ruminococcaceae | Butyricicoccus |
| 378 | TRUE | Bacteria | Firmicutes | Clostridia | Clostridiales | Family_XI | Ezakiella |
| 376 | TRUE | Bacteria | Synergistetes | Synergistia | Synergistales | Synergistaceae | Cloacibacillus |
| 374 | TRUE | Bacteria | Firmicutes | Negativicutes | Selenomonadales | Veillonellaceae | Megasphaera |
| 372 | TRUE | Bacteria | Firmicutes | Clostridia | Clostridiales | Lachnospiraceae | Coprococcus_3 |
| 368 | TRUE | Bacteria | Firmicutes | Erysipelotrichia | Erysipelotrichales | Erysipelotrichaceae | Coprobacillus |
| 367 | TRUE | Bacteria | Firmicutes | Clostridia | Clostridiales | Ruminococcaceae | Oscillospira |
| 365 | TRUE | Bacteria | Verrucomicrobia | Verrucomicrobiae | Verrucomicrobiales | Akkermansiaceae | Akkermansia |
| 360 | TRUE | Bacteria | Actinobacteria | Actinobacteria | Corynebacteriales | Corynebacteriaceae | Corynebacterium_1 |
| 356 | TRUE | Bacteria | Proteobacteria | Gammaproteobacteria | Pasteurellales | Pasteurellaceae | Haemophilus |
| 347 | FALSE | Bacteria | Firmicutes | Clostridia | Clostridiales | Lachnospiraceae | Anaerostipes |
| 331 | FALSE | Bacteria | Firmicutes | Erysipelotrichia | Erysipelotrichales | Erysipelotrichaceae | NA |
| 327 | FALSE | Archaea | Euryarchaeota | Methanobacteria | Methanobacteriales | Methanobacteriaceae | Methanobrevibacter |
| 326 | FALSE | Bacteria | Firmicutes | Clostridia | Clostridiales | Ruminococcaceae | UBA1819 |
| 323 | FALSE | Bacteria | Firmicutes | Clostridia | Clostridiales | Ruminococcaceae | Ruminococcaceae_UCG-013 |
| 319 | FALSE | Bacteria | Firmicutes | Clostridia | Clostridiales | Family_XI | Anaerococcus |
| 318 | FALSE | Bacteria | Firmicutes | Clostridia | Clostridiales | Ruminococcaceae | Ruminococcaceae_UCG-004 |
| 306 | FALSE | Bacteria | Firmicutes | Clostridia | Clostridiales | Ruminococcaceae | Anaerotruncus |
| 302 | FALSE | Bacteria | Actinobacteria | Actinobacteria | Actinomycetales | Actinomycetaceae | Varibaculum |

| | | | | | | | |
|---|---|---|---|---|---|---|---|
| 293 | FALSE | Bacteria | Firmicutes | Clostridia | Clostridiales | Ruminococcaceae | NA |
| 275 | FALSE | Bacteria | Actinobacteria | Actinobacteria | Actinomycetales | Actinomycetaceae | Mobiluncus |
| 263 | FALSE | Bacteria | Firmicutes | Clostridia | Clostridiales | NA | NA |
| 252 | FALSE | Bacteria | Firmicutes | Clostridia | Clostridiales | Lachnospiraceae | Lachnospiraceae_NK4B4_group |
| 249 | FALSE | Bacteria | Firmicutes | Clostridia | Clostridiales | Family_XI | Peptoniphilus |
| 65 | FALSE | Bacteria | Actinobacteria | Actinobacteria | Bifidobacteriales | Bifidobacteriaceae | NA |
| 31 | FALSE | Bacteria | Tenericutes | Mollicutes | Anaeroplasmatales | Anaeroplasmataceae | Anaeroplasma |
| 20 | FALSE | NA | NA | NA | NA | NA | NA |
| 19 | FALSE | Bacteria | Firmicutes | Clostridia | Clostridiales | Ruminococcaceae | Fournierella |
| 17 | FALSE | Bacteria | Proteobacteria | Gammaproteobacteria | Betaproteobacteriales | Burkholderiaceae | Alcaligenes |
| 16 | FALSE | Bacteria | Bacteroidetes | Bacteroidia | Bacteroidales | Prevotellaceae | Prevotellaceae_UCG-001 |
| 15 | FALSE | Bacteria | Bacteroidetes | Bacteroidia | Bacteroidales | Prevotellaceae | Prevotella_6 |
| 15 | FALSE | Bacteria | Bacteroidetes | Bacteroidia | Bacteroidales | NA | NA |
| 15 | FALSE | Bacteria | Firmicutes | Bacilli | Lactobacillales | Aerococcaceae | NA |
| 14 | FALSE | Bacteria | Firmicutes | Clostridia | Clostridiales | Lachnospiraceae | NA |
| 14 | FALSE | Bacteria | Proteobacteria | Deltaproteobacteria | Desulfovibrionales | Desulfovibrionaceae | Bilophila |
| 14 | FALSE | Bacteria | Firmicutes | Clostridia | Clostridiales | Peptostreptococcaceae | Peptoclostridium |
| 14 | FALSE | Bacteria | Actinobacteria | Coriobacteriia | Coriobacteriales | Eggerthellaceae | CHKCI002 |
| 14 | FALSE | Bacteria | Proteobacteria | Gammaproteobacteria | Betaproteobacteriales | Burkholderiaceae | Paenalcaligenes |
| 14 | FALSE | Bacteria | Bacteroidetes | Bacteroidia | Sphingobacteriales | env.OPS_17 | NA |
| 14 | FALSE | Bacteria | Proteobacteria | Gammaproteobacteria | Enterobacteriales | Enterobacteriaceae | Pseudocitrobacter |
| 14 | FALSE | Bacteria | Firmicutes | Bacilli | Lactobacillales | Leuconostocaceae | Leuconostoc |
| 14 | FALSE | Bacteria | Actinobacteria | Actinobacteria | Propionibacteriales | Nocardioidaceae | Nocardioides |
| 14 | FALSE | Bacteria | Firmicutes | Clostridia | Clostridiales | Ruminococcaceae | Ruminiclostridium |
| 14 | FALSE | Bacteria | Proteobacteria | Gammaproteobacteria | Betaproteobacteriales | Burkholderiaceae | Pelomonas |
| 14 | FALSE | Bacteria | Proteobacteria | Gammaproteobacteria | Pseudomonadales | Moraxellaceae | Enhydrobacter |
| 14 | FALSE | Bacteria | Firmicutes | Erysipelotrichia | Erysipelotrichales | Erysipelotrichaceae | Erysipelothrix |
| 14 | FALSE | Bacteria | Dependentiae | Babeliae | Babeliales | Vermiphilaceae | NA |
| 14 | FALSE | Bacteria | Actinobacteria | Actinobacteria | Micrococcales | Intrasporangiaceae | Ornithinimicrobium |
| 14 | FALSE | Bacteria | Firmicutes | Bacilli | Lactobacillales | Enterococcaceae | Melissococcus |
| 14 | FALSE | Bacteria | Proteobacteria | Gammaproteobacteria | Betaproteobacteriales | Methylophilaceae | Methylobacillus |
| 14 | FALSE | Bacteria | Actinobacteria | Actinobacteria | Corynebacteriales | Nocardiaceae | Rhodococcus |
| 14 | FALSE | Bacteria | Firmicutes | Clostridia | Clostridiales | Eubacteriaceae | Anaerofustis |
| 14 | FALSE | Bacteria | Bacteroidetes | Bacteroidia | Bacteroidales | Tannerellaceae | NA |
| 13 | FALSE | Bacteria | Bacteroidetes | Bacteroidia | Bacteroidales | Rikenellaceae | Alistipes |
| 13 | FALSE | Bacteria | Firmicutes | Clostridia | Clostridiales | Lachnospiraceae | Sellimonas |
| 13 | FALSE | Bacteria | Bacteroidetes | Bacteroidia | Flavobacteriales | Crocinitomicaceae | NA |
| 13 | FALSE | Bacteria | Proteobacteria | Gammaproteobacteria | Enterobacteriales | Enterobacteriaceae | Yersinia |
| 13 | FALSE | Bacteria | Actinobacteria | Actinobacteria | Streptosporangiales | Nocardiopsaceae | Nocardiopsis |
| 13 | FALSE | Bacteria | Proteobacteria | Gammaproteobacteria | Enterobacteriales | Enterobacteriaceae | Hafnia-Obesumbacterium |
| 13 | FALSE | Bacteria | Actinobacteria | Actinobacteria | Micrococcales | Micrococcaceae | Glutamicibacter |

| | | | | | | | |
|---|---|---|---|---|---|---|---|
| 13 | FALSE | Bacteria | Firmicutes | Clostridia | Clostridiales | Family_XI | Parvimonas |
| 13 | FALSE | Bacteria | Proteobacteria | Gammaproteobacteria | Betaproteobacteriales | Burkholderiaceae | Xylophilus |
| 13 | FALSE | Bacteria | Actinobacteria | Actinobacteria | Micrococcales | Microbacteriaceae | Pseudoclavibacter |
| 13 | FALSE | Bacteria | Bacteroidetes | Bacteroidia | Bacteroidales | Prevotellaceae | Prevotellaceae_Ga6A1_group |
| 13 | FALSE | Bacteria | Proteobacteria | Alphaproteobacteria | Rhizobiales | Xanthobacteraceae | Bradyrhizobium |
| 13 | FALSE | Bacteria | Actinobacteria | Actinobacteria | Micrococcales | Microbacteriaceae | Leucobacter |
| 13 | FALSE | Bacteria | Actinobacteria | Actinobacteria | Pseudonocardiales | Pseudonocardiaceae | Pseudonocardia |
| 13 | FALSE | Bacteria | Actinobacteria | Actinobacteria | NA | NA | NA |
| 13 | FALSE | Bacteria | Actinobacteria | Actinobacteria | Micrococcales | Micrococcaceae | Paenarthrobacter |
| 13 | FALSE | Bacteria | Bacteroidetes | Bacteroidia | Bacteroidales | Marinifilaceae | NA |
| 13 | FALSE | Bacteria | Actinobacteria | Actinobacteria | Frankiales | Geodermatophilaceae | Blastococcus |
| 13 | FALSE | Bacteria | Firmicutes | Bacilli | Bacillales | Family_X | Thermicanus |
| 13 | FALSE | Bacteria | Bacteroidetes | Bacteroidia | Flavobacteriales | Weeksellaceae | Empedobacter |
| 13 | FALSE | Eukaryota | NA | NA | NA | NA | NA |
| 13 | FALSE | Bacteria | Firmicutes | Bacilli | Lactobacillales | Enterococcaceae | NA |
| 13 | FALSE | Bacteria | Proteobacteria | Deltaproteobacteria | NA | NA | NA |
| 13 | FALSE | Bacteria | Entotheonellaeota | Entotheonellia | Entotheonellales | Entotheonellaceae | NA |
| 13 | FALSE | Bacteria | Proteobacteria | Gammaproteobacteria | Pseudomonadales | Moraxellaceae | NA |
| 13 | FALSE | Bacteria | Cyanobacteria | Melainabacteria | Obscuribacterales | NA | NA |
| 12 | FALSE | Bacteria | Firmicutes | Clostridia | NA | NA | NA |
| 12 | FALSE | Bacteria | Epsilonbacteraeota | Campylobacteria | Campylobacterales | Campylobacteraceae | Campylobacter |
| 12 | FALSE | Bacteria | Proteobacteria | Gammaproteobacteria | Betaproteobacteriales | Burkholderiaceae | Comamonas |
| 12 | FALSE | Bacteria | Firmicutes | Bacilli | Bacillales | Paenibacillaceae | Paenibacillus |
| 12 | FALSE | Bacteria | Proteobacteria | Alphaproteobacteria | Paracaedibacterales | Paracaedibacteraceae | Candidatus_Odyssella |
| 12 | FALSE | Bacteria | Actinobacteria | Actinobacteria | Corynebacteriales | Corynebacteriaceae | Lawsonella |
| 12 | FALSE | Bacteria | Bacteroidetes | Bacteroidia | Sphingobacteriales | Sphingobacteriaceae | NA |
| 12 | FALSE | Bacteria | Firmicutes | Bacilli | Bacillales | Bacillaceae | NA |
| 12 | FALSE | Bacteria | Firmicutes | Erysipelotrichia | Erysipelotrichales | Erysipelotrichaceae | Catenisphaera |
| 12 | FALSE | Bacteria | Proteobacteria | Gammaproteobacteria | Betaproteobacteriales | Rhodocyclaceae | NA |
| 12 | FALSE | Bacteria | Proteobacteria | Gammaproteobacteria | Betaproteobacteriales | Methylophilaceae | Methylophilus |
| 12 | FALSE | Bacteria | Proteobacteria | Gammaproteobacteria | Betaproteobacteriales | Burkholderiaceae | Herminiimonas |
| 12 | FALSE | Bacteria | Firmicutes | Clostridia | Clostridiales | Ruminococcaceae | Phocea |
| 12 | FALSE | Bacteria | Firmicutes | Clostridia | Clostridiales | Family_XIII | Family_XIII_UCG-001 |
| 12 | FALSE | Bacteria | Bacteroidetes | Bacteroidia | Sphingobacteriales | Sphingobacteriaceae | Pedobacter |
| 12 | FALSE | Bacteria | Firmicutes | Clostridia | Clostridiales | Lachnospiraceae | Moryella |
| 12 | FALSE | Bacteria | Proteobacteria | Alphaproteobacteria | Sphingomonadales | Sphingomonadaceae | Altererythrobacter |
| 12 | FALSE | Bacteria | Firmicutes | Negativicutes | Selenomonadales | Veillonellaceae | Mitsuokella |
| 12 | FALSE | Bacteria | Fusobacteria | Fusobacteriia | Fusobacteriales | Fusobacteriaceae | Cetobacterium |
| 12 | FALSE | Bacteria | Proteobacteria | Deltaproteobacteria | Bdellovibrionales | Bdellovibrionaceae | Bdellovibrio |
| 12 | FALSE | Bacteria | Firmicutes | Clostridia | Clostridiales | Lachnospiraceae | Lachnospiraceae_FCS020_group |
| 12 | FALSE | Bacteria | Proteobacteria | Alphaproteobacteria | Rhizobiales | Rhizobiaceae | Allorhizobium-Neorhizobium-Pararhizobium-Rhizobium |

| | | | | | | | |
|---|---|---|---|---|---|---|---|
| 12 | FALSE | Bacteria | Proteobacteria | Deltaproteobacteria | Bdellovibrionales | Bacteriovoracaceae | Peredibacter |
| 12 | FALSE | Bacteria | Firmicutes | Clostridia | Clostridiales | Ruminococcaceae | Ruminococcaceae_UCG-008 |
| 12 | FALSE | Bacteria | Bacteroidetes | Bacteroidia | Bacteroidales | Prevotellaceae | Prevotellaceae_UCG-003 |
| 12 | FALSE | Bacteria | Actinobacteria | Actinobacteria | Actinomycetales | Actinomycetaceae | Actinotignum |
| 12 | FALSE | Bacteria | Proteobacteria | Alphaproteobacteria | Rhizobiales | Rhizobiaceae | Ochrobactrum |
| 12 | FALSE | Bacteria | Firmicutes | Bacilli | Lactobacillales | Aerococcaceae | Facklamia |
| 12 | FALSE | Bacteria | Proteobacteria | Gammaproteobacteria | Salinisphaerales | Solimonadaceae | Nevskia |
| 12 | FALSE | Bacteria | Proteobacteria | Gammaproteobacteria | Betaproteobacteriales | Burkholderiaceae | Simplicispira |
| 12 | FALSE | Bacteria | Firmicutes | Clostridia | Clostridiales | Peptostreptococcaceae | Peptostreptococcus |
| 12 | FALSE | Bacteria | Actinobacteria | Coriobacteriia | Coriobacteriales | Eggerthellaceae | Enterorhabdus |
| 12 | FALSE | Bacteria | Bacteroidetes | Bacteroidia | Bacteroidales | Rikenellaceae | Rikenella |
| 12 | FALSE | Bacteria | Proteobacteria | Alphaproteobacteria | Caulobacterales | Caulobacteraceae | NA |
| 12 | FALSE | Bacteria | Tenericutes | Mollicutes | NA | NA | NA |
| 12 | FALSE | Bacteria | Lentisphaerae | Oligosphaeria | Oligosphaerales | Oligosphaeraceae | Z20 |
| 12 | FALSE | Bacteria | Firmicutes | Clostridia | Clostridiales | Lachnospiraceae | Herbinix |
| 12 | FALSE | Bacteria | Bacteroidetes | Bacteroidia | Flavobacteriales | Weeksellaceae | Elizabethkingia |
| 12 | FALSE | Bacteria | Firmicutes | Clostridia | Clostridiales | Family_XI | Helcococcus |
| 12 | FALSE | Bacteria | Actinobacteria | Actinobacteria | Actinomycetales | Actinomycetaceae | NA |
| 12 | FALSE | Bacteria | Proteobacteria | Gammaproteobacteria | Betaproteobacteriales | Rhodocyclaceae | Methyloversatilis |
| 12 | FALSE | Bacteria | Firmicutes | Clostridia | Clostridiales | Heliobacteriaceae | Hydrogenispora |
| 12 | FALSE | Bacteria | Cyanobacteria | Oxyphotobacteria | Chloroplast | NA | NA |
| 12 | FALSE | Bacteria | Proteobacteria | Gammaproteobacteria | Betaproteobacteriales | Burkholderiaceae | Verticia |
| 12 | FALSE | Bacteria | Spirochaetes | Brachyspirae | Brachyspirales | Brachyspiraceae | Brachyspira |
| 12 | FALSE | Bacteria | Firmicutes | Negativicutes | Selenomonadales | Veillonellaceae | Anaerovibrio |
| 12 | FALSE | Bacteria | Proteobacteria | Gammaproteobacteria | Betaproteobacteriales | Burkholderiaceae | Bordetella |
| 12 | FALSE | Bacteria | Firmicutes | Clostridia | Clostridiales | Lachnospiraceae | Robinsoniella |
| 12 | FALSE | Bacteria | Actinobacteria | Actinobacteria | Micrococcales | Micrococcaceae | NA |
| 12 | FALSE | Bacteria | Actinobacteria | Actinobacteria | Actinomycetales | Actinomycetaceae | Arcanobacterium |
| 12 | FALSE | Bacteria | Proteobacteria | Alphaproteobacteria | Rhizobiales | Rhizobiaceae | Brucella |
| 12 | FALSE | Bacteria | Actinobacteria | Actinobacteria | Micrococcales | Micrococcaceae | Nesterenkonia |
| 12 | FALSE | Bacteria | Firmicutes | Clostridia | Clostridiales | Lachnospiraceae | GCA-900066755 |
| 12 | FALSE | Bacteria | Proteobacteria | Gammaproteobacteria | JTB23 | NA | NA |
| 12 | FALSE | Bacteria | Firmicutes | Clostridia | Clostridiales | Family_XIII | Anaerovorax |
| 12 | FALSE | Bacteria | Kiritimatiellaeota | Kiritimatiellae | WCHB1-41 | NA | NA |
| 12 | FALSE | Bacteria | Bacteroidetes | Bacteroidia | Cytophagales | Spirosomaceae | Dyadobacter |
| 12 | FALSE | Bacteria | Bacteroidetes | Rhodothermia | Rhodothermales | Rhodothermaceae | NA |
| 12 | FALSE | Bacteria | Actinobacteria | Actinobacteria | Micrococcales | Microbacteriaceae | Amnibacterium |
| 12 | FALSE | Bacteria | Firmicutes | Bacilli | Lactobacillales | Carnobacteriaceae | Granulicatella |
| 12 | FALSE | Bacteria | Firmicutes | Bacilli | Lactobacillales | NA | NA |
| 12 | FALSE | Bacteria | Actinobacteria | Actinobacteria | Actinomycetales | Actinomycetaceae | Trueperella |
| 12 | FALSE | Bacteria | Actinobacteria | Actinobacteria | Micrococcales | Promicromonosporaceae | Cellulosimicrobium |

| | | | | | | | |
|---|---|---|---|---|---|---|---|
| 12 | FALSE | Bacteria | Firmicutes | Clostridia | Clostridiales | Ruminococcaceae | Harryflintia |
| 12 | FALSE | Bacteria | Patescibacteria | Saccharimonadia | Saccharimonadales | Saccharimonadaceae | NA |
| 12 | FALSE | Bacteria | Proteobacteria | Gammaproteobacteria | Betaproteobacteriales | Neisseriaceae | Neisseria |
| 12 | FALSE | Bacteria | Firmicutes | Clostridia | Clostridiales | Lachnospiraceae | Epulopiscium |
| 12 | FALSE | Bacteria | Bacteroidetes | Bacteroidia | Flavobacteriales | Weeksellaceae | NA |
| 12 | FALSE | Bacteria | Actinobacteria | Actinobacteria | Micrococcales | Micrococcaceae | Micrococcus |
| 12 | FALSE | Bacteria | Firmicutes | Clostridia | Clostridiales | Peptostreptococcaceae | Clostridioides |
| 12 | FALSE | Bacteria | Firmicutes | Clostridia | Clostridiales | Lachnospiraceae | Cuneatibacter |
| 12 | FALSE | Bacteria | Proteobacteria | Gammaproteobacteria | Pasteurellales | Pasteurellaceae | NA |
| 12 | FALSE | Bacteria | Firmicutes | Clostridia | Clostridiales | Ruminococcaceae | Acetanaerobacterium |
| 12 | FALSE | Bacteria | Proteobacteria | Alphaproteobacteria | Sphingomonadales | Sphingomonadaceae | NA |
| 12 | FALSE | Bacteria | Proteobacteria | Alphaproteobacteria | Rhizobiales | Rhizobiaceae | Paenochrobactrum |
| 12 | FALSE | Bacteria | Proteobacteria | Alphaproteobacteria | Rhizobiales | Rhizobiaceae | Shinella |
| 12 | FALSE | Bacteria | Actinobacteria | Coriobacteriia | Coriobacteriales | Atopobiaceae | NA |
| 12 | FALSE | Bacteria | Firmicutes | Clostridia | Clostridiales | Ruminococcaceae | Candidatus_Soleaferrea |
| 12 | FALSE | Bacteria | Bacteroidetes | Bacteroidia | Chitinophagales | Saprospiraceae | NA |
| 12 | FALSE | Bacteria | Proteobacteria | Gammaproteobacteria | Betaproteobacteriales | Burkholderiaceae | Parapusillimonas |
| 12 | FALSE | Bacteria | Firmicutes | Negativicutes | Selenomonadales | Acidaminococcaceae | Succiniclasticum |
| 12 | FALSE | Bacteria | Verrucomicrobia | Verrucomicrobiae | Opitutales | Puniceicoccaceae | NA |
| 12 | FALSE | Bacteria | Proteobacteria | Gammaproteobacteria | Betaproteobacteriales | Burkholderiaceae | Ralstonia |
| 12 | FALSE | Bacteria | Firmicutes | Bacilli | Bacillales | Paenibacillaceae | Ammoniphilus |
| 12 | FALSE | Bacteria | Proteobacteria | Gammaproteobacteria | Betaproteobacteriales | NA | NA |
| 12 | FALSE | Bacteria | Proteobacteria | Gammaproteobacteria | Xanthomonadales | Xanthomonadaceae | Thermomonas |
| 12 | FALSE | Bacteria | Firmicutes | Clostridia | Clostridiales | Peptococcaceae | Desulfotomaculum |
| 12 | FALSE | Archaea | Euryarchaeota | Methanobacteria | Methanobacteriales | Methanobacteriaceae | Methanobacterium |
| 12 | FALSE | Bacteria | Actinobacteria | Actinobacteria | Micrococcales | Dermabacteraceae | Dermabacter |
| 12 | FALSE | Bacteria | Fusobacteria | Fusobacteriia | Fusobacteriales | Fusobacteriaceae | NA |
| 12 | FALSE | Bacteria | Firmicutes | Bacilli | Bacillales | Staphylococcaceae | Jeotgalicoccus |
| 12 | FALSE | Bacteria | Planctomycetes | Planctomycetacia | Pirellulales | Pirellulaceae | Rhodopirellula |
| 12 | FALSE | Bacteria | Proteobacteria | Alphaproteobacteria | Rhizobiales | Rhizobiaceae | Neorhizobium |
| 12 | FALSE | Bacteria | Firmicutes | Clostridia | Clostridiales | Family_XI | NA |
| 12 | FALSE | Bacteria | Proteobacteria | Alphaproteobacteria | Rhizobiales | Beijerinckiaceae | Methylobacterium |
| 12 | FALSE | Bacteria | Firmicutes | Clostridia | Clostridiales | Peptostreptococcaceae | Paeniclostridium |
| 12 | FALSE | Bacteria | Proteobacteria | Gammaproteobacteria | Aeromonadales | Aeromonadaceae | Tolumonas |
| 12 | FALSE | Bacteria | Chloroflexi | Dehalococcoidia | SAR202_clade | NA | NA |
| 12 | FALSE | Bacteria | Firmicutes | Bacilli | Bacillales | Bacillaceae | Oceanobacillus |
| 12 | FALSE | Bacteria | Bacteroidetes | Bacteroidia | Bacteroidales | Dysgonomonadaceae | Proteiniphilum |
| 12 | FALSE | Bacteria | Firmicutes | Clostridia | Clostridiales | Eubacteriaceae | Pseudoramibacter |
| 12 | FALSE | Bacteria | Proteobacteria | Alphaproteobacteria | Rhizobiales | Xanthobacteraceae | NA |
| 12 | FALSE | Bacteria | Firmicutes | Bacilli | Lactobacillales | Aerococcaceae | Aerococcus |
| 12 | FALSE | Bacteria | Firmicutes | Bacilli | NA | NA | NA |

| | | | | | | | |
|---|---|---|---|---|---|---|---|
| 12 | FALSE | Bacteria | Bacteroidetes | NA | NA | NA | NA |
| 12 | FALSE | Bacteria | Proteobacteria | Gammaproteobacteria | Xanthomonadales | Rhodanobacteraceae | Rhodanobacter |
| 12 | FALSE | Bacteria | Bacteroidetes | Bacteroidia | Flavobacteriales | Weeksellaceae | Moheibacter |
| 12 | FALSE | Bacteria | Proteobacteria | Alphaproteobacteria | Rhizobiales | NA | NA |
| 12 | FALSE | Bacteria | Firmicutes | Negativicutes | Selenomonadales | Veillonellaceae | Selenomonas_4 |
| 12 | FALSE | Bacteria | Proteobacteria | Alphaproteobacteria | NA | NA | NA |
| 12 | FALSE | Bacteria | Firmicutes | Clostridia | Clostridiales | Defluviitaleaceae | NA |
| 12 | FALSE | Bacteria | Bacteroidetes | Bacteroidia | Flavobacteriales | Flavobacteriaceae | NA |
| 12 | FALSE | Bacteria | Proteobacteria | Deltaproteobacteria | Myxococcales | mle1-27 | NA |
| 12 | FALSE | Bacteria | Firmicutes | Clostridia | Clostridiales | Syntrophomonadaceae | NA |
| 12 | FALSE | Bacteria | Actinobacteria | Actinobacteria | Actinomycetales | Actinomycetaceae | Actinobaculum |
| 12 | FALSE | Bacteria | Bacteroidetes | Bacteroidia | Cytophagales | Spirosomaceae | Rhabdobacter |
| 12 | FALSE | Bacteria | Actinobacteria | NA | NA | NA | NA |
| 12 | FALSE | Bacteria | Firmicutes | Clostridia | Clostridiales | Ruminococcaceae | Pseudoflavonifractor |
| 12 | FALSE | Bacteria | Patescibacteria | Saccharimonadia | Saccharimonadales | NA | NA |
| 12 | FALSE | Archaea | Euryarchaeota | Methanobacteria | Methanobacteriales | Methanobacteriaceae | NA |
| 12 | FALSE | Bacteria | Firmicutes | Erysipelotrichia | Erysipelotrichales | Erysipelotrichaceae | Asteroleplasma |
| 12 | FALSE | Bacteria | Proteobacteria | Alphaproteobacteria | Rhizobiales | Rhizobiaceae | Mesorhizobium |
| 12 | FALSE | Bacteria | Proteobacteria | Alphaproteobacteria | Rickettsiales | Mitochondria | NA |
| 12 | FALSE | Bacteria | Proteobacteria | Gammaproteobacteria | Betaproteobacteriales | Rhodocyclaceae | Dechloromonas |
| 12 | FALSE | Bacteria | Bacteroidetes | Bacteroidia | Chitinophagales | Chitinophagaceae | Flavihumibacter |
| 12 | FALSE | Bacteria | Verrucomicrobia | Verrucomicrobiae | NA | NA | NA |
| 11 | FALSE | Bacteria | Proteobacteria | Gammaproteobacteria | Enterobacteriales | Enterobacteriaceae | Providencia |
| 11 | FALSE | Bacteria | Proteobacteria | Gammaproteobacteria | Aeromonadales | Aeromonadaceae | Aeromonas |
| 11 | FALSE | Bacteria | Proteobacteria | Gammaproteobacteria | Betaproteobacteriales | Burkholderiaceae | Achromobacter |
| 11 | FALSE | Bacteria | Firmicutes | Clostridia | Clostridiales | Family_XIII | S5-A14a |
| 11 | FALSE | Bacteria | Epsilonbacteraeota | Campylobacteria | Campylobacterales | Arcobacteraceae | Arcobacter |
| 11 | FALSE | Bacteria | Firmicutes | Clostridia | Clostridiales | Lachnospiraceae | UC5-1-2E3 |
| 11 | FALSE | Bacteria | Actinobacteria | Actinobacteria | Micrococcales | Micrococcaceae | Pseudoglutamicibacter |
| 11 | FALSE | Bacteria | Firmicutes | Clostridia | Clostridiales | Lachnospiraceae | Shuttleworthia |
| 11 | FALSE | Bacteria | Proteobacteria | Alphaproteobacteria | Sphingomonadales | Sphingomonadaceae | Sphingopyxis |
| 11 | FALSE | Bacteria | Actinobacteria | Actinobacteria | Streptomycetales | Streptomycetaceae | Streptomyces |
| 11 | FALSE | Bacteria | Synergistetes | Synergistia | Synergistales | Synergistaceae | Synergistes |
| 11 | FALSE | Bacteria | Firmicutes | Clostridia | Clostridiales | Lachnospiraceae | Anaerosporobacter |
| 11 | FALSE | Bacteria | Proteobacteria | Gammaproteobacteria | Betaproteobacteriales | Burkholderiaceae | Massilia |
| 11 | FALSE | Bacteria | Proteobacteria | Alphaproteobacteria | Rhizobiales | Rhizobiaceae | Pseudochrobactrum |
| 11 | FALSE | Bacteria | Bacteroidetes | Bacteroidia | Sphingobacteriales | Sphingobacteriaceae | Nubsella |
| 11 | FALSE | Bacteria | Firmicutes | Bacilli | Lactobacillales | Leuconostocaceae | Weissella |
| 11 | FALSE | Bacteria | Actinobacteria | Actinobacteria | Bifidobacteriales | Bifidobacteriaceae | Gardnerella |
| 11 | FALSE | Bacteria | Proteobacteria | Alphaproteobacteria | Caulobacterales | Caulobacteraceae | Caulobacter |
| 11 | FALSE | Bacteria | Firmicutes | Erysipelotrichia | Erysipelotrichales | Erysipelotrichaceae | Solobacterium |

| | | | | | | | |
|---|---|---|---|---|---|---|---|
| 11 | FALSE | Bacteria | Actinobacteria | Coriobacteriia | Coriobacteriales | Eggerthellaceae | Senegalimassilia |
| 11 | FALSE | Bacteria | Firmicutes | Clostridia | Clostridiales | Ruminococcaceae | Ruminococcaceae_UCG-011 |
| 11 | FALSE | Bacteria | Proteobacteria | Gammaproteobacteria | Betaproteobacteriales | Burkholderiaceae | Cupriavidus |
| 11 | FALSE | Bacteria | Firmicutes | Clostridia | Clostridiales | Defluviitaleaceae | Defluviitaleaceae_UCG-011 |
| 11 | FALSE | Bacteria | Actinobacteria | Actinobacteria | Micrococcales | Micrococcaceae | Rothia |
| 11 | FALSE | Bacteria | Proteobacteria | Alphaproteobacteria | Sphingomonadales | Sphingomonadaceae | Novosphingobium |
| 11 | FALSE | Bacteria | Proteobacteria | Gammaproteobacteria | Betaproteobacteriales | Burkholderiaceae | Aquabacterium |
| 11 | FALSE | Bacteria | Actinobacteria | Actinobacteria | Micrococcales | Microbacteriaceae | NA |
| 11 | FALSE | Bacteria | Verrucomicrobia | Verrucomicrobiae | Verrucomicrobiales | NA | NA |
| 10 | FALSE | Bacteria | Proteobacteria | Gammaproteobacteria | Aeromonadales | Succinivibrionaceae | Succinivibrio |
| 10 | FALSE | Bacteria | Proteobacteria | Gammaproteobacteria | Aeromonadales | Succinivibrionaceae | NA |
| 10 | FALSE | Bacteria | Firmicutes | Clostridia | Clostridiales | Peptococcaceae | NA |
| 10 | FALSE | Bacteria | Firmicutes | Clostridia | Clostridiales | Lachnospiraceae | Tyzzerella_3 |
| 10 | FALSE | Bacteria | Firmicutes | Bacilli | Bacillales | Planococcaceae | Lysinibacillus |
| 10 | FALSE | Bacteria | Fusobacteria | Fusobacteriia | Fusobacteriales | Leptotrichiaceae | Sneathia |
| 10 | FALSE | Bacteria | Proteobacteria | Alphaproteobacteria | Sphingomonadales | Sphingomonadaceae | Sphingobium |
| 10 | FALSE | Bacteria | Firmicutes | Clostridia | Clostridiales | Ruminococcaceae | Caproiciproducens |
| 10 | FALSE | Bacteria | Actinobacteria | Actinobacteria | Bifidobacteriales | Bifidobacteriaceae | Alloscardovia |
| 10 | FALSE | Bacteria | Actinobacteria | Coriobacteriia | Coriobacteriales | Eggerthellaceae | Adlercreutzia |
| 10 | FALSE | Bacteria | Bacteroidetes | Bacteroidia | Bacteroidales | Rikenellaceae | Millionella |
| 10 | FALSE | Bacteria | Firmicutes | Clostridia | Clostridiales | Clostridiaceae_1 | Sarcina |
| 10 | FALSE | Bacteria | Lentisphaerae | Lentisphaeria | Victivallales | NA | NA |
| 10 | FALSE | Bacteria | Synergistetes | Synergistia | Synergistales | Synergistaceae | Pyramidobacter |
| 10 | FALSE | Bacteria | Synergistetes | Synergistia | Synergistales | Synergistaceae | Jonquetella |
| 10 | FALSE | Bacteria | Actinobacteria | Coriobacteriia | Coriobacteriales | Atopobiaceae | Olsenella |
| 10 | FALSE | Bacteria | Firmicutes | Bacilli | Bacillales | Family_XI | Gemella |
| 10 | FALSE | Bacteria | Elusimicrobia | Elusimicrobia | Elusimicrobiales | Elusimicrobiaceae | Elusimicrobium |
| 10 | FALSE | Bacteria | Proteobacteria | Alphaproteobacteria | Rhodobacterales | Rhodobacteraceae | Paracoccus |
| 10 | FALSE | Bacteria | Firmicutes | Negativicutes | Selenomonadales | Acidaminococcaceae | NA |
| 10 | FALSE | Bacteria | Firmicutes | Negativicutes | Selenomonadales | NA | NA |
| 10 | FALSE | Bacteria | Proteobacteria | Gammaproteobacteria | Enterobacteriales | Enterobacteriaceae | Cosenzaea |
| 10 | FALSE | Bacteria | Actinobacteria | Actinobacteria | Corynebacteriales | NA | NA |
| 10 | FALSE | Bacteria | Proteobacteria | NA | NA | NA | NA |
| 10 | FALSE | Bacteria | Proteobacteria | Alphaproteobacteria | Rhodobacterales | Rhodobacteraceae | NA |
| 10 | FALSE | Bacteria | Firmicutes | Bacilli | Lactobacillales | Streptococcaceae | NA |
| 9 | FALSE | Bacteria | Bacteroidetes | Bacteroidia | Bacteroidales | Bacteroidaceae | Bacteroides |
| 9 | FALSE | Bacteria | Bacteroidetes | Bacteroidia | Bacteroidales | Prevotellaceae | Alloprevotella |
| 9 | FALSE | Bacteria | Firmicutes | Clostridia | Clostridiales | Lachnospiraceae | Lachnospiraceae_UCG-003 |
| 9 | FALSE | Bacteria | Bacteroidetes | Bacteroidia | Bacteroidales | Barnesiellaceae | Coprobacter |
| 9 | FALSE | Bacteria | Firmicutes | Clostridia | Clostridiales | Family_XI | Murdochiella |
| 9 | FALSE | Bacteria | Proteobacteria | Gammaproteobacteria | Betaproteobacteriales | Burkholderiaceae | Ottowia |

| | | | | | | | |
|---|---|---|---|---|---|---|---|
| 9 | FALSE | Bacteria | Firmicutes | Erysipelotrichia | Erysipelotrichales | Erysipelotrichaceae | Merdibacter |
| 9 | FALSE | Bacteria | Firmicutes | Clostridia | DTU014 | NA | NA |
| 9 | FALSE | Bacteria | Firmicutes | Clostridia | Clostridiales | Ruminococcaceae | Anaerofilum |
| 9 | FALSE | Archaea | Euryarchaeota | Methanobacteria | Methanobacteriales | Methanobacteriaceae | Methanosphaera |
| 9 | FALSE | Bacteria | Bacteroidetes | Bacteroidia | Bacteroidales | Rikenellaceae | NA |
| 9 | FALSE | Bacteria | Actinobacteria | Actinobacteria | Corynebacteriales | Corynebacteriaceae | Corynebacterium |
| 8 | FALSE | Bacteria | Firmicutes | Clostridia | Clostridiales | Ruminococcaceae | Ruminococcaceae_UCG-005 |
| 8 | FALSE | Bacteria | Proteobacteria | Gammaproteobacteria | Betaproteobacteriales | Burkholderiaceae | Variovorax |
| 8 | FALSE | Bacteria | Proteobacteria | Gammaproteobacteria | Betaproteobacteriales | Burkholderiaceae | Oligella |
| 8 | FALSE | Bacteria | Proteobacteria | Gammaproteobacteria | Xanthomonadales | Xanthomonadaceae | NA |
| 8 | FALSE | Bacteria | Bacteroidetes | Bacteroidia | Bacteroidales | Muribaculaceae | CAG-873 |
| 8 | FALSE | Bacteria | Proteobacteria | Alphaproteobacteria | Rhizobiales | Rhizobiaceae | NA |
| 8 | FALSE | Bacteria | Firmicutes | Bacilli | Bacillales | Paenibacillaceae | Brevibacillus |
| 8 | FALSE | Bacteria | Proteobacteria | Gammaproteobacteria | Betaproteobacteriales | Burkholderiaceae | Oxalobacter |
| 8 | FALSE | Bacteria | Firmicutes | Clostridia | Clostridiales | Christensenellaceae | NA |
| 8 | FALSE | Bacteria | Firmicutes | Bacilli | Bacillales | Planococcaceae | Rummeliibacillus |
| 8 | FALSE | Bacteria | Firmicutes | Erysipelotrichia | Erysipelotrichales | Erysipelotrichaceae | Faecalicoccus |
| 8 | FALSE | Bacteria | Firmicutes | Clostridia | Clostridiales | Ruminococcaceae | Ruminococcaceae_UCG-009 |
| 8 | FALSE | Bacteria | Firmicutes | Clostridia | Clostridiales | Clostridiaceae_1 | Clostridium_sensu_stricto_13 |
| 8 | FALSE | Bacteria | Firmicutes | Clostridia | Clostridiales | Lachnospiraceae | Cellulosilyticum |
| 8 | FALSE | Bacteria | Proteobacteria | Deltaproteobacteria | Desulfovibrionales | Desulfovibrionaceae | Mailhella |
| 8 | FALSE | Bacteria | Actinobacteria | Coriobacteriia | Coriobacteriales | Eggerthellaceae | Gordonibacter |
| 8 | FALSE | Bacteria | Firmicutes | Clostridia | Clostridiales | Lachnospiraceae | Oribacterium |
| 8 | FALSE | Bacteria | Firmicutes | Bacilli | Bacillales | Staphylococcaceae | Nosocomiicoccus |
| 8 | FALSE | Bacteria | Actinobacteria | Coriobacteriia | Coriobacteriales | NA | NA |
| 8 | FALSE | Bacteria | Lentisphaerae | Lentisphaeria | Victivallales | Victivallaceae | NA |
| 8 | FALSE | Bacteria | Firmicutes | Erysipelotrichia | Erysipelotrichales | Erysipelotrichaceae | Allobaculum |
| 8 | FALSE | Bacteria | Proteobacteria | Alphaproteobacteria | Sphingomonadales | Sphingomonadaceae | Sphingomonas |
| 8 | FALSE | Bacteria | Firmicutes | Clostridia | Clostridiales | Clostridiaceae_1 | NA |
| 7 | FALSE | Bacteria | Firmicutes | Clostridia | Clostridiales | Lachnospiraceae | Dorea |
| 7 | FALSE | Bacteria | Bacteroidetes | Bacteroidia | Bacteroidales | Prevotellaceae | Prevotellaceae_NK3B31_group |
| 7 | FALSE | Bacteria | Proteobacteria | Deltaproteobacteria | Desulfovibrionales | Desulfovibrionaceae | Desulfovibrio |
| 7 | FALSE | Bacteria | Firmicutes | Erysipelotrichia | Erysipelotrichales | Erysipelotrichaceae | Erysipelotrichaceae_UCG-004 |
| 7 | FALSE | Bacteria | Firmicutes | Erysipelotrichia | Erysipelotrichales | Erysipelotrichaceae | Candidatus_Stoquefichus |
| 7 | FALSE | Bacteria | Firmicutes | Erysipelotrichia | Erysipelotrichales | Erysipelotrichaceae | Holdemania |
| 7 | FALSE | Bacteria | Proteobacteria | Gammaproteobacteria | Betaproteobacteriales | Burkholderiaceae | Acidovorax |
| 7 | FALSE | Bacteria | Firmicutes | Clostridia | Clostridiales | Lachnospiraceae | GCA-900066575 |
| 7 | FALSE | Bacteria | Firmicutes | Clostridia | Clostridiales | Family_XI | W5053 |
| 7 | FALSE | Bacteria | Firmicutes | Negativicutes | Selenomonadales | Veillonellaceae | Negativicoccus |
| 7 | FALSE | Bacteria | Firmicutes | Negativicutes | Selenomonadales | Veillonellaceae | Anaeroglobus |
| 7 | FALSE | Bacteria | Bacteroidetes | Bacteroidia | Bacteroidales | Marinifilaceae | Sanguibacteroides |

| | | | | | | | |
|---|---|---|---|---|---|---|---|
| 7 | FALSE | Bacteria | Firmicutes | Clostridia | Clostridiales | Peptostreptococcaceae | NA |
| 6 | FALSE | Bacteria | Firmicutes | Clostridia | Clostridiales | Ruminococcaceae | Intestinimonas |
| 6 | FALSE | Bacteria | Actinobacteria | Actinobacteria | Corynebacteriales | Corynebacteriaceae | NA |
| 6 | FALSE | Bacteria | Firmicutes | Bacilli | Lactobacillales | Lactobacillaceae | Pediococcus |
| 6 | FALSE | Bacteria | Firmicutes | Clostridia | Clostridiales | Ruminococcaceae | Fastidiosipila |
| 6 | FALSE | Bacteria | Firmicutes | Clostridia | Clostridiales | Family_XI | Gallicola |
| 6 | FALSE | Archaea | Euryarchaeota | Thermoplasmata | Methanomassiliicoccales | Methanomassiliicoccaceae | Methanomassiliicoccus |
| 5 | FALSE | Bacteria | Firmicutes | Clostridia | Clostridiales | Lachnospiraceae | Lachnoclostridium |
| 5 | FALSE | Bacteria | Bacteroidetes | Bacteroidia | Flavobacteriales | Flavobacteriaceae | Flavobacterium |
| 5 | FALSE | Bacteria | Proteobacteria | Gammaproteobacteria | Betaproteobacteriales | Burkholderiaceae | NA |
| 5 | FALSE | Bacteria | Firmicutes | Clostridia | Clostridiales | Clostridiales_vadinBB60_group | NA |
| 5 | FALSE | Bacteria | Firmicutes | Negativicutes | Selenomonadales | Veillonellaceae | Megamonas |
| 5 | FALSE | Bacteria | Firmicutes | Erysipelotrichia | Erysipelotrichales | Erysipelotrichaceae | Erysipelatoclostridium |
| 5 | FALSE | Bacteria | Firmicutes | Negativicutes | Selenomonadales | Veillonellaceae | NA |
| 5 | FALSE | Bacteria | Bacteroidetes | Bacteroidia | Bacteroidales | Dysgonomonadaceae | Dysgonomonas |
| 5 | FALSE | Bacteria | Firmicutes | Clostridia | Clostridiales | Peptostreptococcaceae | Terrisporobacter |
| 5 | FALSE | Bacteria | Firmicutes | Clostridia | Clostridiales | Ruminococcaceae | Ruminococcaceae_UCG-010 |
| 5 | FALSE | Bacteria | Bacteroidetes | Bacteroidia | Bacteroidales | Barnesiellaceae | NA |
| 5 | FALSE | Bacteria | Firmicutes | Clostridia | Clostridiales | Ruminococcaceae | GCA-900066225 |
| 5 | FALSE | Bacteria | Proteobacteria | Gammaproteobacteria | Xanthomonadales | Xanthomonadaceae | Pseudoxanthomonas |
| 5 | FALSE | Bacteria | Firmicutes | Clostridia | Clostridiales | Lachnospiraceae | Marvinbryantia |
| 5 | FALSE | Bacteria | Firmicutes | Bacilli | Lactobacillales | Streptococcaceae | Lactococcus |
| 5 | FALSE | Bacteria | Actinobacteria | Coriobacteriia | Coriobacteriales | Eggerthellaceae | NA |
| 5 | FALSE | Bacteria | Firmicutes | Clostridia | Clostridiales | Ruminococcaceae | Angelakisella |
| 5 | FALSE | Bacteria | Firmicutes | Negativicutes | Selenomonadales | Veillonellaceae | Allisonella |
| 5 | FALSE | Bacteria | Firmicutes | Clostridia | Clostridiales | Peptococcaceae | Peptococcus |
| 5 | FALSE | Bacteria | Firmicutes | Bacilli | Lactobacillales | Lactobacillaceae | NA |
| 5 | FALSE | Archaea | Euryarchaeota | Thermoplasmata | Methanomassiliicoccales | Methanomethylophilaceae | NA |
| 5 | FALSE | Bacteria | Firmicutes | Clostridia | Clostridiales | Lachnospiraceae | Howardella |
| 5 | FALSE | Bacteria | Actinobacteria | Actinobacteria | Micrococcales | Brevibacteriaceae | Brevibacterium |
| 5 | FALSE | Bacteria | Actinobacteria | Actinobacteria | Actinomycetales | Actinomycetaceae | Actinomyces |
| 5 | FALSE | Bacteria | Firmicutes | Clostridia | Clostridiales | Eubacteriaceae | Eubacterium |
| 5 | FALSE | Bacteria | Actinobacteria | Coriobacteriia | Coriobacteriales | Atopobiaceae | Atopobium |
| 5 | FALSE | Bacteria | Actinobacteria | Coriobacteriia | Coriobacteriales | Eggerthellaceae | Slackia |
| 4 | FALSE | Bacteria | Firmicutes | Clostridia | Clostridiales | Ruminococcaceae | Oscillibacter |
| 4 | FALSE | Bacteria | Firmicutes | Clostridia | Clostridiales | Clostridiaceae_1 | Clostridium_sensu_stricto_1 |
| 4 | FALSE | Bacteria | Firmicutes | Clostridia | Clostridiales | Family_XI | Finegoldia |
| 4 | FALSE | Bacteria | Firmicutes | Clostridia | Clostridiales | Family_XIII | Family_XIII_AD3011_group |
| 4 | FALSE | Bacteria | Firmicutes | Clostridia | Clostridiales | Lachnospiraceae | CAG-56 |
| 4 | FALSE | Bacteria | Firmicutes | Erysipelotrichia | Erysipelotrichales | Erysipelotrichaceae | Dielma |
| 4 | FALSE | Bacteria | Firmicutes | Clostridia | Clostridiales | Ruminococcaceae | Ruminiclostridium_1 |

| | | | | | | | |
|---|---|---|---|---|---|---|---|
| 4 | FALSE | Bacteria | Firmicutes | Clostridia | Clostridiales | Family_XIII | NA |
| 4 | FALSE | Bacteria | Bacteroidetes | Bacteroidia | Sphingobacteriales | Sphingobacteriaceae | Sphingobacterium |
| 4 | FALSE | Bacteria | Bacteroidetes | Bacteroidia | NA | NA | NA |
| 4 | FALSE | Bacteria | Lentisphaerae | Lentisphaeria | Victivallales | Victivallaceae | Victivallis |
| 4 | FALSE | Bacteria | Actinobacteria | Coriobacteriia | Coriobacteriales | Coriobacteriales_Incertae_Sedis | NA |
| 4 | FALSE | Bacteria | Bacteroidetes | Bacteroidia | Flavobacteriales | Weeksellaceae | Cloacibacterium |
| 4 | FALSE | Bacteria | Firmicutes | Clostridia | Clostridiales | Ruminococcaceae | Hydrogenoanaerobacterium |
| 4 | FALSE | Bacteria | Actinobacteria | Coriobacteriia | Coriobacteriales | Eggerthellaceae | Eggerthella |
| 4 | FALSE | Bacteria | Bacteroidetes | Bacteroidia | Bacteroidales | Prevotellaceae | NA |
| 4 | FALSE | Bacteria | Firmicutes | NA | NA | NA | NA |
| 4 | FALSE | Bacteria | Firmicutes | Clostridia | Clostridiales | Family_XIII | Mogibacterium |
| 4 | FALSE | Bacteria | Firmicutes | Bacilli | Bacillales | Staphylococcaceae | Staphylococcus |
| 4 | FALSE | Bacteria | Proteobacteria | Gammaproteobacteria | Pseudomonadales | Pseudomonadaceae | NA |
| 3 | FALSE | Bacteria | Proteobacteria | Gammaproteobacteria | Enterobacteriales | Enterobacteriaceae | Proteus |
| 3 | FALSE | Bacteria | Firmicutes | Negativicutes | Selenomonadales | Acidaminococcaceae | Acidaminococcus |
| 3 | FALSE | Bacteria | Firmicutes | Clostridia | Clostridiales | Christensenellaceae | Christensenellaceae_R-7_group |
| 3 | FALSE | Bacteria | Firmicutes | Clostridia | Clostridiales | Ruminococcaceae | Ruminococcaceae_UCG-003 |
| 3 | FALSE | Bacteria | Proteobacteria | Gammaproteobacteria | Enterobacteriales | Enterobacteriaceae | Morganella |
| 3 | FALSE | Bacteria | Firmicutes | Clostridia | Clostridiales | Ruminococcaceae | Ruminococcaceae_NK4A214_group |
| 3 | FALSE | Bacteria | Bacteroidetes | Bacteroidia | Bacteroidales | Marinifilaceae | Odoribacter |
| 3 | FALSE | Bacteria | Firmicutes | Clostridia | Clostridiales | Peptostreptococcaceae | Intestinibacter |
| 3 | FALSE | Bacteria | Firmicutes | Clostridia | Clostridiales | Ruminococcaceae | CAG-352 |
| 3 | FALSE | Bacteria | Bacteroidetes | Bacteroidia | Bacteroidales | Prevotellaceae | Prevotella_7 |
| 3 | FALSE | Bacteria | Bacteroidetes | Bacteroidia | Bacteroidales | Rikenellaceae | Rikenellaceae_RC9_gut_group |
| 3 | FALSE | Bacteria | Firmicutes | Erysipelotrichia | Erysipelotrichales | Erysipelotrichaceae | Catenibacterium |
| 3 | FALSE | Bacteria | Firmicutes | Clostridia | Clostridiales | Lachnospiraceae | Coprococcus_2 |
| 3 | FALSE | Bacteria | Firmicutes | Erysipelotrichia | Erysipelotrichales | Erysipelotrichaceae | Faecalitalea |
| 3 | FALSE | Bacteria | Bacteroidetes | Bacteroidia | Bacteroidales | Marinifilaceae | Butyricimonas |
| 3 | FALSE | Bacteria | Bacteroidetes | Bacteroidia | Flavobacteriales | Weeksellaceae | Chryseobacterium |
| 3 | FALSE | Bacteria | Bacteroidetes | Bacteroidia | Bacteroidales | Prevotellaceae | Prevotella_2 |
| 3 | FALSE | Bacteria | Firmicutes | Negativicutes | Selenomonadales | Veillonellaceae | Veillonella |
| 3 | FALSE | Bacteria | Firmicutes | Bacilli | Bacillales | Planococcaceae | NA |
| 3 | FALSE | Bacteria | Firmicutes | Bacilli | Bacillales | NA | NA |
| 3 | FALSE | Bacteria | Lentisphaerae | Lentisphaeria | Victivallales | vadinBE97 | NA |
| 3 | FALSE | Bacteria | Proteobacteria | Deltaproteobacteria | Desulfovibrionales | Desulfovibrionaceae | NA |
| 2 | FALSE | Bacteria | Firmicutes | Negativicutes | Selenomonadales | Acidaminococcaceae | Phascolarctobacterium |
| 2 | FALSE | Bacteria | Firmicutes | Clostridia | Clostridiales | Peptostreptococcaceae | Romboutsia |
| 2 | FALSE | Bacteria | Firmicutes | Clostridia | Clostridiales | Ruminococcaceae | Ruminiclostridium_5 |
| 2 | FALSE | Bacteria | Firmicutes | Erysipelotrichia | Erysipelotrichales | Erysipelotrichaceae | Turicibacter |
| 2 | FALSE | Bacteria | Proteobacteria | Gammaproteobacteria | Xanthomonadales | Xanthomonadaceae | Stenotrophomonas |
| 2 | FALSE | Bacteria | Firmicutes | Clostridia | Clostridiales | Lachnospiraceae | Lachnospiraceae_UCG-001 |

| | | | | | | | |
|---|---|---|---|---|---|---|---|
| 2 | FALSE | Bacteria | Firmicutes | Clostridia | Clostridiales | Ruminococcaceae | DTU089 |
| 2 | FALSE | Bacteria | Proteobacteria | Gammaproteobacteria | NA | NA | NA |
| 1 | FALSE | Bacteria | Bacteroidetes | Bacteroidia | Bacteroidales | Tannerellaceae | Parabacteroides |
| 1 | FALSE | Bacteria | Firmicutes | Negativicutes | Selenomonadales | Veillonellaceae | Dialister |
| 1 | FALSE | Bacteria | Firmicutes | Clostridia | Clostridiales | Ruminococcaceae | Ruminococcaceae_UCG-002 |
| 1 | FALSE | Bacteria | Proteobacteria | Gammaproteobacteria | Betaproteobacteriales | Burkholderiaceae | Parasutterella |
| 1 | FALSE | Bacteria | Firmicutes | Clostridia | Clostridiales | Lachnospiraceae | Butyrivibrio |
| 1 | FALSE | Bacteria | Firmicutes | Clostridia | Clostridiales | Lachnospiraceae | Lachnospiraceae_NK4A136_group |
| 1 | FALSE | Bacteria | Firmicutes | Clostridia | Clostridiales | Lachnospiraceae | Eisenbergiella |
| 1 | FALSE | Bacteria | Firmicutes | Erysipelotrichia | Erysipelotrichales | Erysipelotrichaceae | Holdemanella |
| 1 | FALSE | Bacteria | Actinobacteria | Coriobacteriia | Coriobacteriales | Coriobacteriaceae | Collinsella |
| 1 | FALSE | Bacteria | Tenericutes | Mollicutes | Izimaplasmatales | NA | NA |
| 1 | FALSE | Bacteria | Fusobacteria | Fusobacteriia | Fusobacteriales | Fusobacteriaceae | Fusobacterium |
| 1 | FALSE | Bacteria | Proteobacteria | Alphaproteobacteria | Caulobacterales | Caulobacteraceae | Brevundimonas |
| 1 | FALSE | Bacteria | Firmicutes | Clostridia | Clostridiales | Lachnospiraceae | Coprococcus_1 |
| 1 | FALSE | Bacteria | Cyanobacteria | Melainabacteria | Gastranaerophilales | NA | NA |
| 1 | FALSE | Bacteria | NA | NA | NA | NA | NA |
| 0 | FALSE | Bacteria | Proteobacteria | Gammaproteobacteria | Enterobacteriales | Enterobacteriaceae | Escherichia/Shigella |
| 0 | FALSE | Bacteria | Proteobacteria | Gammaproteobacteria | Enterobacteriales | Enterobacteriaceae | NA |
| 0 | FALSE | Bacteria | Firmicutes | Clostridia | Clostridiales | Ruminococcaceae | Ruminococcus_2 |
| 0 | FALSE | Bacteria | Firmicutes | Clostridia | Clostridiales | Ruminococcaceae | Subdoligranulum |
| 0 | FALSE | Bacteria | Bacteroidetes | Bacteroidia | Bacteroidales | Prevotellaceae | Prevotella_9 |
| 0 | FALSE | Bacteria | Proteobacteria | Gammaproteobacteria | Pseudomonadales | Pseudomonadaceae | Pseudomonas |
| 0 | FALSE | Bacteria | Proteobacteria | Gammaproteobacteria | Pseudomonadales | Moraxellaceae | Acinetobacter |
| 0 | FALSE | Bacteria | Firmicutes | Clostridia | Clostridiales | Lachnospiraceae | Tyzzerella_4 |
| 0 | FALSE | Bacteria | Proteobacteria | Gammaproteobacteria | Enterobacteriales | Enterobacteriaceae | Klebsiella |
| 0 | FALSE | Bacteria | Firmicutes | Clostridia | Clostridiales | Ruminococcaceae | Ruminiclostridium_6 |
| 0 | FALSE | Bacteria | Firmicutes | Clostridia | Clostridiales | Ruminococcaceae | Ruminococcus_1 |
| 0 | FALSE | Bacteria | Firmicutes | Erysipelotrichia | Erysipelotrichales | Erysipelotrichaceae | Erysipelotrichaceae_UCG-003 |
| 0 | FALSE | Bacteria | Firmicutes | Clostridia | Clostridiales | Ruminococcaceae | Flavonifractor |
| 0 | FALSE | Bacteria | Firmicutes | Bacilli | Lactobacillales | Streptococcaceae | Streptococcus |
| 0 | FALSE | Bacteria | Firmicutes | Clostridia | Clostridiales | Lachnospiraceae | Tyzzerella |
| 0 | FALSE | Bacteria | Proteobacteria | Gammaproteobacteria | Betaproteobacteriales | Burkholderiaceae | Delftia |
| 0 | FALSE | Bacteria | Proteobacteria | Gammaproteobacteria | Betaproteobacteriales | Burkholderiaceae | Sutterella |
| 0 | FALSE | Bacteria | Bacteroidetes | Bacteroidia | Bacteroidales | Barnesiellaceae | Barnesiella |
| 0 | FALSE | Bacteria | Proteobacteria | Alphaproteobacteria | Rhodospirillales | NA | NA |
| 0 | FALSE | Bacteria | Firmicutes | Clostridia | Clostridiales | Ruminococcaceae | Ruminiclostridium_9 |
| 0 | FALSE | Bacteria | Firmicutes | Clostridia | Clostridiales | Ruminococcaceae | Ruminococcaceae_UCG-014 |
| 0 | FALSE | Bacteria | Bacteroidetes | Bacteroidia | Bacteroidales | Prevotellaceae | Paraprevotella |
| 0 | FALSE | Bacteria | Firmicutes | Clostridia | Clostridiales | Ruminococcaceae | Negativibacillus |
| 0 | FALSE | Bacteria | Tenericutes | Mollicutes | Mollicutes_RF39 | NA | NA |

| | | | | | | | |
|---|---|---|---|---|---|---|---|
| 0 | FALSE | Bacteria | Firmicutes | Bacilli | Lactobacillales | Enterococcaceae | Enterococcus |
| 0 | FALSE | Bacteria | Bacteroidetes | Bacteroidia | Bacteroidales | Muribaculaceae | NA |
| 0 | FALSE | Bacteria | Firmicutes | Clostridia | Clostridiales | Lachnospiraceae | Lachnospiraceae_UCG-010 |

**Supplementary Table 3. MWAS of dataset 2 conducted using ANCOM**

W= ANCOM score indicating the number of times a genus achieved FDR<0.05 as compared to other genera (maximum W possible: 444 in dataset 1, 560 in dataset 2).

0.8= Threshold at which results were considered significant (TRUE).

| W | 0.8 | Kingdom | Phylum | Class | Order | Family | Genus |
|---|---|---|---|---|---|---|---|
| 553 | TRUE | Bacteria | Actinobacteria | Actinobacteria | Bifidobacteriales | Bifidobacteriaceae | Bifidobacterium |
| 545 | TRUE | Bacteria | Firmicutes | Clostridia | Clostridiales | Lachnospiraceae | Agathobacter |
| 544 | TRUE | Bacteria | Firmicutes | Clostridia | Clostridiales | Lachnospiraceae | Lachnospiraceae_UCG-004 |
| 541 | TRUE | Bacteria | Firmicutes | Clostridia | Clostridiales | Lachnospiraceae | Roseburia |
| 541 | TRUE | Bacteria | Firmicutes | Clostridia | Clostridiales | Ruminococcaceae | Ruminococcaceae_UCG-013 |
| 538 | TRUE | Bacteria | Firmicutes | Clostridia | Clostridiales | Lachnospiraceae | Lachnospiraceae_ND3007_group |
| 536 | TRUE | Bacteria | Firmicutes | Clostridia | Clostridiales | Lachnospiraceae | Anaerostipes |
| 535 | TRUE | Bacteria | Firmicutes | Clostridia | Clostridiales | Ruminococcaceae | Faecalibacterium |
| 533 | TRUE | Bacteria | Firmicutes | Clostridia | Clostridiales | Lachnospiraceae | Blautia |
| 530 | TRUE | Bacteria | Firmicutes | Clostridia | Clostridiales | Eubacteriaceae | Eubacterium |
| 525 | TRUE | Bacteria | Firmicutes | Clostridia | Clostridiales | Ruminococcaceae | Oscillospira |
| 524 | TRUE | Bacteria | Firmicutes | Clostridia | Clostridiales | Ruminococcaceae | Ruminococcus_2 |
| 521 | TRUE | Bacteria | Firmicutes | Clostridia | Clostridiales | Lachnospiraceae | Fusicatenibacter |
| 521 | TRUE | Bacteria | Firmicutes | Clostridia | Clostridiales | Lachnospiraceae | Lachnospira |
| 521 | TRUE | Bacteria | Firmicutes | Clostridia | Clostridiales | Ruminococcaceae | Ruminiclostridium_6 |
| 521 | TRUE | Bacteria | Firmicutes | Clostridia | Clostridiales | Ruminococcaceae | Ruminococcus_1 |
| 505 | TRUE | Bacteria | Firmicutes | Clostridia | Clostridiales | Ruminococcaceae | Butyricicoccus |
| 505 | TRUE | Bacteria | Proteobacteria | Gammaproteobacteria | Pseudomonadales | Pseudomonadaceae | Pseudomonas |
| 503 | TRUE | Bacteria | Firmicutes | Clostridia | Clostridiales | Lachnospiraceae | Lachnospiraceae_UCG-001 |
| 496 | TRUE | Bacteria | Actinobacteria | Actinobacteria | Corynebacteriales | Corynebacteriaceae | Lawsonella |
| 493 | TRUE | Bacteria | Proteobacteria | Deltaproteobacteria | Desulfovibrionales | Desulfovibrionaceae | Desulfovibrio |
| 493 | TRUE | Bacteria | Firmicutes | Erysipelotrichia | Erysipelotrichales | Erysipelotrichaceae | Turicibacter |
| 491 | TRUE | Archaea | Euryarchaeota | Methanobacteria | Methanobacteriales | Methanobacteriaceae | Methanobrevibacter |
| 479 | TRUE | Bacteria | Firmicutes | Clostridia | Clostridiales | Ruminococcaceae | DTU089 |
| 477 | TRUE | Bacteria | Firmicutes | Erysipelotrichia | Erysipelotrichales | Erysipelotrichaceae | Erysipelotrichaceae_UCG-003 |
| 468 | TRUE | Bacteria | Bacteroidetes | Bacteroidia | Bacteroidales | Porphyromonadaceae | Porphyromonas |
| 465 | TRUE | Bacteria | Actinobacteria | Actinobacteria | Corynebacteriales | Corynebacteriaceae | Corynebacterium_1 |

| | | | | | | | |
|---|---|---|---|---|---|---|---|
| 463 | TRUE | Bacteria | Bacteroidetes | Bacteroidia | Bacteroidales | Prevotellaceae | Prevotella |
| 459 | TRUE | Bacteria | Firmicutes | Clostridia | Clostridiales | Lachnospiraceae | Lachnoclostridium |
| 458 | TRUE | Bacteria | Firmicutes | Bacilli | Lactobacillales | Lactobacillaceae | Lactobacillus |
| 458 | TRUE | Bacteria | Firmicutes | Clostridia | Clostridiales | Ruminococcaceae | Ruminococcaceae_UCG-014 |
| 454 | TRUE | Bacteria | Firmicutes | Negativicutes | Selenomonadales | Veillonellaceae | Veillonella |
| 452 | TRUE | Bacteria | Firmicutes | Clostridia | Clostridiales | Lachnospiraceae | NA |
| 449 | TRUE | Bacteria | Firmicutes | Clostridia | Clostridiales | Ruminococcaceae | Candidatus_Soleaferrea |
| 440 | FALSE | Bacteria | Proteobacteria | Gammaproteobacteria | Pseudomonadales | Moraxellaceae | Acinetobacter |
| 439 | FALSE | Bacteria | Firmicutes | Erysipelotrichia | Erysipelotrichales | Erysipelotrichaceae | NA |
| 438 | FALSE | Bacteria | Firmicutes | Clostridia | Clostridiales | Ruminococcaceae | Intestinimonas |
| 433 | FALSE | Bacteria | Firmicutes | Clostridia | Clostridiales | Ruminococcaceae | Ruminiclostridium_9 |
| 428 | FALSE | Bacteria | Firmicutes | Clostridia | Clostridiales | Family_XI | Anaerococcus |
| 425 | FALSE | Bacteria | Firmicutes | Erysipelotrichia | Erysipelotrichales | Erysipelotrichaceae | Erysipelatoclostridium |
| 422 | FALSE | Bacteria | Firmicutes | Clostridia | Clostridiales | Family_XIII | Family_XIII_UCG-001 |
| 411 | FALSE | Bacteria | Firmicutes | Clostridia | Clostridiales | Ruminococcaceae | Phocea |
| 408 | FALSE | Bacteria | Cyanobacteria | Oxyphotobacteria | Chloroplast | NA | NA |
| 408 | FALSE | Bacteria | Firmicutes | Clostridia | Clostridiales | Family_XIII | S5-A14a |
| 407 | FALSE | Bacteria | Bacteroidetes | Bacteroidia | Bacteroidales | Bacteroidaceae | Bacteroides |
| 405 | FALSE | Bacteria | Firmicutes | Erysipelotrichia | Erysipelotrichales | Erysipelotrichaceae | Holdemania |
| 399 | FALSE | Bacteria | Firmicutes | Bacilli | Lactobacillales | Carnobacteriaceae | Granulicatella |
| 395 | FALSE | Bacteria | Firmicutes | Clostridia | Clostridiales | Lachnospiraceae | Cuneatibacter |
| 388 | FALSE | Bacteria | Firmicutes | Clostridia | Clostridiales | Lachnospiraceae | Lachnospiraceae_UCG-003 |
| 386 | FALSE | Bacteria | Firmicutes | Clostridia | Clostridiales | Lachnospiraceae | GCA-900066575 |
| 364 | FALSE | Bacteria | Proteobacteria | Gammaproteobacteria | Betaproteobacteriales | Burkholderiaceae | Delftia |
| 356 | FALSE | Bacteria | Firmicutes | Clostridia | Clostridiales | Family_XI | Peptoniphilus |
| 350 | FALSE | Bacteria | Firmicutes | Clostridia | Clostridiales | Lachnospiraceae | Lachnospiraceae_UCG-008 |
| 348 | FALSE | Bacteria | Firmicutes | Clostridia | Clostridiales | Family_XI | Parvimonas |
| 344 | FALSE | Bacteria | Firmicutes | Bacilli | Bacillales | Family_XI | Gemella |
| 307 | FALSE | Bacteria | Actinobacteria | Actinobacteria | Actinomycetales | Actinomycetaceae | Varibaculum |
| 299 | FALSE | Bacteria | Proteobacteria | Gammaproteobacteria | Xanthomonadales | Xanthomonadaceae | Stenotrophomonas |
| 268 | FALSE | Bacteria | Firmicutes | Bacilli | Bacillales | Bacillaceae | Bacillus |
| 218 | FALSE | Bacteria | Firmicutes | Negativicutes | Selenomonadales | Veillonellaceae | Selenomonas_3 |
| 209 | FALSE | Bacteria | Firmicutes | Erysipelotrichia | Erysipelotrichales | Erysipelotrichaceae | Asteroleplasma |
| 62 | FALSE | Bacteria | Lentisphaerae | Lentisphaeria | Victivallales | Victivallaceae | Victivallis |
| 58 | FALSE | Bacteria | Actinobacteria | Coriobacteriia | Coriobacteriales | Atopobiaceae | Olsenella |

| | | | | | | | |
|---|---|---|---|---|---|---|---|
| 54 | FALSE | Bacteria | Actinobacteria | Actinobacteria | Propionibacteriales | Propionibacteriaceae | Tessaracoccus |
| 50 | FALSE | Bacteria | Epsilonbacteraeota | Campylobacteria | Campylobacterales | Campylobacteraceae | Campylobacter |
| 49 | FALSE | Bacteria | Firmicutes | Clostridia | Clostridiales | Family_XI | Ezakiella |
| 47 | FALSE | Bacteria | Proteobacteria | Alphaproteobacteria | Rhizobiales | Beijerinckiaceae | Bosea |
| 47 | FALSE | Bacteria | Proteobacteria | Alphaproteobacteria | Rhizobiales | Beijerinckiaceae | Methylobacterium |
| 46 | FALSE | Bacteria | Firmicutes | Clostridia | Clostridiales | Peptostreptococcaceae | NA |
| 44 | FALSE | Bacteria | Actinobacteria | Actinobacteria | Propionibacteriales | Propionibacteriaceae | Cutibacterium |
| 44 | FALSE | Bacteria | Proteobacteria | Alphaproteobacteria | Rhizobiales | Rhizobiaceae | NA |
| 43 | FALSE | Bacteria | Firmicutes | Clostridia | Clostridiales | Clostridiaceae_1 | Clostridium_sensu_stricto_13 |
| 41 | FALSE | Bacteria | Firmicutes | Bacilli | Lactobacillales | Lactobacillaceae | Pediococcus |
| 40 | FALSE | Bacteria | Firmicutes | Clostridia | Clostridiales | Ruminococcaceae | Fastidiosipila |
| 40 | FALSE | Bacteria | Actinobacteria | Actinobacteria | Actinomycetales | Actinomycetaceae | Mobiluncus |
| 40 | FALSE | Bacteria | Actinobacteria | Actinobacteria | Micrococcales | Microbacteriaceae | Pseudoclavibacter |
| 39 | FALSE | Bacteria | Firmicutes | Clostridia | Clostridiales | Family_XI | Gallicola |
| 39 | FALSE | Bacteria | Firmicutes | Bacilli | Lactobacillales | Lactobacillaceae | NA |
| 38 | FALSE | Bacteria | Proteobacteria | Alphaproteobacteria | Sphingomonadales | Sphingomonadaceae | NA |
| 38 | FALSE | Bacteria | Bacteroidetes | Bacteroidia | Sphingobacteriales | Sphingobacteriaceae | Sphingobacterium |
| 37 | FALSE | Bacteria | Firmicutes | Clostridia | Clostridiales | Family_XI | Murdochiella |
| 37 | FALSE | Bacteria | Firmicutes | Clostridia | Clostridiales | Ruminococcaceae | Ruminococcaceae_UCG-009 |
| 37 | FALSE | Bacteria | Firmicutes | Clostridia | Clostridiales | Ruminococcaceae | Ruminococcaceae_UCG-011 |
| 37 | FALSE | Bacteria | Actinobacteria | Actinobacteria | Bifidobacteriales | Bifidobacteriaceae | Scardovia |
| 36 | FALSE | Bacteria | Proteobacteria | Alphaproteobacteria | Sphingomonadales | Sphingomonadaceae | Sphingopyxis |
| 35 | FALSE | Bacteria | Proteobacteria | Gammaproteobacteria | Betaproteobacteriales | Burkholderiaceae | Achromobacter |
| 35 | FALSE | Bacteria | Firmicutes | Clostridia | Clostridiales | Peptostreptococcaceae | Clostridioides |
| 35 | FALSE | Bacteria | Actinobacteria | Actinobacteria | Micrococcales | Dermabacteraceae | Dermabacter |
| 35 | FALSE | Bacteria | Synergistetes | Synergistia | Synergistales | Synergistaceae | Jonquetella |
| 34 | FALSE | Bacteria | Bacteroidetes | Bacteroidia | Flavobacteriales | Weeksellaceae | Chryseobacterium |
| 34 | FALSE | Bacteria | Firmicutes | Clostridia | Clostridiales | Ruminococcaceae | Ruminococcaceae_UCG-008 |
| 33 | FALSE | Bacteria | Tenericutes | Mollicutes | Anaeroplasmatales | Anaeroplasmataceae | Anaeroplasma |
| 32 | FALSE | Bacteria | Firmicutes | Clostridia | Clostridiales | Ruminococcaceae | Anaerotruncus |
| 32 | FALSE | Bacteria | Firmicutes | Clostridia | Clostridiales | Family_XI | Finegoldia |
| 32 | FALSE | Bacteria | Proteobacteria | Gammaproteobacteria | Enterobacteriales | Enterobacteriaceae | Hafnia-Obesumbacterium |
| 31 | FALSE | Bacteria | Firmicutes | Bacilli | Lactobacillales | Carnobacteriaceae | Carnobacterium |
| 31 | FALSE | Bacteria | Firmicutes | Clostridia | Clostridiales | Lachnospiraceae | Howardella |
| 31 | FALSE | Bacteria | Firmicutes | Clostridia | Clostridiales | Lachnospiraceae | Hungatella |

| | | | | | | | |
|---|---|---|---|---|---|---|---|
| 30 | FALSE | Bacteria | Actinobacteria | Coriobacteriia | Coriobacteriales | Coriobacteriales_Incertae_Sedis | Raoultibacter |
| 30 | FALSE | Bacteria | Firmicutes | Clostridia | Clostridiales | Ruminococcaceae | UBA1819 |
| 29 | FALSE | Bacteria | Firmicutes | Clostridia | Clostridiales | Lachnospiraceae | Anaerosporobacter |
| 29 | FALSE | Bacteria | Proteobacteria | Gammaproteobacteria | Enterobacteriales | Enterobacteriaceae | Citrobacter |
| 29 | FALSE | Bacteria | Actinobacteria | Coriobacteriia | Coriobacteriales | Eggerthellaceae | Cryptobacterium |
| 29 | FALSE | Bacteria | Actinobacteria | Acidimicrobiia | Microtrichales | NA | NA |
| 29 | FALSE | Bacteria | Actinobacteria | Actinobacteria | Corynebacteriales | Nocardiaceae | Rhodococcus |
| 29 | FALSE | Bacteria | Proteobacteria | Alphaproteobacteria | Acetobacterales | Acetobacteraceae | Roseomonas |
| 28 | FALSE | Bacteria | Firmicutes | Clostridia | Clostridiales | Lachnospiraceae | 28-4 |
| 28 | FALSE | Bacteria | Firmicutes | Clostridia | Clostridiales | Lachnospiraceae | Acetatifactor |
| 28 | FALSE | Bacteria | Actinobacteria | Actinobacteria | Propionibacteriales | Propionibacteriaceae | Acidipropionibacterium |
| 28 | FALSE | Bacteria | Proteobacteria | Gammaproteobacteria | Betaproteobacteriales | Burkholderiaceae | Acidovorax |
| 28 | FALSE | Bacteria | Firmicutes | Bacilli | Bacillales | Paenibacillaceae | Ammoniphilus |
| 28 | FALSE | Bacteria | Proteobacteria | Alphaproteobacteria | Caulobacterales | Caulobacteraceae | Brevundimonas |
| 28 | FALSE | Bacteria | Firmicutes | Bacilli | Bacillales | Paenibacillaceae | Cohnella |
| 28 | FALSE | Bacteria | Firmicutes | Bacilli | Bacillales | Planococcaceae | Domibacillus |
| 28 | FALSE | Bacteria | Firmicutes | Bacilli | Lactobacillales | Aerococcaceae | Facklamia |
| 28 | FALSE | Bacteria | Proteobacteria | Gammaproteobacteria | Betaproteobacteriales | Burkholderiaceae | Kerstersia |
| 28 | FALSE | Bacteria | Firmicutes | Clostridia | Clostridiales | Lachnospiraceae | Lachnospiraceae_UCG-006 |
| 28 | FALSE | Bacteria | Actinobacteria | Actinobacteria | Micromonosporales | Micromonosporaceae | Micromonospora |
| 28 | FALSE | Bacteria | Bacteroidetes | Bacteroidia | Bacteroidales | Muribaculaceae | Muribaculum |
| 28 | FALSE | Bacteria | Actinobacteria | Coriobacteriia | Coriobacteriales | NA | NA |
| 28 | FALSE | Bacteria | Proteobacteria | Alphaproteobacteria | Acetobacterales | Acetobacteraceae | NA |
| 28 | FALSE | Bacteria | Actinobacteria | Coriobacteriia | Coriobacteriales | Atopobiaceae | NA |
| 28 | FALSE | Bacteria | Firmicutes | Bacilli | NA | NA | NA |
| 28 | FALSE | Bacteria | Verrucomicrobia | Verrucomicrobiae | Opitutales | NA | NA |
| 28 | FALSE | Bacteria | Cyanobacteria | Oxyphotobacteria | Phormidesmiales | Nodosilineaceae | NA |
| 28 | FALSE | Bacteria | Firmicutes | Negativicutes | Selenomonadales | NA | NA |
| 28 | FALSE | Bacteria | Actinobacteria | Actinobacteria | Corynebacteriales | NA | NA |
| 28 | FALSE | Bacteria | Acidobacteria | FFCH5909 | NA | NA | NA |
| 28 | FALSE | Bacteria | Actinobacteria | Actinobacteria | Propionibacteriales | Nocardioidaceae | Nocardioides |
| 28 | FALSE | Bacteria | Firmicutes | Clostridia | Clostridiales | Peptostreptococcaceae | Paraclostridium |
| 28 | FALSE | Bacteria | Proteobacteria | Alphaproteobacteria | Rhizobiales | Beijerinckiaceae | Psychroglaciecola |
| 28 | FALSE | Bacteria | Proteobacteria | Gammaproteobacteria | Xanthomonadales | Xanthomonadaceae | SN8 |
| 28 | FALSE | Bacteria | Firmicutes | Bacilli | Bacillales | Bacillaceae | Terribacillus |

| | | | | | | | |
|---|---|---|---|---|---|---|---|
| 28 | FALSE | Bacteria | Actinobacteria | Actinobacteria | Actinomycetales | Actinomycetaceae | Trueperella |
| 27 | FALSE | Bacteria | Proteobacteria | Gammaproteobacteria | Aeromonadales | Aeromonadaceae | Aeromonas |
| 27 | FALSE | Bacteria | Proteobacteria | Gammaproteobacteria | Oceanospirillales | Alcanivoracaceae | Alcanivorax |
| 27 | FALSE | Bacteria | Firmicutes | Erysipelotrichia | Erysipelotrichales | Erysipelotrichaceae | Allobaculum |
| 27 | FALSE | Bacteria | Firmicutes | Bacilli | Lactobacillales | Carnobacteriaceae | Alloiococcus |
| 27 | FALSE | Bacteria | Proteobacteria | Alphaproteobacteria | Rhizobiales | Rhizobiaceae | Aminobacter |
| 27 | FALSE | Bacteria | Proteobacteria | Gammaproteobacteria | Aeromonadales | Succinivibrionaceae | Anaerobiospirillum |
| 27 | FALSE | Bacteria | Firmicutes | Clostridia | Clostridiales | Lachnospiraceae | Anaerocolumna |
| 27 | FALSE | Bacteria | Actinobacteria | Actinobacteria | Actinomycetales | Actinomycetaceae | Arcanobacterium |
| 27 | FALSE | Bacteria | Proteobacteria | Gammaproteobacteria | Enterobacteriales | Enterobacteriaceae | ATCC-39006 |
| 27 | FALSE | Bacteria | Proteobacteria | Alphaproteobacteria | Rhizobiales | Rhizobiaceae | Aureimonas |
| 27 | FALSE | Bacteria | Proteobacteria | Alphaproteobacteria | Sphingomonadales | Sphingomonadaceae | Blastomonas |
| 27 | FALSE | Bacteria | Spirochaetes | Brachyspirae | Brachyspirales | Brachyspiraceae | Brachyspira |
| 27 | FALSE | Bacteria | Firmicutes | Clostridia | Clostridiales | Caldicoprobacteraceae | Caldicoprobacter |
| 27 | FALSE | Bacteria | Proteobacteria | Alphaproteobacteria | Caedibacterales | Caedibacteraceae | Candidatus_Nucleicultrix |
| 27 | FALSE | Bacteria | Firmicutes | Negativicutes | Selenomonadales | Veillonellaceae | Centipeda |
| 27 | FALSE | Bacteria | Firmicutes | Clostridia | Clostridiales | Christensenellaceae | Christensenella |
| 27 | FALSE | Bacteria | Cyanobacteria | Oxyphotobacteria | Nostocales | Chroococcidiopsaceae | Chroococcidiopsis_SAG_2023 |
| 27 | FALSE | Bacteria | Bacteroidetes | Bacteroidia | Flavobacteriales | Weeksellaceae | Cloacibacterium |
| 27 | FALSE | Bacteria | Proteobacteria | Gammaproteobacteria | Enterobacteriales | Enterobacteriaceae | Cosenzaea |
| 27 | FALSE | Bacteria | Proteobacteria | Gammaproteobacteria | Betaproteobacteriales | Burkholderiaceae | Cupriavidus |
| 27 | FALSE | Bacteria | Deinococcus-Thermus | Deinococci | Deinococcales | Deinococcaceae | Deinococcus |
| 27 | FALSE | Bacteria | Actinobacteria | Coriobacteriia | Coriobacteriales | Eggerthellaceae | Denitrobacterium |
| 27 | FALSE | Bacteria | Firmicutes | Clostridia | Clostridiales | Peptococcaceae | Desulfitibacter |
| 27 | FALSE | Bacteria | Proteobacteria | Deltaproteobacteria | Desulfobacterales | Desulfobulbaceae | Desulfobulbus |
| 27 | FALSE | Bacteria | Proteobacteria | Alphaproteobacteria | Rhizobiales | Devosiaceae | Devosia |
| 27 | FALSE | Bacteria | Proteobacteria | Gammaproteobacteria | Xanthomonadales | Rhodanobacteraceae | Dokdonella |
| 27 | FALSE | Bacteria | Bacteroidetes | Bacteroidia | Cytophagales | Spirosomaceae | Dyadobacter |
| 27 | FALSE | Bacteria | Proteobacteria | Gammaproteobacteria | Pseudomonadales | Moraxellaceae | Enhydrobacter |
| 27 | FALSE | Bacteria | Firmicutes | Erysipelotrichia | Erysipelotrichales | Erysipelotrichaceae | Faecalicoccus |
| 27 | FALSE | Bacteria | Firmicutes | Bacilli | Bacillales | Bacillaceae | Fictibacillus |
| 27 | FALSE | Bacteria | Chloroflexi | Anaerolineae | Anaerolineales | Anaerolineaceae | Flexilinea |
| 27 | FALSE | Bacteria | Synergistetes | Synergistia | Synergistales | Synergistaceae | Fretibacterium |
| 27 | FALSE | Bacteria | Firmicutes | Bacilli | Lactobacillales | Aerococcaceae | Globicatella |

| | | | | | | | |
|---|---|---|---|---|---|---|---|
| 27 | FALSE | Bacteria | Actinobacteria | Actinobacteria | Micrococcales | Micrococcaceae | Glutamicibacter |
| 27 | FALSE | Bacteria | Proteobacteria | Alphaproteobacteria | Rhodobacterales | Rhodobacteraceae | Haematobacter |
| 27 | FALSE | Bacteria | Proteobacteria | Gammaproteobacteria | Oceanospirillales | Halomonadaceae | Halomonas |
| 27 | FALSE | Bacteria | Firmicutes | Clostridia | Clostridiales | Family_XI | Helcococcus |
| 27 | FALSE | Bacteria | Proteobacteria | Gammaproteobacteria | Betaproteobacteriales | Burkholderiaceae | Hydrogenophaga |
| 27 | FALSE | Bacteria | Proteobacteria | Alphaproteobacteria | Rhizobiales | Hyphomicrobiaceae | Hyphomicrobium |
| 27 | FALSE | Bacteria | Firmicutes | Bacilli | Lactobacillales | Aerococcaceae | Ignavigranum |
| 27 | FALSE | Bacteria | Actinobacteria | Actinobacteria | Micrococcales | Intrasporangiaceae | Janibacter |
| 27 | FALSE | Bacteria | Firmicutes | Clostridia | Clostridiales | Lachnospiraceae | Johnsonella |
| 27 | FALSE | Bacteria | Firmicutes | Clostridia | Clostridiales | Lachnospiraceae | Lachnoanaerobaculum |
| 27 | FALSE | Bacteria | Bacteroidetes | Bacteroidia | Chitinophagales | Chitinophagaceae | Lacibacter |
| 27 | FALSE | Bacteria | Actinobacteria | Actinobacteria | Micrococcales | Microbacteriaceae | Leucobacter |
| 27 | FALSE | Bacteria | Proteobacteria | Gammaproteobacteria | Betaproteobacteriales | Burkholderiaceae | Massilia |
| 27 | FALSE | Bacteria | Firmicutes | Erysipelotrichia | Erysipelotrichales | Erysipelotrichaceae | Merdibacter |
| 27 | FALSE | Bacteria | Actinobacteria | Actinobacteria | Propionibacteriales | Propionibacteriaceae | Micropruina |
| 27 | FALSE | Bacteria | Proteobacteria | Gammaproteobacteria | Pseudomonadales | Moraxellaceae | Moraxella |
| 27 | FALSE | Bacteria | Actinobacteria | Actinobacteria | Corynebacteriales | Mycobacteriaceae | Mycobacterium |
| 27 | FALSE | Bacteria | Firmicutes | Clostridia | Clostridiales | Family_XI | NA |
| 27 | FALSE | Bacteria | Firmicutes | Bacilli | Bacillales | Paenibacillaceae | NA |
| 27 | FALSE | Bacteria | Proteobacteria | Gammaproteobacteria | Betaproteobacteriales | NA | NA |
| 27 | FALSE | Bacteria | Actinobacteria | Coriobacteriia | Coriobacteriales | Coriobacteriaceae | NA |
| 27 | FALSE | Bacteria | Firmicutes | Clostridia | Clostridiales | Syntrophomonadaceae | NA |
| 27 | FALSE | Bacteria | Chloroflexi | Chloroflexia | Thermomicrobiales | JG30-KF-CM45 | NA |
| 27 | FALSE | Bacteria | Actinobacteria | Actinobacteria | Propionibacteriales | Propionibacteriaceae | NA |
| 27 | FALSE | Bacteria | Verrucomicrobia | Verrucomicrobiae | NA | NA | NA |
| 27 | FALSE | Bacteria | Actinobacteria | Actinobacteria | Micromonosporales | Micromonosporaceae | NA |
| 27 | FALSE | Bacteria | Actinobacteria | Actinobacteria | Streptomycetales | Streptomycetaceae | NA |
| 27 | FALSE | Bacteria | Firmicutes | Bacilli | Lactobacillales | Streptococcaceae | NA |
| 27 | FALSE | Bacteria | Proteobacteria | Alphaproteobacteria | NA | NA | NA |
| 27 | FALSE | Archaea | Euryarchaeota | Methanobacteria | Methanobacteriales | Methanobacteriaceae | NA |
| 27 | FALSE | Bacteria | Actinobacteria | Actinobacteria | NA | NA | NA |
| 27 | FALSE | Bacteria | Proteobacteria | Alphaproteobacteria | Micavibrionales | NA | NA |
| 27 | FALSE | Bacteria | Actinobacteria | Actinobacteria | Micrococcales | NA | NA |
| 27 | FALSE | Bacteria | Chloroflexi | Chloroflexia | Kallotenuales | NA | NA |
| 27 | FALSE | Bacteria | Tenericutes | Mollicutes | NA | NA | NA |

| | | | | | | | |
|---|---|---|---|---|---|---|---|
| 27 | FALSE | Bacteria | Proteobacteria | Gammaproteobacteria | Betaproteobacteriales | Rhodocyclaceae | NA |
| 27 | FALSE | Bacteria | Proteobacteria | Gammaproteobacteria | Betaproteobacteriales | Neisseriaceae | NA |
| 27 | FALSE | Eukaryota | NA | NA | NA | NA | NA |
| 27 | FALSE | Bacteria | Verrucomicrobia | Verrucomicrobiae | Verrucomicrobiales | NA | NA |
| 27 | FALSE | Bacteria | Actinobacteria | Actinobacteria | Micrococcales | Micrococcaceae | NA |
| 27 | FALSE | Bacteria | Bacteroidetes | Bacteroidia | Flavobacteriales | Weeksellaceae | NA |
| 27 | FALSE | Bacteria | Proteobacteria | Gammaproteobacteria | Betaproteobacteriales | Neisseriaceae | Neisseria |
| 27 | FALSE | Bacteria | Actinobacteria | Actinobacteria | Streptosporangiales | Nocardiopsaceae | Nocardiopsis |
| 27 | FALSE | Bacteria | Proteobacteria | Alphaproteobacteria | Sphingomonadales | Sphingomonadaceae | Novosphingobium |
| 27 | FALSE | Bacteria | Bacteroidetes | Bacteroidia | Sphingobacteriales | Sphingobacteriaceae | Nubsella |
| 27 | FALSE | Bacteria | Bacteroidetes | Bacteroidia | Sphingobacteriales | Sphingobacteriaceae | Pedobacter |
| 27 | FALSE | Bacteria | Proteobacteria | Alphaproteobacteria | Caulobacterales | Caulobacteraceae | Phenylobacterium |
| 27 | FALSE | Bacteria | Proteobacteria | Alphaproteobacteria | Rhizobiales | Rhizobiaceae | Phyllobacterium |
| 27 | FALSE | Bacteria | Proteobacteria | Gammaproteobacteria | Betaproteobacteriales | Burkholderiaceae | Pigmentiphaga |
| 27 | FALSE | Bacteria | Planctomycetes | Planctomycetacia | Pirellulales | Pirellulaceae | Pirellula |
| 27 | FALSE | Bacteria | Firmicutes | Clostridia | Clostridiales | Eubacteriaceae | Pseudoramibacter |
| 27 | FALSE | Bacteria | Firmicutes | Bacilli | Bacillales | Planococcaceae | Psychrobacillus |
| 27 | FALSE | Bacteria | Proteobacteria | Gammaproteobacteria | Betaproteobacteriales | Burkholderiaceae | Pusillimonas |
| 27 | FALSE | Bacteria | Firmicutes | Clostridia | Clostridiales | Ruminococcaceae | Pygmaiobacter |
| 27 | FALSE | Bacteria | Proteobacteria | Gammaproteobacteria | Betaproteobacteriales | Burkholderiaceae | Ralstonia |
| 27 | FALSE | Bacteria | Proteobacteria | Alphaproteobacteria | Reyranellales | Reyranellaceae | Reyranella |
| 27 | FALSE | Bacteria | Firmicutes | Bacilli | Bacillales | Planococcaceae | Rummeliibacillus |
| 27 | FALSE | Bacteria | Firmicutes | Negativicutes | Selenomonadales | Veillonellaceae | Selenomonas |
| 27 | FALSE | Bacteria | Firmicutes | Bacilli | Bacillales | Planococcaceae | Solibacillus |
| 27 | FALSE | Bacteria | Proteobacteria | Alphaproteobacteria | Sphingomonadales | Sphingomonadaceae | Sphingomonas |
| 27 | FALSE | Bacteria | Bacteroidetes | Bacteroidia | Cytophagales | Spirosomaceae | Spirosoma |
| 27 | FALSE | Bacteria | Firmicutes | Clostridia | Clostridiales | Lachnospiraceae | Stomatobaculum |
| 27 | FALSE | Bacteria | Proteobacteria | Gammaproteobacteria | Aeromonadales | Succinivibrionaceae | Succinivibrio |
| 27 | FALSE | Bacteria | Firmicutes | Bacilli | Lactobacillales | Enterococcaceae | Tetragenococcus |
| 27 | FALSE | Bacteria | Firmicutes | Clostridia | Clostridiales | Family_XI | Tissierella |
| 27 | FALSE | Bacteria | Spirochaetes | Spirochaetia | Spirochaetales | Spirochaetaceae | Treponema_2 |
| 27 | FALSE | Bacteria | Tenericutes | Mollicutes | Mycoplasmatales | Mycoplasmataceae | Ureaplasma |
| 27 | FALSE | Bacteria | Proteobacteria | Gammaproteobacteria | Cardiobacteriales | Wohlfahrtiimonadaceae | Wohlfahrtiimonas |
| 27 | FALSE | Bacteria | Proteobacteria | Alphaproteobacteria | Rhizobiales | Xanthobacteraceae | Xanthobacter |
| 27 | FALSE | Bacteria | Firmicutes | Clostridia | Clostridiales | Lachnospiraceae | XBB1006 |

| | | | | | | | |
|---|---|---|---|---|---|---|---|
| 27 | FALSE | Bacteria | Proteobacteria | Gammaproteobacteria | Betaproteobacteriales | Rhodocyclaceae | Zoogloea |
| 26 | FALSE | Bacteria | Firmicutes | Bacilli | Lactobacillales | Aerococcaceae | Abiotrophia |
| 26 | FALSE | Bacteria | Proteobacteria | Gammaproteobacteria | Gammaproteobacteria_Incertae_Sedis | Unknown_Family | Acidibacter |
| 26 | FALSE | Bacteria | Actinobacteria | Actinobacteria | Streptosporangiales | Thermomonosporaceae | Actinomadura |
| 26 | FALSE | Bacteria | Actinobacteria | Actinobacteria | Actinomycetales | Actinomycetaceae | Actinomyces |
| 26 | FALSE | Bacteria | Actinobacteria | Actinobacteria | Propionibacteriales | Nocardioidaceae | Aeromicrobium |
| 26 | FALSE | Bacteria | Proteobacteria | Gammaproteobacteria | Pasteurellales | Pasteurellaceae | Aggregatibacter |
| 26 | FALSE | Bacteria | Proteobacteria | Gammaproteobacteria | Betaproteobacteriales | Burkholderiaceae | Alcaligenes |
| 26 | FALSE | Bacteria | Firmicutes | Bacilli | Lactobacillales | Carnobacteriaceae | Allofustis |
| 26 | FALSE | Bacteria | Proteobacteria | Alphaproteobacteria | Sphingomonadales | Sphingomonadaceae | Altererythrobacter |
| 26 | FALSE | Bacteria | Firmicutes | Bacilli | Bacillales | Bacillaceae | Anaerobacillus |
| 26 | FALSE | Bacteria | Firmicutes | Bacilli | Bacillales | Paenibacillaceae | Aneurinibacillus |
| 26 | FALSE | Bacteria | Epsilonbacteraeota | Campylobacteria | Campylobacterales | Arcobacteraceae | Arcobacter |
| 26 | FALSE | Bacteria | Actinobacteria | Coriobacteriia | Coriobacteriales | Atopobiaceae | Atopobium |
| 26 | FALSE | Bacteria | Bacteroidetes | Bacteroidia | Flavobacteriales | Weeksellaceae | Bergeyella |
| 26 | FALSE | Bacteria | Proteobacteria | Alphaproteobacteria | Rhizobiales | Xanthobacteraceae | Bradyrhizobium |
| 26 | FALSE | Bacteria | Bacteroidetes | Bacteroidia | Cytophagales | Amoebophilaceae | Candidatus_Amoebophilus |
| 26 | FALSE | Bacteria | Bacteroidetes | Bacteroidia | Flavobacteriales | Flavobacteriaceae | Capnocytophaga |
| 26 | FALSE | Bacteria | Proteobacteria | Gammaproteobacteria | Cardiobacteriales | Cardiobacteriaceae | Cardiobacterium |
| 26 | FALSE | Bacteria | Firmicutes | Erysipelotrichia | Erysipelotrichales | Erysipelotrichaceae | Catenisphaera |
| 26 | FALSE | Bacteria | Actinobacteria | Actinobacteria | Micrococcales | Cellulomonadaceae | Cellulomonas |
| 26 | FALSE | Bacteria | Firmicutes | Clostridia | Clostridiales | Lachnospiraceae | Cellulosilyticum |
| 26 | FALSE | Bacteria | Fusobacteria | Fusobacteriia | Fusobacteriales | Fusobacteriaceae | Cetobacterium |
| 26 | FALSE | Bacteria | Actinobacteria | Coriobacteriia | Coriobacteriales | Eggerthellaceae | CHKCI002 |
| 26 | FALSE | Bacteria | Firmicutes | Clostridia | Clostridiales | Clostridiaceae_1 | Clostridium_sensu_stricto_11 |
| 26 | FALSE | Bacteria | Firmicutes | Clostridia | Clostridiales | Clostridiaceae_1 | Clostridium_sensu_stricto_7 |
| 26 | FALSE | Bacteria | Actinobacteria | Coriobacteriia | Coriobacteriales | Atopobiaceae | Coriobacteriaceae_UCG-003 |
| 26 | FALSE | Bacteria | Actinobacteria | Actinobacteria | Corynebacteriales | Dietziaceae | Dietzia |
| 26 | FALSE | Bacteria | Proteobacteria | Gammaproteobacteria | Betaproteobacteriales | Neisseriaceae | Eikenella |
| 26 | FALSE | Bacteria | Proteobacteria | Alphaproteobacteria | Rhizobiales | Rhizobiaceae | Ensifer |
| 26 | FALSE | Bacteria | Actinobacteria | Actinobacteria | Actinomycetales | Actinomycetaceae | F0332 |
| 26 | FALSE | Bacteria | Bacteroidetes | Bacteroidia | Cytophagales | Spirosomaceae | Flectobacillus |
| 26 | FALSE | Bacteria | Firmicutes | Bacilli | Bacillales | Bacillaceae | Geobacillus |
| 26 | FALSE | Bacteria | Firmicutes | Bacilli | Bacillales | Bacillaceae | Gracilibacillus |
| 26 | FALSE | Bacteria | Firmicutes | Clostridia | Clostridiales | Clostridiaceae_1 | Hathewaya |

| | | | | | | | |
|---|---|---|---|---|---|---|---|
| 26 | FALSE | Bacteria | Firmicutes | Bacilli | Bacillales | Staphylococcaceae | Jeotgalicoccus |
| 26 | FALSE | Bacteria | Firmicutes | Bacilli | Bacillales | Thermoactinomycetaceae | Kroppenstedtia |
| 26 | FALSE | Bacteria | Verrucomicrobia | Verrucomicrobiae | Chthoniobacterales | Chthoniobacteraceae | LD29 |
| 26 | FALSE | Bacteria | Actinobacteria | Actinobacteria | Micrococcales | Microbacteriaceae | Leifsonia |
| 26 | FALSE | Bacteria | Fusobacteria | Fusobacteriia | Fusobacteriales | Leptotrichiaceae | Leptotrichia |
| 26 | FALSE | Bacteria | Actinobacteria | Coriobacteriia | Coriobacteriales | Atopobiaceae | Libanicoccus |
| 26 | FALSE | Bacteria | Firmicutes | Bacilli | Bacillales | Staphylococcaceae | Macrococcus |
| 26 | FALSE | Bacteria | Proteobacteria | Alphaproteobacteria | Rhizobiales | Rhizobiaceae | Mesorhizobium |
| 26 | FALSE | Bacteria | Tenericutes | Mollicutes | Mycoplasmatales | Mycoplasmataceae | Mycoplasma |
| 26 | FALSE | NA | NA | NA | NA | NA | NA |
| 26 | FALSE | Bacteria | Proteobacteria | Gammaproteobacteria | Betaproteobacteriales | Burkholderiaceae | NA |
| 26 | FALSE | Bacteria | Firmicutes | Bacilli | Lactobacillales | Enterococcaceae | NA |
| 26 | FALSE | Bacteria | Actinobacteria | Actinobacteria | Corynebacteriales | Nocardiaceae | NA |
| 26 | FALSE | Bacteria | Proteobacteria | Alphaproteobacteria | Rickettsiales | Mitochondria | NA |
| 26 | FALSE | Bacteria | Actinobacteria | Actinobacteria | Micrococcales | Microbacteriaceae | NA |
| 26 | FALSE | Bacteria | Actinobacteria | Actinobacteria | Bifidobacteriales | Bifidobacteriaceae | NA |
| 26 | FALSE | Bacteria | Bacteroidetes | Bacteroidia | Bacteroidales | Dysgonomonadaceae | NA |
| 26 | FALSE | Bacteria | Actinobacteria | NA | NA | NA | NA |
| 26 | FALSE | Bacteria | Verrucomicrobia | Verrucomicrobiae | Opitutales | Puniceicoccaceae | NA |
| 26 | FALSE | Bacteria | Proteobacteria | Alphaproteobacteria | Rhizobiales | Xanthobacteraceae | NA |
| 26 | FALSE | Bacteria | Firmicutes | Bacilli | Bacillales | Staphylococcaceae | NA |
| 26 | FALSE | Bacteria | Spirochaetes | Spirochaetia | Spirochaetales | Spirochaetaceae | NA |
| 26 | FALSE | Bacteria | Firmicutes | Bacilli | Lactobacillales | Carnobacteriaceae | NA |
| 26 | FALSE | Bacteria | Bacteroidetes | Bacteroidia | Sphingobacteriales | Sphingobacteriaceae | NA |
| 26 | FALSE | Bacteria | Proteobacteria | Gammaproteobacteria | NA | NA | NA |
| 26 | FALSE | Bacteria | Kiritimatiellaeota | Kiritimatiellae | WCHB1-41 | NA | NA |
| 26 | FALSE | Bacteria | Synergistetes | Synergistia | Synergistales | Synergistaceae | NA |
| 26 | FALSE | Bacteria | Bacteroidetes | Bacteroidia | Bacteroidales | Tannerellaceae | NA |
| 26 | FALSE | Bacteria | Proteobacteria | Alphaproteobacteria | Rhizobiales | Rhizobiaceae | Neorhizobium |
| 26 | FALSE | Bacteria | Firmicutes | Clostridia | Clostridiales | Ruminococcaceae | Papillibacter |
| 26 | FALSE | Bacteria | Actinobacteria | Coriobacteriia | Coriobacteriales | Eggerthellaceae | Paraeggerthella |
| 26 | FALSE | Bacteria | Firmicutes | Bacilli | Bacillales | Bacillaceae | Paucisalibacillus |
| 26 | FALSE | Bacteria | Firmicutes | Negativicutes | Selenomonadales | Veillonellaceae | Pectinatus |
| 26 | FALSE | Bacteria | Proteobacteria | Gammaproteobacteria | Betaproteobacteriales | Burkholderiaceae | Pelomonas |
| 26 | FALSE | Bacteria | Proteobacteria | Gammaproteobacteria | Enterobacteriales | Enterobacteriaceae | Plesiomonas |

| | | | | | | | |
|---|---|---|---|---|---|---|---|
| 26 | FALSE | Bacteria | Proteobacteria | Gammaproteobacteria | Enterobacteriales | Enterobacteriaceae | Pluralibacter |
| 26 | FALSE | Bacteria | Bacteroidetes | Bacteroidia | Bacteroidales | Prevotellaceae | Prevotellaceae_Ga6A1_group |
| 26 | FALSE | Bacteria | Actinobacteria | Actinobacteria | Propionibacteriales | Propionibacteriaceae | Propioniferax |
| 26 | FALSE | Bacteria | Proteobacteria | Alphaproteobacteria | Rhizobiales | Rhizobiaceae | Pseudochrobactrum |
| 26 | FALSE | Bacteria | Firmicutes | Clostridia | Clostridiales | Ruminococcaceae | Pseudoflavonifractor |
| 26 | FALSE | Bacteria | Actinobacteria | Actinobacteria | Propionibacteriales | Propionibacteriaceae | Pseudopropionibacterium |
| 26 | FALSE | Bacteria | Proteobacteria | Gammaproteobacteria | Betaproteobacteriales | Burkholderiaceae | Pseudorhodoferax |
| 26 | FALSE | Bacteria | Firmicutes | Clostridia | Clostridiales | Lachnospiraceae | Robinsoniella |
| 26 | FALSE | Bacteria | Firmicutes | Negativicutes | Selenomonadales | Veillonellaceae | Selenomonas_4 |
| 26 | FALSE | Bacteria | Proteobacteria | Alphaproteobacteria | Rhizobiales | Rhizobiaceae | Shinella |
| 26 | FALSE | Bacteria | Proteobacteria | Alphaproteobacteria | Azospirillales | Azospirillaceae | Skermanella |
| 26 | FALSE | Bacteria | Spirochaetes | Spirochaetia | Spirochaetales | Spirochaetaceae | Sphaerochaeta |
| 26 | FALSE | Bacteria | Proteobacteria | Alphaproteobacteria | Sphingomonadales | Sphingomonadaceae | Sphingobium |
| 26 | FALSE | Bacteria | Firmicutes | Clostridia | Clostridiales | Lachnospiraceae | Sporobacterium |
| 26 | FALSE | Bacteria | Firmicutes | Bacilli | Bacillales | Planococcaceae | Sporosarcina |
| 26 | FALSE | Bacteria | Proteobacteria | Gammaproteobacteria | Aeromonadales | Succinivibrionaceae | Succinatimonas |
| 26 | FALSE | Bacteria | Firmicutes | Clostridia | Clostridiales | Syntrophomonadaceae | Syntrophomonas |
| 26 | FALSE | Bacteria | Proteobacteria | Gammaproteobacteria | Betaproteobacteriales | Burkholderiaceae | Tepidimonas |
| 26 | FALSE | Bacteria | Deinococcus-Thermus | Deinococci | Thermales | Thermaceae | Thermus |
| 26 | FALSE | Bacteria | Firmicutes | Bacilli | Lactobacillales | Enterococcaceae | Vagococcus |
| 26 | FALSE | Bacteria | Proteobacteria | Gammaproteobacteria | Betaproteobacteriales | Burkholderiaceae | Variovorax |
| 26 | FALSE | Bacteria | Firmicutes | Bacilli | Bacillales | Bacillaceae | Virgibacillus |
| 26 | FALSE | Bacteria | Firmicutes | Clostridia | Clostridiales | Family_XI | W5053 |
| 26 | FALSE | Bacteria | Proteobacteria | Gammaproteobacteria | Betaproteobacteriales | Burkholderiaceae | Xylophilus |
| 26 | FALSE | Bacteria | Proteobacteria | Gammaproteobacteria | Enterobacteriales | Enterobacteriaceae | Yersinia |
| 25 | FALSE | Bacteria | Actinobacteria | Actinobacteria | Actinomycetales | Actinomycetaceae | Actinotignum |
| 25 | FALSE | Bacteria | Proteobacteria | Alphaproteobacteria | Rhizobiales | Rhizobiaceae | Allorhizobium-Neorhizobium-Pararhizobium-Rhizobium |
| 25 | FALSE | Bacteria | Firmicutes | Clostridia | Clostridiales | Family_XIII | Anaerovorax |
| 25 | FALSE | Archaea | Euryarchaeota | Thermoplasmata | Methanomassiliicoccales | Methanomethylophilaceae | Candidatus_Methanomethylophilus |
| 25 | FALSE | Bacteria | Patescibacteria | Saccharimonadia | Saccharimonadales | Saccharimonadaceae | Candidatus_Saccharimonas |
| 25 | FALSE | Bacteria | Firmicutes | Clostridia | Clostridiales | Clostridiaceae_1 | Clostridium_sensu_stricto_2 |
| 25 | FALSE | Bacteria | Firmicutes | Bacilli | Lactobacillales | Aerococcaceae | Eremococcus |
| 25 | FALSE | Bacteria | Firmicutes | Erysipelotrichia | Erysipelotrichales | Erysipelotrichaceae | Erysipelotrichaceae_UCG-004 |
| 25 | FALSE | Bacteria | Firmicutes | Erysipelotrichia | Erysipelotrichales | Erysipelotrichaceae | Erysipelotrichaceae_UCG-006 |

| | | | | | | | |
|---|---|---|---|---|---|---|---|
| 25 | FALSE | Bacteria | Bacteroidetes | Bacteroidia | Flavobacteriales | Flavobacteriaceae | Flavobacterium |
| 25 | FALSE | Bacteria | Actinobacteria | Actinobacteria | Corynebacteriales | Nocardiaceae | Gordonia |
| 25 | FALSE | Bacteria | Proteobacteria | Gammaproteobacteria | Betaproteobacteriales | Burkholderiaceae | Herbaspirillum |
| 25 | FALSE | Bacteria | Firmicutes | Clostridia | Clostridiales | Lachnospiraceae | Lachnoclostridium_10 |
| 25 | FALSE | Bacteria | Proteobacteria | Gammaproteobacteria | Betaproteobacteriales | Burkholderiaceae | Lautropia |
| 25 | FALSE | Bacteria | Proteobacteria | Deltaproteobacteria | Desulfovibrionales | Desulfovibrionaceae | Mailhella |
| 25 | FALSE | Bacteria | Firmicutes | Negativicutes | Selenomonadales | Veillonellaceae | Mitsuokella |
| 25 | FALSE | Bacteria | Bacteroidetes | Bacteroidia | Bacteroidales | Rikenellaceae | NA |
| 25 | FALSE | Bacteria | Proteobacteria | NA | NA | NA | NA |
| 25 | FALSE | Bacteria | Lentisphaerae | Lentisphaeria | Victivallales | NA | NA |
| 25 | FALSE | Bacteria | Firmicutes | Bacilli | Lactobacillales | NA | NA |
| 25 | FALSE | Bacteria | Bacteroidetes | NA | NA | NA | NA |
| 25 | FALSE | Bacteria | Actinobacteria | Actinobacteria | Actinomycetales | Actinomycetaceae | NA |
| 25 | FALSE | Bacteria | Firmicutes | Negativicutes | Selenomonadales | Acidaminococcaceae | NA |
| 25 | FALSE | Bacteria | Firmicutes | Bacilli | Bacillales | Staphylococcaceae | Nosocomiicoccus |
| 25 | FALSE | Bacteria | Proteobacteria | Alphaproteobacteria | Rhizobiales | Rhizobiaceae | Ochrobactrum |
| 25 | FALSE | Bacteria | Firmicutes | Clostridia | Clostridiales | Lachnospiraceae | Oribacterium |
| 25 | FALSE | Bacteria | Firmicutes | Bacilli | Bacillales | Bacillaceae | Ornithinibacillus |
| 25 | FALSE | Bacteria | Proteobacteria | Alphaproteobacteria | Rhodobacterales | Rhodobacteraceae | Paracoccus |
| 25 | FALSE | Bacteria | Actinobacteria | Actinobacteria | Propionibacteriales | Propionibacteriaceae | Propionimicrobium |
| 25 | FALSE | Bacteria | Firmicutes | Clostridia | Clostridiales | Ruminococcaceae | Ruminococcaceae_V9D2013_group |
| 25 | FALSE | Bacteria | Firmicutes | Clostridia | Clostridiales | Clostridiaceae_1 | Sarcina |
| 25 | FALSE | Bacteria | Proteobacteria | Gammaproteobacteria | Enterobacteriales | Enterobacteriaceae | Shimwellia |
| 25 | FALSE | Bacteria | Fusobacteria | Fusobacteriia | Fusobacteriales | Leptotrichiaceae | Sneathia |
| 25 | FALSE | Bacteria | Synergistetes | Synergistia | Synergistales | Synergistaceae | Synergistes |
| 24 | FALSE | Bacteria | Actinobacteria | Actinobacteria | Actinomycetales | Actinomycetaceae | Actinobaculum |
| 24 | FALSE | Bacteria | Firmicutes | Bacilli | Lactobacillales | Aerococcaceae | Aerococcus |
| 24 | FALSE | Bacteria | Bacteroidetes | Bacteroidia | Bacteroidales | Prevotellaceae | Alloprevotella |
| 24 | FALSE | Bacteria | Actinobacteria | Actinobacteria | Bifidobacteriales | Bifidobacteriaceae | Alloscardovia |
| 24 | FALSE | Bacteria | Firmicutes | Clostridia | Clostridiales | Eubacteriaceae | Anaerofustis |
| 24 | FALSE | Bacteria | Firmicutes | Negativicutes | Selenomonadales | Veillonellaceae | Anaeroglobus |
| 24 | FALSE | Bacteria | Bacteroidetes | Bacteroidia | Bacteroidales | Muribaculaceae | CAG-873 |
| 24 | FALSE | Bacteria | Actinobacteria | Actinobacteria | Micrococcales | Promicromonosporaceae | Cellulosimicrobium |
| 24 | FALSE | Bacteria | Verrucomicrobia | Verrucomicrobiae | Opitutales | Puniceicoccaceae | Cerasicoccus |
| 24 | FALSE | Bacteria | Firmicutes | Clostridia | Clostridiales | Clostridiaceae_1 | Clostridium_sensu_stricto_3 |

| | | | | | | | |
|---|---|---|---|---|---|---|---|
| 24 | FALSE | Bacteria | Proteobacteria | Gammaproteobacteria | Betaproteobacteriales | Burkholderiaceae | Comamonas |
| 24 | FALSE | Bacteria | Actinobacteria | Coriobacteriia | Coriobacteriales | Eggerthellaceae | DNF00809 |
| 24 | FALSE | Bacteria | Actinobacteria | Coriobacteriia | Coriobacteriales | Coriobacteriaceae | Enorma |
| 24 | FALSE | Bacteria | Firmicutes | Clostridia | Clostridiales | Lachnospiraceae | Herbinix |
| 24 | FALSE | Bacteria | Proteobacteria | Gammaproteobacteria | Cardiobacteriales | Wohlfahrtiimonadaceae | Ignatzschineria |
| 24 | FALSE | Bacteria | Proteobacteria | Gammaproteobacteria | Enterobacteriales | Enterobacteriaceae | Kosakonia |
| 24 | FALSE | Bacteria | Actinobacteria | Actinobacteria | Micrococcales | Microbacteriaceae | Microbacterium |
| 24 | FALSE | Bacteria | Firmicutes | Clostridia | Clostridiales | Family_XIII | Mogibacterium |
| 24 | FALSE | Bacteria | Actinobacteria | Coriobacteriia | Coriobacteriales | Eggerthellaceae | NA |
| 24 | FALSE | Bacteria | Lentisphaerae | Lentisphaeria | Victivallales | Victivallaceae | NA |
| 24 | FALSE | Bacteria | Firmicutes | Bacilli | Lactobacillales | Aerococcaceae | NA |
| 24 | FALSE | Bacteria | Proteobacteria | Gammaproteobacteria | Aeromonadales | Succinivibrionaceae | NA |
| 24 | FALSE | Bacteria | Proteobacteria | Alphaproteobacteria | Rhodobacterales | Rhodobacteraceae | NA |
| 24 | FALSE | Bacteria | Firmicutes | Clostridia | Clostridiales | Peptostreptococcaceae | Paeniclostridium |
| 24 | FALSE | Bacteria | Bacteroidetes | Bacteroidia | Bacteroidales | Prevotellaceae | Prevotellaceae_UCG-001 |
| 24 | FALSE | Bacteria | Actinobacteria | Actinobacteria | Micrococcales | Micrococcaceae | Pseudoglutamicibacter |
| 24 | FALSE | Bacteria | Bacteroidetes | Bacteroidia | Bacteroidales | Rikenellaceae | Rikenella |
| 24 | FALSE | Bacteria | Firmicutes | Clostridia | Clostridiales | Ruminococcaceae | Ruminococcaceae_UCG-007 |
| 24 | FALSE | Bacteria | Firmicutes | Clostridia | Clostridiales | Family_XI | Sedimentibacter |
| 23 | FALSE | Bacteria | Firmicutes | Negativicutes | Selenomonadales | Veillonellaceae | Allisonella |
| 23 | FALSE | Bacteria | Bacteroidetes | Bacteroidia | Bacteroidales | Dysgonomonadaceae | Dysgonomonas |
| 23 | FALSE | Archaea | Euryarchaeota | Thermoplasmata | Methanomassiliicoccales | Methanomassiliicoccaceae | Methanomassiliicoccus |
| 23 | FALSE | Bacteria | Firmicutes | Clostridia | Clostridiales | Clostridiaceae_1 | NA |
| 23 | FALSE | Bacteria | Firmicutes | Clostridia | Clostridiales | Peptostreptococcaceae | Peptostreptococcus |
| 23 | FALSE | Bacteria | Bacteroidetes | Bacteroidia | Bacteroidales | Rikenellaceae | Rikenellaceae_RC9_gut_group |
| 23 | FALSE | Bacteria | Firmicutes | Bacilli | Lactobacillales | Streptococcaceae | Streptococcus |
| 22 | FALSE | Bacteria | Firmicutes | Erysipelotrichia | Erysipelotrichales | Erysipelotrichaceae | Dielma |
| 22 | FALSE | Bacteria | Fusobacteria | Fusobacteriia | Fusobacteriales | Fusobacteriaceae | Fusobacterium |
| 22 | FALSE | Bacteria | Firmicutes | Erysipelotrichia | Erysipelotrichales | Erysipelotrichaceae | Holdemanella |
| 22 | FALSE | Bacteria | Firmicutes | Clostridia | Clostridiales | Ruminococcaceae | NA |
| 22 | FALSE | Bacteria | Proteobacteria | Gammaproteobacteria | Pseudomonadales | Pseudomonadaceae | NA |
| 22 | FALSE | Bacteria | NA | NA | NA | NA | NA |
| 22 | FALSE | Bacteria | Lentisphaerae | Lentisphaeria | Victivallales | vadinBE97 | NA |
| 22 | FALSE | Bacteria | Bacteroidetes | Bacteroidia | Bacteroidales | Marinifilaceae | Sanguibacteroides |
| 22 | FALSE | Bacteria | Firmicutes | Clostridia | Clostridiales | Lachnospiraceae | Tyzzerella_4 |

| | | | | | | | |
|---|---|---|---|---|---|---|---|
| 21 | FALSE | Bacteria | Firmicutes | Clostridia | Clostridiales | Defluviitaleaceae | Defluviitaleaceae_UCG-011 |
| 21 | FALSE | Bacteria | Actinobacteria | Coriobacteriia | Coriobacteriales | Eggerthellaceae | Enterorhabdus |
| 21 | FALSE | Bacteria | Proteobacteria | Gammaproteobacteria | Pasteurellales | Pasteurellaceae | Haemophilus |
| 21 | FALSE | Bacteria | Proteobacteria | Gammaproteobacteria | Pasteurellales | Pasteurellaceae | NA |
| 21 | FALSE | Bacteria | Firmicutes | Negativicutes | Selenomonadales | Veillonellaceae | Negativicoccus |
| 21 | FALSE | Bacteria | Firmicutes | Clostridia | Clostridiales | Peptococcaceae | Peptococcus |
| 21 | FALSE | Bacteria | Proteobacteria | Gammaproteobacteria | Xanthomonadales | Xanthomonadaceae | Pseudoxanthomonas |
| 21 | FALSE | Bacteria | Proteobacteria | Gammaproteobacteria | Enterobacteriales | Enterobacteriaceae | Salmonella |
| 21 | FALSE | Bacteria | Actinobacteria | Coriobacteriia | Coriobacteriales | Eggerthellaceae | Senegalimassilia |
| 21 | FALSE | Bacteria | Actinobacteria | Actinobacteria | Streptomycetales | Streptomycetaceae | Streptomyces |
| 20 | FALSE | Bacteria | Actinobacteria | Actinobacteria | Micrococcales | Dermabacteraceae | Brachybacterium |
| 20 | FALSE | Bacteria | Firmicutes | Bacilli | Bacillales | Paenibacillaceae | Brevibacillus |
| 20 | FALSE | Bacteria | Firmicutes | Clostridia | Clostridiales | Lachnospiraceae | Eisenbergiella |
| 20 | FALSE | Bacteria | Firmicutes | Clostridia | Clostridiales | Lachnospiraceae | Moryella |
| 20 | FALSE | Archaea | Euryarchaeota | Thermoplasmata | Methanomassiliicoccales | Methanomethylophilaceae | NA |
| 20 | FALSE | Bacteria | Firmicutes | Clostridia | Clostridiales | Christensenellaceae | NA |
| 20 | FALSE | Bacteria | Proteobacteria | Gammaproteobacteria | Betaproteobacteriales | Burkholderiaceae | Oligella |
| 20 | FALSE | Bacteria | Proteobacteria | Gammaproteobacteria | Enterobacteriales | Enterobacteriaceae | Providencia |
| 20 | FALSE | Bacteria | Firmicutes | Clostridia | Clostridiales | Ruminococcaceae | Ruminiclostridium_1 |
| 20 | FALSE | Bacteria | Firmicutes | Erysipelotrichia | Erysipelotrichales | Erysipelotrichaceae | Solobacterium |
| 19 | FALSE | Bacteria | Actinobacteria | Actinobacteria | Micrococcales | Brevibacteriaceae | Brevibacterium |
| 19 | FALSE | Bacteria | Firmicutes | Clostridia | Clostridiales | Ruminococcaceae | CAG-352 |
| 19 | FALSE | Bacteria | Firmicutes | Clostridia | Clostridiales | Lachnospiraceae | Epulopiscium |
| 19 | FALSE | Bacteria | Firmicutes | Clostridia | Clostridiales | Ruminococcaceae | Hydrogenoanaerobacterium |
| 19 | FALSE | Bacteria | Bacteroidetes | Bacteroidia | Bacteroidales | Prevotellaceae | Prevotella_6 |
| 19 | FALSE | Bacteria | Synergistetes | Synergistia | Synergistales | Synergistaceae | Pyramidobacter |
| 18 | FALSE | Bacteria | Firmicutes | Erysipelotrichia | Erysipelotrichales | Erysipelotrichaceae | Catenibacterium |
| 18 | FALSE | Bacteria | Proteobacteria | Gammaproteobacteria | Enterobacteriales | Enterobacteriaceae | Edwardsiella |
| 18 | FALSE | Bacteria | Firmicutes | Clostridia | Clostridiales | Ruminococcaceae | GCA-900066225 |
| 18 | FALSE | Bacteria | Firmicutes | Bacilli | Bacillales | Planococcaceae | Lysinibacillus |
| 18 | FALSE | Bacteria | Actinobacteria | Actinobacteria | Micrococcales | Micrococcaceae | Micrococcus |
| 18 | FALSE | Bacteria | Bacteroidetes | Bacteroidia | Bacteroidales | Prevotellaceae | NA |
| 18 | FALSE | Bacteria | Bacteroidetes | Bacteroidia | Bacteroidales | Prevotellaceae | Prevotella_2 |
| 18 | FALSE | Bacteria | Firmicutes | Clostridia | Clostridiales | Lachnospiraceae | Sellimonas |
| 17 | FALSE | Bacteria | Bacteroidetes | Bacteroidia | Bacteroidales | Barnesiellaceae | Coprobacter |

| | | | | | | | |
|---|---|---|---|---|---|---|---|
| 17 | FALSE | Bacteria | Firmicutes | Negativicutes | Selenomonadales | Veillonellaceae | Megasphaera |
| 17 | FALSE | Bacteria | Actinobacteria | Actinobacteria | Corynebacteriales | Corynebacteriaceae | NA |
| 17 | FALSE | Bacteria | Bacteroidetes | Bacteroidia | Bacteroidales | Muribaculaceae | NA |
| 17 | FALSE | Bacteria | Firmicutes | Clostridia | DTU014 | NA | NA |
| 17 | FALSE | Bacteria | Firmicutes | Bacilli | Bacillales | Bacillaceae | Pseudogracilibacillus |
| 17 | FALSE | Bacteria | Firmicutes | Clostridia | Clostridiales | Peptostreptococcaceae | Terrisporobacter |
| 16 | FALSE | Bacteria | Firmicutes | Negativicutes | Selenomonadales | Acidaminococcaceae | Acidaminococcus |
| 16 | FALSE | Bacteria | Bacteroidetes | Bacteroidia | Bacteroidales | Marinifilaceae | Butyricimonas |
| 16 | FALSE | Bacteria | Firmicutes | Clostridia | Clostridiales | Lachnospiraceae | Butyrivibrio |
| 16 | FALSE | Bacteria | Actinobacteria | Actinobacteria | Bifidobacteriales | Bifidobacteriaceae | Gardnerella |
| 16 | FALSE | Bacteria | Firmicutes | Clostridia | Clostridiales | Peptococcaceae | NA |
| 15 | FALSE | Bacteria | Actinobacteria | Actinobacteria | Corynebacteriales | Corynebacteriaceae | Corynebacterium |
| 15 | FALSE | Bacteria | Firmicutes | Erysipelotrichia | Erysipelotrichales | Erysipelotrichaceae | Faecalitalea |
| 15 | FALSE | Bacteria | Actinobacteria | Actinobacteria | Micrococcales | Micrococcaceae | Kocuria |
| 15 | FALSE | Archaea | Euryarchaeota | Methanobacteria | Methanobacteriales | Methanobacteriaceae | Methanosphaera |
| 15 | FALSE | Bacteria | Firmicutes | Clostridia | Clostridiales | NA | NA |
| 15 | FALSE | Bacteria | Actinobacteria | Actinobacteria | Propionibacteriales | Propionibacteriaceae | Propionibacterium |
| 15 | FALSE | Bacteria | Firmicutes | Clostridia | Clostridiales | Ruminococcaceae | Ruminococcaceae_UCG-005 |
| 14 | FALSE | Bacteria | Firmicutes | Clostridia | Clostridiales | Ruminococcaceae | Acetanaerobacterium |
| 14 | FALSE | Bacteria | Firmicutes | Clostridia | Clostridiales | Ruminococcaceae | Anaerofilum |
| 14 | FALSE | Bacteria | Synergistetes | Synergistia | Synergistales | Synergistaceae | Cloacibacillus |
| 14 | FALSE | Bacteria | Firmicutes | Bacilli | Lactobacillales | Leuconostocaceae | Leuconostoc |
| 14 | FALSE | Bacteria | Firmicutes | Clostridia | NA | NA | NA |
| 14 | FALSE | Bacteria | Firmicutes | Clostridia | Clostridiales | Peptostreptococcaceae | Peptoclostridium |
| 14 | FALSE | Bacteria | Firmicutes | Clostridia | Clostridiales | Ruminococcaceae | Ruminiclostridium_5 |
| 14 | FALSE | Bacteria | Firmicutes | Negativicutes | Selenomonadales | Acidaminococcaceae | Succiniclasticum |
| 13 | FALSE | Bacteria | Proteobacteria | Alphaproteobacteria | Rhizobiales | Rhizobiaceae | Brucella |
| 13 | FALSE | Bacteria | Firmicutes | Clostridia | Clostridiales | Lachnospiraceae | Lactonifactor |
| 13 | FALSE | Bacteria | Bacteroidetes | Bacteroidia | Bacteroidales | Barnesiellaceae | NA |
| 13 | FALSE | Bacteria | Patescibacteria | Saccharimonadia | Saccharimonadales | NA | NA |
| 13 | FALSE | Bacteria | Tenericutes | Mollicutes | Mollicutes_RF39 | NA | NA |
| 13 | FALSE | Bacteria | Bacteroidetes | Bacteroidia | Bacteroidales | Prevotellaceae | Prevotellaceae_NK3B31_group |
| 13 | FALSE | Bacteria | Firmicutes | Clostridia | Clostridiales | Ruminococcaceae | Ruminiclostridium |
| 12 | FALSE | Bacteria | Firmicutes | Clostridia | Clostridiales | Ruminococcaceae | Angelakisella |
| 12 | FALSE | Bacteria | Firmicutes | Clostridia | Clostridiales | Ruminococcaceae | Caproiciproducens |

| | | | | | | | |
|---|---|---|---|---|---|---|---|
| 12 | FALSE | Bacteria | Firmicutes | Clostridia | Clostridiales | Christensenellaceae | Catabacter |
| 12 | FALSE | Bacteria | Firmicutes | Erysipelotrichia | Erysipelotrichales | Erysipelotrichaceae | Coprobacillus |
| 12 | FALSE | Bacteria | Firmicutes | Clostridia | Clostridiales | Lachnospiraceae | GCA-900066755 |
| 12 | FALSE | Bacteria | Firmicutes | Clostridia | Clostridiales | Lachnospiraceae | Lachnospiraceae_NK4B4_group |
| 12 | FALSE | Bacteria | Firmicutes | Negativicutes | Selenomonadales | Veillonellaceae | NA |
| 12 | FALSE | Bacteria | Firmicutes | Clostridia | Clostridiales | Family_XIII | NA |
| 12 | FALSE | Bacteria | Tenericutes | Mollicutes | Izimaplasmatales | NA | NA |
| 12 | FALSE | Bacteria | Proteobacteria | Gammaproteobacteria | Betaproteobacteriales | Burkholderiaceae | Oxalobacter |
| 12 | FALSE | Bacteria | Firmicutes | Clostridia | Clostridiales | Ruminococcaceae | Ruminococcaceae_UCG-004 |
| 12 | FALSE | Bacteria | Firmicutes | Bacilli | Lactobacillales | Leuconostocaceae | Weissella |
| 11 | FALSE | Bacteria | Actinobacteria | Coriobacteriia | Coriobacteriales | Eggerthellaceae | Adlercreutzia |
| 11 | FALSE | Bacteria | Firmicutes | Clostridia | Clostridiales | Christensenellaceae | Christensenellaceae_R-7_group |
| 11 | FALSE | Bacteria | Actinobacteria | Coriobacteriia | Coriobacteriales | Eggerthellaceae | Gordonibacter |
| 11 | FALSE | Bacteria | Firmicutes | Clostridia | Clostridiales | Ruminococcaceae | Harryflintia |
| 11 | FALSE | Bacteria | Firmicutes | Clostridia | Clostridiales | Lachnospiraceae | Lachnospiraceae_FCS020_group |
| 11 | FALSE | Bacteria | Firmicutes | Clostridia | Clostridiales | Lachnospiraceae | Lachnospiraceae_UCG-010 |
| 11 | FALSE | Bacteria | Firmicutes | NA | NA | NA | NA |
| 11 | FALSE | Bacteria | Bacteroidetes | Bacteroidia | NA | NA | NA |
| 11 | FALSE | Bacteria | Actinobacteria | Actinobacteria | Micrococcales | Micrococcaceae | Rothia |
| 11 | FALSE | Bacteria | Firmicutes | Clostridia | Clostridiales | Lachnospiraceae | Shuttleworthia |
| 10 | FALSE | Bacteria | Firmicutes | Bacilli | Lactobacillales | Streptococcaceae | Lactococcus |
| 10 | FALSE | Bacteria | Firmicutes | Bacilli | Bacillales | Bacillaceae | NA |
| 10 | FALSE | Bacteria | Firmicutes | Bacilli | Bacillales | Paenibacillaceae | Paenibacillus |
| 10 | FALSE | Bacteria | Actinobacteria | Coriobacteriia | Coriobacteriales | Eggerthellaceae | Slackia |
| 9 | FALSE | Bacteria | Firmicutes | Erysipelotrichia | Erysipelotrichales | Erysipelotrichaceae | Candidatus_Stoquefichus |
| 9 | FALSE | Bacteria | Firmicutes | Clostridia | Clostridiales | Family_XIII | Family_XIII_AD3011_group |
| 9 | FALSE | Bacteria | Firmicutes | Negativicutes | Selenomonadales | Veillonellaceae | Megamonas |
| 9 | FALSE | Bacteria | Patescibacteria | Saccharimonadia | Saccharimonadales | Saccharimonadaceae | NA |
| 9 | FALSE | Bacteria | Actinobacteria | Coriobacteriia | Coriobacteriales | Coriobacteriales_Incertae_Sedis | NA |
| 9 | FALSE | Bacteria | Proteobacteria | Deltaproteobacteria | Desulfovibrionales | Desulfovibrionaceae | NA |
| 9 | FALSE | Bacteria | Firmicutes | Bacilli | Bacillales | Bacillaceae | Oceanobacillus |
| 9 | FALSE | Bacteria | Firmicutes | Clostridia | Clostridiales | Ruminococcaceae | Oscillibacter |
| 9 | FALSE | Bacteria | Firmicutes | Clostridia | Clostridiales | Peptostreptococcaceae | Romboutsia |
| 9 | FALSE | Bacteria | Proteobacteria | Gammaproteobacteria | Betaproteobacteriales | Burkholderiaceae | Sutterella |
| 8 | FALSE | Bacteria | Actinobacteria | Coriobacteriia | Coriobacteriales | Eggerthellaceae | Eggerthella |

| | | | | | | | |
|---|---|---|---|---|---|---|---|
| 8 | FALSE | Bacteria | Firmicutes | Clostridia | Clostridiales | Ruminococcaceae | Flavonifractor |
| 8 | FALSE | Bacteria | Proteobacteria | Gammaproteobacteria | Enterobacteriales | Enterobacteriaceae | Morganella |
| 8 | FALSE | Bacteria | Firmicutes | Bacilli | Bacillales | Planococcaceae | NA |
| 8 | FALSE | Bacteria | Bacteroidetes | Bacteroidia | Bacteroidales | NA | NA |
| 8 | FALSE | Bacteria | Cyanobacteria | Melainabacteria | Gastranaerophilales | NA | NA |
| 8 | FALSE | Bacteria | Firmicutes | Clostridia | Clostridiales | Lachnospiraceae | Tyzzerella_3 |
| 7 | FALSE | Bacteria | Firmicutes | Clostridia | Clostridiales | Lachnospiraceae | Coprococcus_1 |
| 7 | FALSE | Bacteria | Firmicutes | Clostridia | Clostridiales | Lachnospiraceae | Marvinbryantia |
| 7 | FALSE | Bacteria | Bacteroidetes | Bacteroidia | Bacteroidales | Prevotellaceae | Prevotella_7 |
| 7 | FALSE | Bacteria | Firmicutes | Clostridia | Clostridiales | Ruminococcaceae | Ruminococcaceae_UCG-010 |
| 7 | FALSE | Bacteria | Firmicutes | Bacilli | Bacillales | Staphylococcaceae | Staphylococcus |
| 7 | FALSE | Bacteria | Firmicutes | Clostridia | Clostridiales | Lachnospiraceae | UC5-1-2E3 |
| 6 | FALSE | Bacteria | Proteobacteria | Deltaproteobacteria | Desulfovibrionales | Desulfovibrionaceae | Bilophila |
| 6 | FALSE | Bacteria | Firmicutes | Clostridia | Clostridiales | Lachnospiraceae | CAG-56 |
| 6 | FALSE | Bacteria | Firmicutes | Clostridia | Clostridiales | Lachnospiraceae | Coprococcus_2 |
| 6 | FALSE | Bacteria | Firmicutes | Clostridia | Clostridiales | Clostridiales_vadinBB60_group | NA |
| 6 | FALSE | Bacteria | Bacteroidetes | Bacteroidia | Bacteroidales | Tannerellaceae | Parabacteroides |
| 6 | FALSE | Bacteria | Bacteroidetes | Bacteroidia | Bacteroidales | Prevotellaceae | Prevotella_9 |
| 5 | FALSE | Bacteria | Bacteroidetes | Bacteroidia | Bacteroidales | Rikenellaceae | Alistipes |
| 5 | FALSE | Bacteria | Firmicutes | Negativicutes | Selenomonadales | Veillonellaceae | Dialister |
| 5 | FALSE | Bacteria | Firmicutes | Clostridia | Clostridiales | Lachnospiraceae | Dorea |
| 4 | FALSE | Bacteria | Firmicutes | Clostridia | Clostridiales | Lachnospiraceae | Lachnospiraceae_NK4A136_group |
| 4 | FALSE | Bacteria | Firmicutes | Negativicutes | Selenomonadales | Acidaminococcaceae | Phascolarctobacterium |
| 3 | FALSE | Bacteria | Proteobacteria | Gammaproteobacteria | Enterobacteriales | Enterobacteriaceae | Escherichia/Shigella |
| 3 | FALSE | Bacteria | Proteobacteria | Alphaproteobacteria | Rhodospirillales | NA | NA |
| 3 | FALSE | Bacteria | Proteobacteria | Gammaproteobacteria | Betaproteobacteriales | Burkholderiaceae | Parasutterella |
| 2 | FALSE | Bacteria | Actinobacteria | Coriobacteriia | Coriobacteriales | Coriobacteriaceae | Collinsella |
| 2 | FALSE | Bacteria | Firmicutes | Clostridia | Clostridiales | Ruminococcaceae | Fournierella |
| 2 | FALSE | Bacteria | Proteobacteria | Gammaproteobacteria | Enterobacteriales | Enterobacteriaceae | Klebsiella |
| 2 | FALSE | Bacteria | Bacteroidetes | Bacteroidia | Bacteroidales | Marinifilaceae | Odoribacter |
| 2 | FALSE | Bacteria | Bacteroidetes | Bacteroidia | Bacteroidales | Prevotellaceae | Paraprevotella |
| 1 | FALSE | Bacteria | Firmicutes | Clostridia | Clostridiales | Clostridiaceae_1 | Clostridium_sensu_stricto_1 |
| 1 | FALSE | Bacteria | Firmicutes | Clostridia | Clostridiales | Lachnospiraceae | Coprococcus_3 |
| 1 | FALSE | Bacteria | Firmicutes | Bacilli | Lactobacillales | Enterococcaceae | Enterococcus |
| 1 | FALSE | Bacteria | Firmicutes | Clostridia | Clostridiales | Peptostreptococcaceae | Intestinibacter |

| | | | | | | | |
|---|---|---|---|---|---|---|---|
| 1 | FALSE | Bacteria | Firmicutes | Clostridia | Clostridiales | Ruminococcaceae | Negativibacillus |
| 1 | FALSE | Bacteria | Proteobacteria | Gammaproteobacteria | Enterobacteriales | Enterobacteriaceae | Proteus |
| 1 | FALSE | Bacteria | Firmicutes | Clostridia | Clostridiales | Ruminococcaceae | Ruminococcaceae_NK4A214_group |
| 1 | FALSE | Bacteria | Firmicutes | Clostridia | Clostridiales | Ruminococcaceae | Ruminococcaceae_UCG-003 |
| 0 | FALSE | Bacteria | Verrucomicrobia | Verrucomicrobiae | Verrucomicrobiales | Akkermansiaceae | Akkermansia |
| 0 | FALSE | Bacteria | Bacteroidetes | Bacteroidia | Bacteroidales | Barnesiellaceae | Barnesiella |
| 0 | FALSE | Bacteria | Proteobacteria | Gammaproteobacteria | Enterobacteriales | Enterobacteriaceae | NA |
| 0 | FALSE | Bacteria | Firmicutes | Bacilli | Bacillales | NA | NA |
| 0 | FALSE | Bacteria | Firmicutes | Clostridia | Clostridiales | Ruminococcaceae | Ruminococcaceae_UCG-002 |
| 0 | FALSE | Bacteria | Firmicutes | Clostridia | Clostridiales | Ruminococcaceae | Subdoligranulum |
| 0 | FALSE | Bacteria | Firmicutes | Clostridia | Clostridiales | Lachnospiraceae | Tyzzerella |

**Supplementary Table 4. MWAS of dataset 1 conducted using Kruskal-Wallis**

MRA= mean relative abundance, FC=fold change in patients (PD MRA/control MRA), P= unadjusted significance, FDR (BH)= false discovery rate, adjusted significance. Unclassified genera and genera present in <10% of subjects were excluded from this analysis.

| PD MRA | Control MRA | FC | P | FDR (BH) | Kingdom | Phylum | Class | Order | Family | Genus |
|---|---|---|---|---|---|---|---|---|---|---|
| 0.0005 | 0.0012 | 0.37 | 4E-06 | 2E-04 | Bacteria | Firmicutes | Clostridia | Clostridiales | Lachnospiraceae | Lachnospiraceae_ND3007_group |
| 0.0027 | 0.0004 | 6.61 | 2E-06 | 2E-04 | Bacteria | Firmicutes | Bacilli | Lactobacillales | Lactobacillaceae | Lactobacillus |
| 0.0191 | 0.0362 | 0.53 | 7E-06 | 2E-04 | Bacteria | Firmicutes | Clostridia | Clostridiales | Lachnospiraceae | Agathobacter |
| 0.0153 | 0.0084 | 1.83 | 5E-05 | 1E-03 | Bacteria | Actinobacteria | Actinobacteria | Bifidobacteriales | Bifidobacteriaceae | Bifidobacterium |
| 0.0008 | 0.0001 | 13.03 | 8E-05 | 1E-03 | Bacteria | Synergistetes | Synergistia | Synergistales | Synergistaceae | Cloacibacillus |
| 0.0353 | 0.0564 | 0.63 | 9E-05 | 1E-03 | Bacteria | Firmicutes | Clostridia | Clostridiales | Ruminococcaceae | Faecalibacterium |
| 0.0036 | 0.0004 | 8.14 | 9E-05 | 1E-03 | Bacteria | Firmicutes | Clostridia | Clostridiales | Lachnospiraceae | Hungatella |
| 0.0029 | 0.0037 | 0.80 | 1E-04 | 1E-03 | Bacteria | Firmicutes | Clostridia | Clostridiales | Lachnospiraceae | Lachnospira |
| 0.0047 | 0.0012 | 3.77 | 1E-04 | 1E-03 | Bacteria | Firmicutes | Negativicutes | Selenomonadales | Veillonellaceae | Megasphaera |
| 0.0034 | 0.0008 | 4.20 | 1E-04 | 1E-03 | Bacteria | Bacteroidetes | Bacteroidia | Bacteroidales | Porphyromonadaceae | Porphyromonas |
| 0.0140 | 0.0205 | 0.68 | 2E-04 | 2E-03 | Bacteria | Firmicutes | Clostridia | Clostridiales | Lachnospiraceae | Blautia |
| 0.0017 | 0.0001 | 12.72 | 4E-04 | 4E-03 | Bacteria | Firmicutes | Erysipelotrichia | Erysipelotrichales | Erysipelotrichaceae | Coprobacillus |
| 0.0077 | 0.0160 | 0.48 | 4E-04 | 4E-03 | Bacteria | Firmicutes | Clostridia | Clostridiales | Lachnospiraceae | Roseburia |
| 0.0038 | 0.0015 | 2.56 | 7E-04 | 6E-03 | Bacteria | Bacteroidetes | Bacteroidia | Bacteroidales | Prevotellaceae | Prevotella |
| 0.0541 | 0.0218 | 2.48 | 1E-03 | 7E-03 | Bacteria | Verrucomicrobia | Verrucomicrobiae | Verrucomicrobiales | Akkermansiaceae | Akkermansia |
| 0.0012 | 0.0019 | 0.66 | 1E-03 | 7E-03 | Bacteria | Firmicutes | Clostridia | Clostridiales | Ruminococcaceae | Butyricicoccus |
| 0.0045 | 0.0023 | 1.95 | 1E-03 | 8E-03 | Bacteria | Firmicutes | Clostridia | Clostridiales | Ruminococcaceae | UBA1819 |
| 0.0005 | 0.0001 | 4.95 | 2E-03 | 0.01 | Bacteria | Actinobacteria | Actinobacteria | Actinomycetales | Actinomycetaceae | Varibaculum |
| 0.0020 | 0.0010 | 1.96 | 2E-03 | 0.01 | Bacteria | Actinobacteria | Actinobacteria | Corynebacteriales | Corynebacteriaceae | Corynebacterium_1 |
| 0.0006 | 0.0003 | 1.76 | 3E-03 | 0.01 | Bacteria | Firmicutes | Clostridia | Clostridiales | Ruminococcaceae | Ruminococcaceae_UCG-004 |
| 0.0021 | 0.0038 | 0.56 | 3E-03 | 0.02 | Bacteria | Firmicutes | Clostridia | Clostridiales | Lachnospiraceae | Fusicatenibacter |
| 0.0003 | 0.0007 | 0.48 | 4E-03 | 0.02 | Bacteria | Firmicutes | Clostridia | Clostridiales | Lachnospiraceae | Lachnospiraceae_UCG-004 |
| 0.0004 | 0.0006 | 0.65 | 4E-03 | 0.02 | Bacteria | Firmicutes | Clostridia | Clostridiales | Ruminococcaceae | Oscillospira |
| 0.0027 | 0.0040 | 0.69 | 5E-03 | 0.02 | Bacteria | Firmicutes | Clostridia | Clostridiales | Lachnospiraceae | Anaerostipes |

| | | | | | | | | | | |
|---|---|---|---|---|---|---|---|---|---|---|
| 0.0015 | 0.0013 | 1.16 | 5E-03 | 0.02 | Bacteria | Proteobacteria | Deltaproteobacteria | Desulfovibrionales | Desulfovibrionaceae | Desulfovibrio |
| 0.0006 | 0.0002 | 2.86 | 6E-03 | 0.03 | Bacteria | Firmicutes | Clostridia | Clostridiales | Ruminococcaceae | Anaerotruncus |
| 0.0006 | 0.0004 | 1.37 | 7E-03 | 0.03 | Archaea | Euryarchaeota | Methanobacteria | Methanobacteriales | Methanobacteriaceae | Methanobrevibacter |
| 0.0071 | 0.0021 | 3.32 | 8E-03 | 0.03 | Bacteria | Firmicutes | Clostridia | Clostridiales | Family_XI | Ezakiella |
| 0.0003 | 0.0017 | 0.17 | 9E-03 | 0.03 | Bacteria | Proteobacteria | Gammaproteobacteria | Pasteurellales | Pasteurellaceae | Haemophilus |
| 0.2148 | 0.2479 | 0.87 | 0.01 | 0.04 | Bacteria | Bacteroidetes | Bacteroidia | Bacteroidales | Bacteroidaceae | Bacteroides |
| 0.0043 | 0.0029 | 1.47 | 0.01 | 0.05 | Bacteria | Proteobacteria | Deltaproteobacteria | Desulfovibrionales | Desulfovibrionaceae | Bilophila |
| 0.0014 | 0.0018 | 0.77 | 0.02 | 0.07 | Bacteria | Firmicutes | Clostridia | Clostridiales | Clostridiaceae_1 | Clostridium_sensu_stricto_1 |
| 0.0008 | 0.0013 | 0.64 | 0.02 | 0.07 | Bacteria | Firmicutes | Clostridia | Clostridiales | Lachnospiraceae | Coprococcus_3 |
| 0.0006 | 0.0005 | 1.13 | 0.03 | 0.09 | Bacteria | Firmicutes | Clostridia | Clostridiales | Family_XIII | Family_XIII_AD3011_group |
| 0.0012 | 0.0009 | 1.32 | 0.03 | 0.10 | Bacteria | Firmicutes | Clostridia | Clostridiales | Ruminococcaceae | Ruminococcaceae_UCG-010 |
| 0.0013 | 0.0014 | 0.95 | 0.03 | 0.10 | Bacteria | Firmicutes | Clostridia | Clostridiales | Ruminococcaceae | Ruminococcaceae_UCG-013 |
| 0.0021 | 0.0014 | 1.42 | 0.04 | 0.10 | Bacteria | Firmicutes | Clostridia | Clostridiales | Lachnospiraceae | Eisenbergiella |
| 0.0023 | 0.0009 | 2.58 | 0.04 | 0.10 | Bacteria | Firmicutes | Clostridia | Clostridiales | Family_XI | Peptoniphilus |
| 0.0008 | 0.0002 | 3.81 | 0.04 | 0.10 | Bacteria | Epsilonbacteraeota | Campylobacteria | Campylobacterales | Campylobacteraceae | Campylobacter |
| 0.0003 | 0.0001 | 2.12 | 0.04 | 0.11 | Bacteria | Firmicutes | Clostridia | Clostridiales | Family_XI | Murdochiella |
| 0.0052 | 0.0079 | 0.66 | 0.04 | 0.12 | Bacteria | Firmicutes | Negativicutes | Selenomonadales | Acidaminococcaceae | Phascolarctobacterium |
| 0.0060 | 0.0092 | 0.65 | 0.04 | 0.12 | Bacteria | Firmicutes | Clostridia | Clostridiales | Lachnospiraceae | Lachnoclostridium |
| 0.0020 | 0.0005 | 4.09 | 0.05 | 0.13 | Bacteria | Bacteroidetes | Bacteroidia | Bacteroidales | Prevotellaceae | Prevotella_6 |
| 0.0009 | 0.0011 | 0.86 | 0.06 | 0.15 | Bacteria | Firmicutes | Clostridia | Clostridiales | Ruminococcaceae | Intestinimonas |
| 0.0302 | 0.0233 | 1.29 | 0.07 | 0.15 | Bacteria | Bacteroidetes | Bacteroidia | Bacteroidales | Tannerellaceae | Parabacteroides |
| 0.0070 | 0.0067 | 1.05 | 0.06 | 0.15 | Bacteria | Firmicutes | Clostridia | Clostridiales | Ruminococcaceae | Ruminococcaceae_UCG-005 |
| 0.0006 | 0.0001 | 5.47 | 0.07 | 0.16 | Bacteria | Firmicutes | Clostridia | Clostridiales | Lachnospiraceae | Sellimonas |
| 0.0015 | 0.0004 | 3.31 | 0.08 | 0.17 | Bacteria | Firmicutes | Clostridia | Clostridiales | Family_XI | Anaerococcus |
| 0.0020 | 0.0031 | 0.67 | 0.08 | 0.17 | Bacteria | Firmicutes | Erysipelotrichia | Erysipelotrichales | Erysipelotrichaceae | Erysipelotrichaceae_UCG-003 |
| 0.0036 | 0.0032 | 1.14 | 0.08 | 0.17 | Bacteria | Firmicutes | Clostridia | Clostridiales | Ruminococcaceae | Oscillibacter |
| 0.0015 | 0.0022 | 0.69 | 0.09 | 0.20 | Bacteria | Firmicutes | Clostridia | Clostridiales | Ruminococcaceae | Ruminococcaceae_UCG-003 |
| 0.0002 | 0.0002 | 0.80 | 0.11 | 0.23 | Bacteria | Actinobacteria | Coriobacteriia | Coriobacteriales | Eggerthellaceae | Eggerthella |
| 0.0002 | 0.0001 | 3.08 | 0.11 | 0.23 | Bacteria | Firmicutes | Clostridia | Clostridiales | Ruminococcaceae | Hydrogenoanaerobacterium |
| 0.0020 | 0.0016 | 1.28 | 0.12 | 0.24 | Bacteria | Bacteroidetes | Bacteroidia | Bacteroidales | Marinifilaceae | Butyricimonas |

| | | | | | | | | | | |
|---|---|---|---|---|---|---|---|---|---|---|
| 0.0032 | 0.0014 | 2.30 | 0.13 | 0.25 | Bacteria | Fusobacteria | Fusobacteriia | Fusobacteriales | Fusobacteriaceae | Fusobacterium |
| 0.0027 | 0.0014 | 1.96 | 0.13 | 0.25 | Bacteria | Firmicutes | Erysipelotrichia | Erysipelotrichales | Erysipelotrichaceae | Holdemanella |
| 0.0038 | 0.0024 | 1.60 | 0.14 | 0.27 | Bacteria | Firmicutes | Clostridia | Clostridiales | Ruminococcaceae | Ruminiclostridium_5 |
| 0.0039 | 0.0022 | 1.74 | 0.15 | 0.27 | Bacteria | Firmicutes | Negativicutes | Selenomonadales | Acidaminococcaceae | Acidaminococcus |
| 0.0058 | 0.0034 | 1.69 | 0.15 | 0.27 | Bacteria | Firmicutes | Clostridia | Clostridiales | Lachnospiraceae | Tyzzerella_4 |
| 0.0386 | 0.0337 | 1.15 | 0.16 | 0.28 | Bacteria | Bacteroidetes | Bacteroidia | Bacteroidales | Rikenellaceae | Alistipes |
| 0.0015 | 0.0004 | 3.79 | 0.16 | 0.28 | Bacteria | Firmicutes | Erysipelotrichia | Erysipelotrichales | Erysipelotrichaceae | Faecalitalea |
| 0.0003 | 0.0001 | 4.63 | 0.16 | 0.28 | Bacteria | Lentisphaerae | Lentisphaeria | Victivallales | Victivallaceae | Victivallis |
| 0.0003 | 0.0005 | 0.57 | 0.17 | 0.29 | Bacteria | Firmicutes | Clostridia | Clostridiales | Lachnospiraceae | CAG-56 |
| 0.0006 | 0.0009 | 0.64 | 0.18 | 0.30 | Bacteria | Firmicutes | Clostridia | Clostridiales | Lachnospiraceae | Lachnospiraceae_UCG-010 |
| 0.0161 | 0.0109 | 1.48 | 0.19 | 0.32 | Bacteria | Firmicutes | Clostridia | Clostridiales | Ruminococcaceae | Ruminococcaceae_UCG-002 |
| 0.0010 | 0.0007 | 1.45 | 0.24 | 0.39 | Bacteria | Firmicutes | Clostridia | Clostridiales | Family_XI | Finegoldia |
| 0.0002 | 0.0003 | 0.65 | 0.25 | 0.40 | Bacteria | Firmicutes | Clostridia | Clostridiales | Ruminococcaceae | GCA-900066225 |
| 0.0065 | 0.0046 | 1.41 | 0.26 | 0.42 | Bacteria | Firmicutes | Clostridia | Clostridiales | Christensenellaceae | Christensenellaceae_R-7_group |
| 0.0004 | 0.0001 | 3.97 | 0.27 | 0.42 | Bacteria | Firmicutes | Erysipelotrichia | Erysipelotrichales | Erysipelotrichaceae | Dielma |
| 0.0072 | 0.0071 | 1.02 | 0.28 | 0.44 | Bacteria | Firmicutes | Clostridia | Clostridiales | Ruminococcaceae | Ruminococcus_1 |
| 0.0024 | 0.0029 | 0.83 | 0.29 | 0.45 | Bacteria | Firmicutes | Clostridia | Clostridiales | Lachnospiraceae | Dorea |
| 0.0002 | 0.0003 | 0.75 | 0.31 | 0.48 | Bacteria | Firmicutes | Clostridia | Clostridiales | Ruminococcaceae | DTU089 |
| 0.0015 | 0.0013 | 1.15 | 0.32 | 0.48 | Bacteria | Firmicutes | Clostridia | Clostridiales | Peptostreptococcaceae | Intestinibacter |
| 0.0038 | 0.0032 | 1.19 | 0.32 | 0.48 | Bacteria | Firmicutes | Clostridia | Clostridiales | Ruminococcaceae | Ruminococcaceae_NK4A214_group |
| 0.0033 | 0.0040 | 0.82 | 0.34 | 0.50 | Bacteria | Proteobacteria | Gammaproteobacteria | Betaproteobacteriales | Burkholderiaceae | Sutterella |
| 0.0077 | 0.0082 | 0.94 | 0.36 | 0.52 | Bacteria | Firmicutes | Clostridia | Clostridiales | Ruminococcaceae | Ruminococcaceae_UCG-014 |
| 0.0023 | 0.0023 | 0.99 | 0.39 | 0.55 | Bacteria | Bacteroidetes | Bacteroidia | Bacteroidales | Marinifilaceae | Odoribacter |
| 0.0004 | 0.0004 | 0.94 | 0.43 | 0.59 | Bacteria | Firmicutes | Erysipelotrichia | Erysipelotrichales | Erysipelotrichaceae | Holdemania |
| 0.0015 | 0.0012 | 1.26 | 0.43 | 0.60 | Bacteria | Firmicutes | Clostridia | Clostridiales | Peptostreptococcaceae | Romboutsia |
| 0.0006 | 0.0005 | 1.19 | 0.44 | 0.60 | Bacteria | Firmicutes | Clostridia | Clostridiales | Lachnospiraceae | Coprococcus_1 |
| 0.0087 | 0.0126 | 0.69 | 0.45 | 0.60 | Bacteria | Proteobacteria | Gammaproteobacteria | Pseudomonadales | Moraxellaceae | Acinetobacter |
| 0.0005 | 0.0008 | 0.66 | 0.45 | 0.60 | Bacteria | Firmicutes | Erysipelotrichia | Erysipelotrichales | Erysipelotrichaceae | Erysipelatoclostridium |
| 0.0003 | 0.0005 | 0.51 | 0.47 | 0.61 | Bacteria | Bacteroidetes | Bacteroidia | Bacteroidales | Barnesiellaceae | Coprobacter |
| 0.0020 | 0.0018 | 1.10 | 0.47 | 0.61 | Bacteria | Firmicutes | Clostridia | Clostridiales | Ruminococcaceae | Negativibacillus |

| | | | | | | | | | | |
|---|---|---|---|---|---|---|---|---|---|---|
| 0.0010 | 0.0029 | 0.35 | 0.46 | 0.61 | Bacteria | Bacteroidetes | Bacteroidia | Bacteroidales | Prevotellaceae | Prevotella_7 |
| 0.0042 | 0.0054 | 0.79 | 0.49 | 0.62 | Bacteria | Firmicutes | Clostridia | Clostridiales | Lachnospiraceae | Lachnospiraceae_NK4A136_group |
| 0.0040 | 0.0023 | 1.72 | 0.50 | 0.63 | Bacteria | Firmicutes | Bacilli | Lactobacillales | Streptococcaceae | Streptococcus |
| 0.0064 | 0.0058 | 1.12 | 0.52 | 0.64 | Bacteria | Bacteroidetes | Bacteroidia | Bacteroidales | Barnesiellaceae | Barnesiella |
| 0.0121 | 0.0133 | 0.90 | 0.52 | 0.64 | Bacteria | Firmicutes | Clostridia | Clostridiales | Ruminococcaceae | Subdoligranulum |
| 0.0007 | 0.0005 | 1.44 | 0.56 | 0.68 | Bacteria | Firmicutes | Erysipelotrichia | Erysipelotrichales | Erysipelotrichaceae | Turicibacter |
| 0.0024 | 0.0026 | 0.90 | 0.58 | 0.69 | Bacteria | Bacteroidetes | Bacteroidia | Bacteroidales | Prevotellaceae | Paraprevotella |
| 0.0010 | 0.0009 | 1.10 | 0.59 | 0.70 | Bacteria | Actinobacteria | Coriobacteriia | Coriobacteriales | Coriobacteriaceae | Collinsella |
| 0.0026 | 0.0024 | 1.08 | 0.60 | 0.70 | Bacteria | Firmicutes | Clostridia | Clostridiales | Ruminococcaceae | Ruminiclostridium_9 |
| 0.0012 | 0.0006 | 1.95 | 0.61 | 0.70 | Bacteria | Firmicutes | Negativicutes | Selenomonadales | Veillonellaceae | Veillonella |
| 0.0008 | 0.0005 | 1.45 | 0.62 | 0.71 | Bacteria | Firmicutes | Bacilli | Lactobacillales | Enterococcaceae | Enterococcus |
| 0.0001 | 0.0002 | 0.82 | 0.63 | 0.72 | Bacteria | Firmicutes | Clostridia | Clostridiales | Ruminococcaceae | Angelakisella |
| 0.1131 | 0.1351 | 0.84 | 0.64 | 0.72 | Bacteria | Proteobacteria | Gammaproteobacteria | Enterobacteriales | Enterobacteriaceae | Escherichia/Shigella |
| 0.0021 | 0.0014 | 1.47 | 0.64 | 0.72 | Bacteria | Firmicutes | Clostridia | Clostridiales | Lachnospiraceae | Tyzzerella |
| 0.0017 | 0.0010 | 1.74 | 0.74 | 0.81 | Bacteria | Firmicutes | Clostridia | Clostridiales | Ruminococcaceae | CAG-352 |
| 0.0211 | 0.0155 | 1.37 | 0.78 | 0.85 | Bacteria | Proteobacteria | Gammaproteobacteria | Pseudomonadales | Pseudomonadaceae | Pseudomonas |
| 0.0079 | 0.0090 | 0.89 | 0.81 | 0.86 | Bacteria | Firmicutes | Negativicutes | Selenomonadales | Veillonellaceae | Dialister |
| 0.0024 | 0.0026 | 0.92 | 0.81 | 0.86 | Bacteria | Firmicutes | Clostridia | Clostridiales | Ruminococcaceae | Flavonifractor |
| 0.0039 | 0.0047 | 0.83 | 0.82 | 0.86 | Bacteria | Proteobacteria | Gammaproteobacteria | Betaproteobacteriales | Burkholderiaceae | Parasutterella |
| 0.0115 | 0.0080 | 1.45 | 0.81 | 0.86 | Bacteria | Firmicutes | Clostridia | Clostridiales | Ruminococcaceae | Ruminococcus_2 |
| 0.0040 | 0.0052 | 0.76 | 0.85 | 0.88 | Bacteria | Firmicutes | Clostridia | Clostridiales | Ruminococcaceae | Ruminiclostridium_6 |
| 0.0006 | 0.0007 | 0.81 | 0.88 | 0.90 | Bacteria | Firmicutes | Clostridia | Clostridiales | Lachnospiraceae | Lachnospiraceae_UCG-001 |
| 0.0106 | 0.0155 | 0.68 | 0.88 | 0.90 | Bacteria | Bacteroidetes | Bacteroidia | Bacteroidales | Prevotellaceae | Prevotella_9 |
| 0.0002 | 0.0002 | 0.94 | 0.91 | 0.92 | Bacteria | Firmicutes | Clostridia | Clostridiales | Lachnospiraceae | GCA-900066575 |
| 0.0020 | 0.0015 | 1.36 | 1.00 | 1.00 | Bacteria | Firmicutes | Clostridia | Clostridiales | Lachnospiraceae | Coprococcus_2 |

**Supplementary Table 5. MWAS of dataset 2 conducted using Kruskal-Wallis**

MRA= mean relative abundance, FC=fold change in patients (PD MRA/control MRA), P= unadjusted significance, FDR (BH)= false discovery rate, adjusted significance. Unclassified genera and genera present in <10% of subjects were excluded from this analysis. MRA values of 0.0000 correspond to MRAs that were <0.0001.

| PD MRA | Control MRA | FC | P | FDR (BH) | Kingdom | Phylum | Class | Order | Family | Genus |
|---|---|---|---|---|---|---|---|---|---|---|
| 0.0239 | 0.0088 | 2.72 | 4E-09 | 6E-07 | Bacteria | Actinobacteria | Actinobacteria | Bifidobacteriales | Bifidobacteriaceae | Bifidobacterium |
| 0.0004 | 0.0011 | 0.38 | 2E-07 | 1E-05 | Bacteria | Firmicutes | Clostridia | Clostridiales | Lachnospiraceae | Lachnospiraceae_UCG-004 |
| 0.0097 | 0.0172 | 0.56 | 1E-06 | 6E-05 | Bacteria | Firmicutes | Clostridia | Clostridiales | Lachnospiraceae | Agathobacter |
| 0.0047 | 0.0078 | 0.60 | 7E-06 | 3E-04 | Bacteria | Firmicutes | Clostridia | Clostridiales | Lachnospiraceae | Roseburia |
| 0.0002 | 0.0001 | 2.19 | 8E-06 | 3E-04 | Bacteria | Firmicutes | Clostridia | Clostridiales | Eubacteriaceae | Eubacterium |
| 0.0007 | 0.0011 | 0.59 | 2E-05 | 6E-04 | Bacteria | Firmicutes | Clostridia | Clostridiales | Lachnospiraceae | Lachnospiraceae_ND3007_group |
| 0.0013 | 0.0018 | 0.73 | 3E-05 | 7E-04 | Bacteria | Firmicutes | Clostridia | Clostridiales | Ruminococcaceae | Ruminococcaceae_UCG-013 |
| 0.0039 | 0.0050 | 0.78 | 7E-05 | 1E-03 | Bacteria | Firmicutes | Clostridia | Clostridiales | Lachnospiraceae | Anaerostipes |
| 0.0003 | 0.0001 | 4.43 | 2E-04 | 3E-03 | Bacteria | Actinobacteria | Actinobacteria | Corynebacteriales | Corynebacteriaceae | Lawsonella |
| 0.0276 | 0.0416 | 0.66 | 2E-04 | 3E-03 | Bacteria | Firmicutes | Clostridia | Clostridiales | Ruminococcaceae | Faecalibacterium |
| 0.0006 | 0.0003 | 2.11 | 4E-04 | 0.01 | Bacteria | Firmicutes | Erysipelotrichia | Erysipelotrichales | Erysipelotrichaceae | Turicibacter |
| 0.0208 | 0.0258 | 0.81 | 6E-04 | 0.01 | Bacteria | Proteobacteria | Gammaproteobacteria | Pseudomonadales | Pseudomonadaceae | Pseudomonas |
| 0.0036 | 0.0026 | 1.38 | 7E-04 | 0.01 | Bacteria | Firmicutes | Clostridia | Clostridiales | Ruminococcaceae | UBA1819 |
| 0.0039 | 0.0015 | 2.53 | 7E-04 | 0.01 | Bacteria | Actinobacteria | Actinobacteria | Corynebacteriales | Corynebacteriaceae | Corynebacterium_1 |
| 0.0001 | 0.0002 | 0.58 | 8E-04 | 0.01 | Bacteria | Firmicutes | Erysipelotrichia | Erysipelotrichales | Erysipelotrichaceae | Erysipelotrichaceae_UCG-003 |
| 0.0017 | 0.0007 | 2.56 | 1E-03 | 0.01 | Bacteria | Firmicutes | Clostridia | Clostridiales | Family_XI | Anaerococcus |
| 0.0003 | 0.0007 | 0.39 | 1E-03 | 0.01 | Bacteria | Firmicutes | Clostridia | Clostridiales | Lachnospiraceae | Lachnospiraceae_UCG-001 |
| 0.0015 | 0.0008 | 1.81 | 1E-03 | 0.01 | Bacteria | Proteobacteria | Deltaproteobacteria | Desulfovibrionales | Desulfovibrionaceae | Desulfovibrio |
| 0.0056 | 0.0036 | 1.57 | 1E-03 | 0.01 | Bacteria | Firmicutes | Bacilli | Lactobacillales | Lactobacillaceae | Lactobacillus |
| 0.0036 | 0.0053 | 0.68 | 2E-03 | 0.01 | Bacteria | Firmicutes | Clostridia | Clostridiales | Lachnospiraceae | Lachnospira |
| 0.0003 | 0.0005 | 0.64 | 2E-03 | 0.01 | Bacteria | Firmicutes | Clostridia | Clostridiales | Ruminococcaceae | Oscillospira |
| 0.0005 | 0.0003 | 1.79 | 2E-03 | 0.01 | Bacteria | Actinobacteria | Actinobacteria | Actinomycetales | Actinomycetaceae | Varibaculum |
| 0.0027 | 0.0012 | 2.14 | 2E-03 | 0.01 | Bacteria | Firmicutes | Clostridia | Clostridiales | Family_XI | Peptoniphilus |
| 0.0015 | 0.0008 | 1.77 | 2E-03 | 0.02 | Bacteria | Firmicutes | Clostridia | Clostridiales | Lachnospiraceae | Hungatella |
| 0.0045 | 0.0028 | 1.61 | 3E-03 | 0.02 | Archaea | Euryarchaeota | Methanobacteria | Methanobacteriales | Methanobacteriaceae | Methanobrevibacter |
| 0.0022 | 0.0000 | 220.2 | 3E-03 | 0.02 | Bacteria | Proteobacteria | Gammaproteobacteria | Betaproteobacteriales | Burkholderiaceae | Delftia |
| 0.0085 | 0.0086 | 0.99 | 3E-03 | 0.02 | Bacteria | Firmicutes | Bacilli | Lactobacillales | Streptococcaceae | Streptococcus |
| 0.0026 | 0.0009 | 2.94 | 3E-03 | 0.02 | Bacteria | Bacteroidetes | Bacteroidia | Bacteroidales | Porphyromonadaceae | Porphyromonas |
| 0.0025 | 0.0006 | 4.39 | 3E-03 | 0.02 | Bacteria | Bacteroidetes | Bacteroidia | Bacteroidales | Prevotellaceae | Prevotella |
| 0.0031 | 0.0046 | 0.69 | 0.01 | 0.03 | Bacteria | Firmicutes | Clostridia | Clostridiales | Lachnospiraceae | Fusicatenibacter |
| 0.0008 | 0.0005 | 1.69 | 0.01 | 0.03 | Bacteria | Firmicutes | Clostridia | Clostridiales | Ruminococcaceae | Anaerotruncus |
| 0.0079 | 0.0101 | 0.78 | 0.01 | 0.03 | Bacteria | Firmicutes | Clostridia | Clostridiales | Ruminococcaceae | Ruminococcus_2 |

| | | | | | | | | | | |
|---|---|---|---|---|---|---|---|---|---|---|
| 0.0001 | 0.0000 | 34.45 | 0.01 | 0.03 | Bacteria | Firmicutes | Clostridia | Clostridiales | Family_XI | Parvimonas |
| 0.0003 | 0.0002 | 1.72 | 0.01 | 0.04 | Bacteria | Actinobacteria | Actinobacteria | Actinomycetales | Actinomycetaceae | Mobiluncus |
| 0.0005 | 0.0003 | 1.45 | 0.01 | 0.04 | Bacteria | Actinobacteria | Actinobacteria | Actinomycetales | Actinomycetaceae | Actinomyces |
| 0.0012 | 0.0005 | 2.60 | 0.01 | 0.04 | Bacteria | Firmicutes | Clostridia | Clostridiales | Family_XI | Finegoldia |
| 0.0003 | 0.0001 | 2.69 | 0.01 | 0.04 | Bacteria | Firmicutes | Clostridia | Clostridiales | Family_XIII | S5-A14a |
| 0.0187 | 0.0237 | 0.79 | 0.01 | 0.04 | Bacteria | Firmicutes | Clostridia | Clostridiales | Lachnospiraceae | Blautia |
| 0.0005 | 0.0002 | 2.25 | 0.01 | 0.04 | Bacteria | Firmicutes | Clostridia | Clostridiales | Family_XI | Murdochiella |
| 0.0048 | 0.0015 | 3.18 | 0.01 | 0.04 | Bacteria | Firmicutes | Clostridia | Clostridiales | Family_XI | Ezakiella |
| 0.0004 | 0.0005 | 0.78 | 0.01 | 0.04 | Bacteria | Firmicutes | Clostridia | Clostridiales | Ruminococcaceae | DTU089 |
| 0.0000 | 0.0000 | 2.06 | 0.01 | 0.05 | Bacteria | Actinobacteria | Actinobacteria | Propionibacteriales | Propionibacteriaceae | Cutibacterium |
| 0.0013 | 0.0019 | 0.68 | 0.02 | 0.06 | Bacteria | Firmicutes | Clostridia | Clostridiales | Ruminococcaceae | Butyricicoccus |
| 0.0000 | 0.0000 | 2.91 | 0.02 | 0.07 | Bacteria | Actinobacteria | Actinobacteria | Bifidobacteriales | Bifidobacteriaceae | Scardovia |
| 0.0034 | 0.0053 | 0.64 | 0.02 | 0.07 | Bacteria | Firmicutes | Clostridia | Clostridiales | Ruminococcaceae | Ruminiclostridium_6 |
| 0.0034 | 0.0044 | 0.79 | 0.02 | 0.07 | Bacteria | Firmicutes | Clostridia | Clostridiales | Ruminococcaceae | Ruminococcus_1 |
| 0.0000 | 0.0000 | 0.72 | 0.02 | 0.07 | Bacteria | Firmicutes | Bacilli | Bacillales | Family_XI | Gemella |
| 0.0004 | 0.0001 | 2.58 | 0.03 | 0.09 | Bacteria | Firmicutes | Clostridia | Clostridiales | Ruminococcaceae | Fastidiosipila |
| 0.0038 | 0.0063 | 0.61 | 0.03 | 0.09 | Bacteria | Firmicutes | Clostridia | Clostridiales | Ruminococcaceae | Ruminococcaceae_UCG-014 |
| 0.0037 | 0.0018 | 2.09 | 0.03 | 0.09 | Bacteria | Firmicutes | Clostridia | Clostridiales | Lachnospiraceae | Eisenbergiella |
| 0.0008 | 0.0007 | 1.23 | 0.03 | 0.09 | Bacteria | Firmicutes | Clostridia | Clostridiales | Ruminococcaceae | Ruminococcaceae_UCG-004 |
| 0.0007 | 0.0002 | 4.09 | 0.03 | 0.09 | Bacteria | Epsilonbacteraeota | Campylobacteria | Campylobacterales | Campylobacteraceae | Campylobacter |
| 0.0027 | 0.0045 | 0.61 | 0.03 | 0.09 | Bacteria | Firmicutes | Negativicutes | Selenomonadales | Veillonellaceae | Veillonella |
| 0.0002 | 0.0000 | 11.43 | 0.03 | 0.10 | Bacteria | Lentisphaerae | Lentisphaeria | Victivallales | Victivallaceae | Victivallis |
| 0.0000 | 0.0001 | 0.59 | 0.04 | 0.12 | Bacteria | Firmicutes | Bacilli | Lactobacillales | Carnobacteriaceae | Granulicatella |
| 0.0000 | 0.0000 | 0.95 | 0.04 | 0.13 | Bacteria | Firmicutes | Clostridia | Clostridiales | Lachnospiraceae | Cuneatibacter |
| 0.0001 | 0.0001 | 1.45 | 0.04 | 0.13 | Bacteria | Firmicutes | Clostridia | Clostridiales | Ruminococcaceae | Ruminococcaceae_UCG-009 |
| 0.0000 | 0.0000 | 1.58 | 0.05 | 0.13 | Bacteria | Firmicutes | Clostridia | Clostridiales | Ruminococcaceae | Ruminococcaceae_UCG-011 |
| 0.0001 | 0.0001 | 0.86 | 0.05 | 0.14 | Bacteria | Firmicutes | Clostridia | Clostridiales | Ruminococcaceae | Candidatus_Soleaferrea |
| 0.0001 | 0.0002 | 0.73 | 0.05 | 0.14 | Bacteria | Proteobacteria | Gammaproteobacteria | Pasteurellales | Pasteurellaceae | Haemophilus |
| 0.0000 | 0.0000 | 0.51 | 0.05 | 0.14 | Bacteria | Firmicutes | Clostridia | Clostridiales | Ruminococcaceae | Ruminococcaceae_UCG-008 |
| 0.0001 | 0.0004 | 0.40 | 0.06 | 0.14 | Bacteria | Firmicutes | Clostridia | Clostridiales | Lachnospiraceae | Tyzzerella_3 |
| 0.0004 | 0.0003 | 1.40 | 0.06 | 0.15 | Bacteria | Firmicutes | Clostridia | Clostridiales | Ruminococcaceae | GCA-900066225 |
| 0.0001 | 0.0002 | 0.58 | 0.06 | 0.16 | Bacteria | Actinobacteria | Actinobacteria | Micrococcales | Brevibacteriaceae | Brevibacterium |
| 0.0002 | 0.0001 | 1.42 | 0.06 | 0.16 | Bacteria | Firmicutes | Clostridia | Clostridiales | Lachnospiraceae | Howardella |
| 0.0023 | 0.0017 | 1.37 | 0.07 | 0.16 | Bacteria | Bacteroidetes | Bacteroidia | Bacteroidales | Marinifilaceae | Butyricimonas |
| 0.0002 | 0.0001 | 2.00 | 0.07 | 0.17 | Bacteria | Firmicutes | Clostridia | Clostridiales | Family_XIII | Mogibacterium |
| 0.0000 | 0.0000 | 1.46 | 0.07 | 0.17 | Bacteria | Firmicutes | Clostridia | Clostridiales | Eubacteriaceae | Anaerofustis |
| 0.0001 | 0.0001 | 0.75 | 0.07 | 0.17 | Bacteria | Firmicutes | Clostridia | Clostridiales | Family_XIII | Family_XIII_UCG-001 |
| 0.0096 | 0.0047 | 2.06 | 0.07 | 0.17 | Bacteria | Firmicutes | Negativicutes | Selenomonadales | Acidaminococcaceae | Acidaminococcus |
| 0.0003 | 0.0002 | 1.33 | 0.07 | 0.17 | Bacteria | Firmicutes | Negativicutes | Selenomonadales | Veillonellaceae | Anaeroglobus |
| 0.0001 | 0.0001 | 0.82 | 0.08 | 0.17 | Bacteria | Firmicutes | Clostridia | Clostridiales | Ruminococcaceae | Phocea |
| 0.0001 | 0.0002 | 0.76 | 0.08 | 0.18 | Bacteria | Firmicutes | Clostridia | Clostridiales | Peptococcaceae | Peptococcus |

| | | | | | | | | | | |
|---|---|---|---|---|---|---|---|---|---|---|
| 0.0373 | 0.0460 | 0.81 | 0.08 | 0.18 | Bacteria | Firmicutes | Negativicutes | Selenomonadales | Acidaminococcaceae | Phascolarctobacterium |
| 0.0000 | 0.0000 | 0.72 | 0.09 | 0.19 | Bacteria | Firmicutes | Clostridia | Clostridiales | Defluviitaleaceae | Defluviitaleaceae_UCG-011 |
| 0.0007 | 0.0004 | 1.68 | 0.10 | 0.20 | Bacteria | Firmicutes | Clostridia | Clostridiales | Lachnospiraceae | Sellimonas |
| 0.0041 | 0.0032 | 1.31 | 0.10 | 0.21 | Bacteria | Firmicutes | Clostridia | Clostridiales | Ruminococcaceae | Ruminococcaceae_UCG-005 |
| 0.0000 | 0.0000 | 0.65 | 0.10 | 0.21 | Bacteria | Firmicutes | Clostridia | Clostridiales | Ruminococcaceae | Ruminococcaceae_UCG-007 |
| 0.0001 | 0.0001 | 0.79 | 0.10 | 0.21 | Bacteria | Firmicutes | Erysipelotrichia | Erysipelotrichales | Erysipelotrichaceae | Erysipelatoclostridium |
| 0.0036 | 0.0008 | 4.50 | 0.10 | 0.21 | Bacteria | Firmicutes | Clostridia | Clostridiales | Lachnospiraceae | Tyzzerella_4 |
| 0.0055 | 0.0040 | 1.36 | 0.10 | 0.21 | Bacteria | Actinobacteria | Coriobacteriia | Coriobacteriales | Coriobacteriaceae | Collinsella |
| 0.1610 | 0.1887 | 0.85 | 0.12 | 0.23 | Bacteria | Proteobacteria | Gammaproteobacteria | Enterobacteriales | Enterobacteriaceae | Escherichia/Shigella |
| 0.0000 | 0.0000 | 0.98 | 0.12 | 0.23 | Bacteria | Firmicutes | Erysipelotrichia | Erysipelotrichales | Erysipelotrichaceae | Holdemania |
| 0.0028 | 0.0028 | 0.99 | 0.14 | 0.26 | Bacteria | Firmicutes | Clostridia | Clostridiales | Lachnospiraceae | Dorea |
| 0.0003 | 0.0003 | 0.90 | 0.15 | 0.28 | Bacteria | Firmicutes | Clostridia | Clostridiales | Lachnospiraceae | CAG-56 |
| 0.0004 | 0.0009 | 0.40 | 0.17 | 0.32 | Bacteria | Firmicutes | Clostridia | Clostridiales | Lachnospiraceae | Coprococcus_2 |
| 0.0051 | 0.0040 | 1.26 | 0.17 | 0.33 | Bacteria | Firmicutes | Clostridia | Clostridiales | Ruminococcaceae | Ruminiclostridium_5 |
| 0.0002 | 0.0003 | 0.86 | 0.18 | 0.33 | Bacteria | Firmicutes | Clostridia | Clostridiales | Lachnospiraceae | GCA-900066575 |
| 0.0032 | 0.0034 | 0.93 | 0.19 | 0.35 | Bacteria | Firmicutes | Clostridia | Clostridiales | Lachnospiraceae | Lachnospiraceae_NK4A136_group |
| 0.0001 | 0.0001 | 1.63 | 0.20 | 0.37 | Bacteria | Firmicutes | Erysipelotrichia | Erysipelotrichales | Erysipelotrichaceae | Faecalitalea |
| 0.0032 | 0.0033 | 0.99 | 0.23 | 0.41 | Bacteria | Firmicutes | Clostridia | Clostridiales | Christensenellaceae | Christensenellaceae_R-7_group |
| 0.0199 | 0.0149 | 1.34 | 0.24 | 0.42 | Bacteria | Verrucomicrobia | Verrucomicrobiae | Verrucomicrobiales | Akkermansiaceae | Akkermansia |
| 0.0018 | 0.0080 | 0.22 | 0.24 | 0.43 | Bacteria | Proteobacteria | Gammaproteobacteria | Enterobacteriales | Enterobacteriaceae | Klebsiella |
| 0.0031 | 0.0032 | 0.98 | 0.26 | 0.46 | Bacteria | Firmicutes | Clostridia | Clostridiales | Ruminococcaceae | Ruminiclostridium_9 |
| 0.0001 | 0.0001 | 1.04 | 0.27 | 0.47 | Bacteria | Firmicutes | Clostridia | Clostridiales | Lachnospiraceae | Shuttleworthia |
| 0.0001 | 0.0000 | 3.41 | 0.28 | 0.48 | Bacteria | Actinobacteria | Coriobacteriia | Coriobacteriales | Atopobiaceae | Atopobium |
| 0.0104 | 0.0114 | 0.91 | 0.30 | 0.50 | Bacteria | Firmicutes | Clostridia | Clostridiales | Lachnospiraceae | Lachnoclostridium |
| 0.0058 | 0.0051 | 1.14 | 0.32 | 0.53 | Bacteria | Firmicutes | Clostridia | Clostridiales | Ruminococcaceae | Oscillibacter |
| 0.0001 | 0.0001 | 1.54 | 0.32 | 0.53 | Bacteria | Firmicutes | Bacilli | Lactobacillales | Streptococcaceae | Lactococcus |
| 0.1960 | 0.2043 | 0.96 | 0.33 | 0.53 | Bacteria | Bacteroidetes | Bacteroidia | Bacteroidales | Bacteroidaceae | Bacteroides |
| 0.0006 | 0.0006 | 0.93 | 0.34 | 0.54 | Bacteria | Actinobacteria | Coriobacteriia | Coriobacteriales | Eggerthellaceae | Eggerthella |
| 0.0044 | 0.0033 | 1.35 | 0.34 | 0.55 | Bacteria | Proteobacteria | Gammaproteobacteria | Betaproteobacteriales | Burkholderiaceae | Parasutterella |
| 0.0020 | 0.0021 | 0.93 | 0.35 | 0.55 | Bacteria | Firmicutes | Clostridia | Clostridiales | Ruminococcaceae | Ruminococcaceae_UCG-003 |
| 0.0000 | 0.0001 | 0.37 | 0.35 | 0.55 | Bacteria | Firmicutes | Clostridia | Clostridiales | Ruminococcaceae | Ruminiclostridium |
| 0.0008 | 0.0007 | 1.16 | 0.35 | 0.55 | Bacteria | Firmicutes | Clostridia | Clostridiales | Lachnospiraceae | Lachnospiraceae_UCG-010 |
| 0.0001 | 0.0001 | 1.53 | 0.36 | 0.55 | Bacteria | Firmicutes | Negativicutes | Selenomonadales | Veillonellaceae | Negativicoccus |
| 0.0042 | 0.0015 | 2.80 | 0.36 | 0.55 | Bacteria | Synergistetes | Synergistia | Synergistales | Synergistaceae | Cloacibacillus |
| 0.0018 | 0.0017 | 1.08 | 0.37 | 0.55 | Bacteria | Proteobacteria | Deltaproteobacteria | Desulfovibrionales | Desulfovibrionaceae | Bilophila |
| 0.0026 | 0.0027 | 0.98 | 0.37 | 0.55 | Bacteria | Firmicutes | Clostridia | Clostridiales | Ruminococcaceae | Ruminococcaceae_NK4A214_group |
| 0.0001 | 0.0001 | 0.67 | 0.37 | 0.55 | Bacteria | Firmicutes | Clostridia | Clostridiales | Lachnospiraceae | Lachnospiraceae_FCS020_group |
| 0.0053 | 0.0037 | 1.45 | 0.38 | 0.56 | Bacteria | Firmicutes | Negativicutes | Selenomonadales | Veillonellaceae | Dialister |
| 0.0006 | 0.0003 | 1.97 | 0.40 | 0.58 | Bacteria | Fusobacteria | Fusobacteriia | Fusobacteriales | Fusobacteriaceae | Fusobacterium |
| 0.0002 | 0.0001 | 2.54 | 0.41 | 0.59 | Bacteria | Firmicutes | Clostridia | Clostridiales | Peptostreptococcaceae | Terrisporobacter |
| 0.0015 | 0.0009 | 1.67 | 0.41 | 0.59 | Bacteria | Proteobacteria | Gammaproteobacteria | Betaproteobacteriales | Burkholderiaceae | Sutterella |

| | | | | | | | | | | |
|---|---|---|---|---|---|---|---|---|---|---|
| 0.0222 | 0.0199 | 1.11 | 0.41 | 0.59 | Bacteria | Bacteroidetes | Bacteroidia | Bacteroidales | Tannerellaceae | Parabacteroides |
| 0.0020 | 0.0024 | 0.83 | 0.42 | 0.59 | Bacteria | Firmicutes | Clostridia | Clostridiales | Ruminococcaceae | Intestinimonas |
| 0.0006 | 0.0002 | 2.78 | 0.43 | 0.60 | Bacteria | Bacteroidetes | Bacteroidia | Bacteroidales | Prevotellaceae | Prevotella_6 |
| 0.0018 | 0.0020 | 0.92 | 0.43 | 0.60 | Bacteria | Firmicutes | Clostridia | Clostridiales | Ruminococcaceae | Negativibacillus |
| 0.0086 | 0.0035 | 2.48 | 0.43 | 0.60 | Bacteria | Firmicutes | Negativicutes | Selenomonadales | Veillonellaceae | Megasphaera |
| 0.0001 | 0.0001 | 0.41 | 0.45 | 0.61 | Bacteria | Firmicutes | Clostridia | Clostridiales | Lachnospiraceae | Lachnospiraceae_NK4B4_group |
| 0.0000 | 0.0001 | 0.76 | 0.45 | 0.61 | Bacteria | Firmicutes | Erysipelotrichia | Erysipelotrichales | Erysipelotrichaceae | Coprobacillus |
| 0.0138 | 0.0112 | 1.24 | 0.46 | 0.62 | Bacteria | Firmicutes | Clostridia | Clostridiales | Ruminococcaceae | Ruminococcaceae_UCG-002 |
| 0.0076 | 0.0131 | 0.58 | 0.49 | 0.64 | Bacteria | Proteobacteria | Gammaproteobacteria | Enterobacteriales | Enterobacteriaceae | Proteus |
| 0.0027 | 0.0020 | 1.39 | 0.49 | 0.64 | Bacteria | Firmicutes | Clostridia | Clostridiales | Peptostreptococcaceae | Romboutsia |
| 0.0001 | 0.0000 | 1.32 | 0.49 | 0.64 | Bacteria | Firmicutes | Clostridia | Clostridiales | Ruminococcaceae | Anaerofilum |
| 0.0008 | 0.0004 | 2.10 | 0.50 | 0.64 | Bacteria | Bacteroidetes | Bacteroidia | Bacteroidales | Barnesiellaceae | Coprobacter |
| 0.0006 | 0.0005 | 1.29 | 0.52 | 0.67 | Bacteria | Firmicutes | Bacilli | Bacillales | Staphylococcaceae | Staphylococcus |
| 0.0000 | 0.0000 | 1.08 | 0.52 | 0.67 | Bacteria | Firmicutes | Clostridia | Clostridiales | Christensenellaceae | Catabacter |
| 0.0000 | 0.0000 | 1.19 | 0.53 | 0.67 | Bacteria | Firmicutes | Clostridia | Clostridiales | Lachnospiraceae | GCA-900066755 |
| 0.0004 | 0.0003 | 1.56 | 0.54 | 0.68 | Bacteria | Synergistetes | Synergistia | Synergistales | Synergistaceae | Pyramidobacter |
| 0.0001 | 0.0000 | 2.06 | 0.55 | 0.68 | Bacteria | Firmicutes | Clostridia | Clostridiales | Ruminococcaceae | Ruminiclostridium_1 |
| 0.0001 | 0.0000 | 1.58 | 0.55 | 0.68 | Bacteria | Firmicutes | Erysipelotrichia | Erysipelotrichales | Erysipelotrichaceae | Candidatus_Stoquefichus |
| 0.0004 | 0.0003 | 1.20 | 0.57 | 0.69 | Bacteria | Firmicutes | Clostridia | Clostridiales | Ruminococcaceae | Angelakisella |
| 0.0007 | 0.0006 | 1.26 | 0.57 | 0.70 | Bacteria | Firmicutes | Clostridia | Clostridiales | Family_XIII | Family_XIII_AD3011_group |
| 0.0009 | 0.0010 | 0.89 | 0.58 | 0.70 | Bacteria | Firmicutes | Clostridia | Clostridiales | Lachnospiraceae | Coprococcus_3 |
| 0.0032 | 0.0033 | 0.97 | 0.59 | 0.71 | Bacteria | Firmicutes | Clostridia | Clostridiales | Ruminococcaceae | Flavonifractor |
| 0.0000 | 0.0000 | 1.12 | 0.62 | 0.73 | Bacteria | Firmicutes | Clostridia | Clostridiales | Ruminococcaceae | Acetanaerobacterium |
| 0.0002 | 0.0001 | 1.31 | 0.62 | 0.73 | Bacteria | Actinobacteria | Coriobacteriia | Coriobacteriales | Eggerthellaceae | Slackia |
| 0.0001 | 0.0001 | 0.63 | 0.62 | 0.73 | Bacteria | Actinobacteria | Actinobacteria | Micrococcales | Micrococcaceae | Rothia |
| 0.0000 | 0.0000 | 3.20 | 0.63 | 0.73 | Bacteria | Firmicutes | Erysipelotrichia | Erysipelotrichales | Erysipelotrichaceae | Dielma |
| 0.0041 | 0.0055 | 0.75 | 0.64 | 0.74 | Bacteria | Bacteroidetes | Bacteroidia | Bacteroidales | Prevotellaceae | Prevotella_9 |
| 0.0005 | 0.0002 | 2.14 | 0.65 | 0.75 | Bacteria | Firmicutes | Clostridia | Clostridiales | Lachnospiraceae | Marvinbryantia |
| 0.0004 | 0.0005 | 0.92 | 0.66 | 0.75 | Bacteria | Firmicutes | Clostridia | Clostridiales | Ruminococcaceae | Fournierella |
| 0.0004 | 0.0003 | 1.30 | 0.67 | 0.75 | Bacteria | Firmicutes | Clostridia | Clostridiales | Lachnospiraceae | Coprococcus_1 |
| 0.0007 | 0.0007 | 0.93 | 0.68 | 0.77 | Bacteria | Firmicutes | Clostridia | Clostridiales | Peptostreptococcaceae | Intestinibacter |
| 0.0001 | 0.0001 | 2.34 | 0.70 | 0.78 | Bacteria | Firmicutes | Clostridia | Clostridiales | Ruminococcaceae | Hydrogenoanaerobacterium |
| 0.0052 | 0.0044 | 1.18 | 0.72 | 0.80 | Bacteria | Bacteroidetes | Bacteroidia | Bacteroidales | Barnesiellaceae | Barnesiella |
| 0.0000 | 0.0000 | 1.48 | 0.73 | 0.80 | Bacteria | Firmicutes | Clostridia | Clostridiales | Lachnospiraceae | Lactonifactor |
| 0.0001 | 0.0001 | 1.16 | 0.74 | 0.80 | Bacteria | Actinobacteria | Coriobacteriia | Coriobacteriales | Eggerthellaceae | Gordonibacter |
| 0.0016 | 0.0017 | 0.98 | 0.74 | 0.80 | Bacteria | Firmicutes | Bacilli | Lactobacillales | Enterococcaceae | Enterococcus |
| 0.0003 | 0.0002 | 1.25 | 0.75 | 0.80 | Bacteria | Actinobacteria | Coriobacteriia | Coriobacteriales | Eggerthellaceae | Adlercreutzia |
| 0.0002 | 0.0002 | 0.79 | 0.75 | 0.80 | Bacteria | Firmicutes | Clostridia | Clostridiales | Lachnospiraceae | UC5-1-2E3 |
| 0.0170 | 0.0159 | 1.07 | 0.79 | 0.84 | Bacteria | Firmicutes | Clostridia | Clostridiales | Ruminococcaceae | Subdoligranulum |
| 0.0028 | 0.0015 | 1.82 | 0.80 | 0.84 | Bacteria | Firmicutes | Clostridia | Clostridiales | Clostridiaceae_1 | Clostridium_sensu_stricto_1 |
| 0.0015 | 0.0011 | 1.33 | 0.88 | 0.92 | Bacteria | Bacteroidetes | Bacteroidia | Bacteroidales | Prevotellaceae | Prevotella_7 |

| | | | | | | | | | | |
|---|---|---|---|---|---|---|---|---|---|---|
| 0.0202 | 0.0206 | 0.98 | 0.88 | 0.92 | Bacteria | Bacteroidetes | Bacteroidia | Bacteroidales | Rikenellaceae | Alistipes |
| 0.0006 | 0.0005 | 1.05 | 0.91 | 0.95 | Bacteria | Firmicutes | Clostridia | Clostridiales | Ruminococcaceae | Ruminococcaceae_UCG-010 |
| 0.0000 | 0.0000 | 1.34 | 0.95 | 0.98 | Bacteria | Firmicutes | Clostridia | Clostridiales | Ruminococcaceae | Caproiciproducens |
| 0.0002 | 0.0002 | 1.32 | 0.95 | 0.98 | Bacteria | Firmicutes | Bacilli | Lactobacillales | Leuconostocaceae | Weissella |
| 0.0014 | 0.0009 | 1.51 | 0.96 | 0.98 | Bacteria | Firmicutes | Clostridia | Clostridiales | Lachnospiraceae | Tyzzerella |
| 0.0000 | 0.0000 | 1.23 | 0.98 | 0.99 | Bacteria | Firmicutes | Clostridia | Clostridiales | Ruminococcaceae | Harryflintia |
| 0.0053 | 0.0049 | 1.09 | 0.98 | 0.99 | Bacteria | Bacteroidetes | Bacteroidia | Bacteroidales | Marinifilaceae | Odoribacter |
| 0.0012 | 0.0015 | 0.80 | 0.99 | 0.99 | Bacteria | Bacteroidetes | Bacteroidia | Bacteroidales | Prevotellaceae | Paraprevotella |

**Supplementary Table 6. PubMed search results for *Porphyromonas, Prevotella*, or *Corynebacterium_1***

Species that comprised each genus (and accounted for at least 80% of the ASVs in the genus) were identified based on 100% sequence identity using DADA2-SILVA reference database or 100% or >99% identity and high statistical confidence using NCBI 16S rRNA database. Then each species was searched in PubMed using "genus species" as search term. Search filters: Humans, English, Title/abstract. The citations were tabulated for articles that addressed function, characteristics or relevance to human health; method papers were omitted. All infections are in human samples, except *C. lactis* (newly discovered) was found in an abscess in a companion dog.

| Search term | PubMed Return | Subject matter |
|---|---|---|
| *Corynebacterium amycolatum* | https://www.ncbi.nlm.nih.gov/pubmed/10342655 | septic arthritis |
| *Corynebacterium amycolatum* | https://www.ncbi.nlm.nih.gov/pubmed/10482033 | surgical or catheter related infection, pilonidal cyst |
| *Corynebacterium amycolatum* | https://www.ncbi.nlm.nih.gov/pubmed/11749760 | endocarditis |
| *Corynebacterium amycolatum* | https://www.ncbi.nlm.nih.gov/pubmed/12235925 | blood cultures |
| *Corynebacterium amycolatum* | https://www.ncbi.nlm.nih.gov/pubmed/12439810 | mastitis |
| *Corynebacterium amycolatum* | https://www.ncbi.nlm.nih.gov/pubmed/12565065 | infective endocarditis |
| *Corynebacterium amycolatum* | https://www.ncbi.nlm.nih.gov/pubmed/15315020 | opportunistic infections |
| *Corynebacterium amycolatum* | https://www.ncbi.nlm.nih.gov/pubmed/15786829 | Peritonitis |
| *Corynebacterium amycolatum* | https://www.ncbi.nlm.nih.gov/pubmed/17284316 | endocarditis |
| *Corynebacterium amycolatum* | https://www.ncbi.nlm.nih.gov/pubmed/18174873 | infections in pediatric oncology |
| *Corynebacterium amycolatum* | https://www.ncbi.nlm.nih.gov/pubmed/18809563 | endocarditis |
| *Corynebacterium amycolatum* | https://www.ncbi.nlm.nih.gov/pubmed/19153032 | response to antibiotic tigecycline |
| *Corynebacterium amycolatum* | https://www.ncbi.nlm.nih.gov/pubmed/19876565 | predominant species in infections in cancer patients |
| *Corynebacterium amycolatum* | https://www.ncbi.nlm.nih.gov/pubmed/20624090 | resistance to antibiotic macrolide |
| *Corynebacterium amycolatum* | https://www.ncbi.nlm.nih.gov/pubmed/22361761 | clinical diphtheroid samples |
| *Corynebacterium amycolatum* | https://www.ncbi.nlm.nih.gov/pubmed/23806703 | surgical site infection |
| *Corynebacterium amycolatum* | https://www.ncbi.nlm.nih.gov/pubmed/26324578 | vaginosis |
| *Corynebacterium amycolatum* | https://www.ncbi.nlm.nih.gov/pubmed/28011352 | blood stream infection |
| *Corynebacterium amycolatum* | https://www.ncbi.nlm.nih.gov/pubmed/28264610 | breast abscess |
| *Corynebacterium amycolatum* | https://www.ncbi.nlm.nih.gov/pubmed/28700261 | infection of orbital implant |
| *Corynebacterium amycolatum* | https://www.ncbi.nlm.nih.gov/pubmed/29793964 | respiratory infection after lung transplant |
| *Corynebacterium amycolatum* | https://www.ncbi.nlm.nih.gov/pubmed/30102894 | Bloodstream and venous catheter-related infections |
| *Corynebacterium amycolatum* | https://www.ncbi.nlm.nih.gov/pubmed/30248572 | cystic neutrophilic granulomatous mastitis |
| *Corynebacterium amycolatum* | https://www.ncbi.nlm.nih.gov/pubmed/30803027 | bacteremia |
| *Corynebacterium amycolatum* | https://www.ncbi.nlm.nih.gov/pubmed/8727888 | clinical isolates, multiple sources |
| *Corynebacterium amycolatum* | https://www.ncbi.nlm.nih.gov/pubmed/8874085 | sepsis |
| *Corynebacterium amycolatum* | https://www.ncbi.nlm.nih.gov/pubmed/9157120 | neonatal sepsis fatal in premature infant |
| *Corynebacterium amycolatum* | https://www.ncbi.nlm.nih.gov/pubmed/9488824 | wound, bloodstream, and urinary tract infections |

| | | |
|---|---|---|
| *Corynebacterium amycolatum* | https://www.ncbi.nlm.nih.gov/pubmed/9505178 | infection after orthopedic surgery |
| *Corynebacterium amycolatum* | https://www.ncbi.nlm.nih.gov/pubmed/9868692 | Cardioverter-Lead Electrode Infection |
| *Corynebacterium lactis* | https://www.ncbi.nlm.nih.gov/pubmed/25937144 | Infection in companion dog |
| *Porphyromonas asaccharolytica* | https://www.ncbi.nlm.nih.gov/pmc/articles/PMC5896039/ | colorectal cancer |
| *Porphyromonas asaccharolytica* | https://www.ncbi.nlm.nih.gov/pmc/articles/PMC6247719/ | causes Lemierre's syndrome |
| *Porphyromonas asaccharolytica* | https://www.ncbi.nlm.nih.gov/pubmed/15528728 | clinical isolates |
| *Porphyromonas asaccharolytica* | https://www.ncbi.nlm.nih.gov/pubmed/15722627 | Clinical isolates, multiple sources |
| *Porphyromonas asaccharolytica* | https://www.ncbi.nlm.nih.gov/pubmed/15888469 | predominant in polymicrobial flora in 48 inflamed sinuses |
| *Porphyromonas asaccharolytica* | https://www.ncbi.nlm.nih.gov/pubmed/15897651 | Lemierre's syndrome |
| *Porphyromonas asaccharolytica* | https://www.ncbi.nlm.nih.gov/pubmed/16887693 | liver abscess |
| *Porphyromonas asaccharolytica* | https://www.ncbi.nlm.nih.gov/pubmed/19390440 | causes Lemierre's syndrome |
| *Porphyromonas asaccharolytica* | https://www.ncbi.nlm.nih.gov/pubmed/21407153 | tubo-ovarian abscess |
| *Porphyromonas asaccharolytica* | https://www.ncbi.nlm.nih.gov/pubmed/23435719 | causes Lemierre's syndrome (acute otopjaryngeal infection) |
| *Porphyromonas asaccharolytica* | https://www.ncbi.nlm.nih.gov/pubmed/23474186 | pleural empyema in immunocompetent diabetic patient |
| *Porphyromonas asaccharolytica* | https://www.ncbi.nlm.nih.gov/pubmed/24679105 | polymicrobial foot infection |
| *Porphyromonas asaccharolytica* | https://www.ncbi.nlm.nih.gov/pubmed/7548548 | extraoral infections |
| *Porphyromonas asaccharolytica* | https://www.ncbi.nlm.nih.gov/pubmed/7752213 | 418 children with infection, found in infections across body sites |
| *Porphyromonas asaccharolytica* | https://www.ncbi.nlm.nih.gov/pubmed/7857230 | cause chest wall abscess in one woman |
| *Porphyromonas asaccharolytica* | https://www.ncbi.nlm.nih.gov/pubmed/8126176 | bacterial vaginosis |
| *Porphyromonas asaccharolytica* | https://www.ncbi.nlm.nih.gov/pubmed/8518760 | male and female genital ulcers |
| *Porphyromonas asaccharolytica* | https://www.ncbi.nlm.nih.gov/pubmed/8907604 | female genital tract infection |
| *Porphyromonas asaccharolytica* | https://www.ncbi.nlm.nih.gov/pubmed/9200028 | intravenous catheter related bacteremia in child with cancer |
| *Porphyromonas asaccharolytica* | https://www.ncbi.nlm.nih.gov/pubmed/9772922 | infected cardiac myxoma |
| *Porphyromonas bennonis* | https://www.ncbi.nlm.nih.gov/pubmed/19542133 | identification and characterization in clinical specimen from various body sites |
| *Porphyromonas somerae* | https://www.ncbi.nlm.nih.gov/pubmed/16145091 | chronic skin, soft tissue and bone infections |
| *Porphyromonas somerae* | https://www.ncbi.nlm.nih.gov/pubmed/30541687 | abscesses, biopsies, wounds |
| *Porphyromonas uenonis* | https://www.ncbi.nlm.nih.gov/pubmed/15528728 | identification as pathogen |
| *Prevotella bivia* | https://www.ncbi.nlm.nih.gov/pubmed/10823756 | enhanced HIV expression |
| *Prevotella bivia* | https://www.ncbi.nlm.nih.gov/pubmed/10875323 | septic arthritis |
| *Prevotella bivia* | https://www.ncbi.nlm.nih.gov/pubmed/11368254 | bacterial vaginosis |
| *Prevotella bivia* | https://www.ncbi.nlm.nih.gov/pubmed/11707013 | septic arthritis |
| *Prevotella bivia* | https://www.ncbi.nlm.nih.gov/pubmed/14532256 | Paronychia |
| *Prevotella bivia* | https://www.ncbi.nlm.nih.gov/pubmed/15722627 | Clinical specimens |
| *Prevotella bivia* | https://www.ncbi.nlm.nih.gov/pubmed/16192439 | abdominal cutaneous ulcer |
| *Prevotella bivia* | https://www.ncbi.nlm.nih.gov/pubmed/16316686 | Lemierre's syndrome |
| *Prevotella bivia* | https://www.ncbi.nlm.nih.gov/pubmed/17367470 | virulence |
| *Prevotella bivia* | https://www.ncbi.nlm.nih.gov/pubmed/1747864 | bacterial vaginosis |
| *Prevotella bivia* | https://www.ncbi.nlm.nih.gov/pubmed/17982605 | penile abscess |

| | | |
|---|---|---|
| *Prevotella bivia* | https://www.ncbi.nlm.nih.gov/pubmed/18237241 | Chorionic plate inflammation |
| *Prevotella bivia* | https://www.ncbi.nlm.nih.gov/pubmed/19053926 | Oral lichen planus |
| *Prevotella bivia* | https://www.ncbi.nlm.nih.gov/pubmed/19271076 | septic arthritis |
| *Prevotella bivia* | https://www.ncbi.nlm.nih.gov/pubmed/19283879 | chest wall abscess |
| *Prevotella bivia* | https://www.ncbi.nlm.nih.gov/pubmed/20711427 | bacterial vaginosis in HIV infected women |
| *Prevotella bivia* | https://www.ncbi.nlm.nih.gov/pubmed/21214658 | amniotic fluid infection |
| *Prevotella bivia* | https://www.ncbi.nlm.nih.gov/pubmed/21376823 | Skin and soft tissue infection |
| *Prevotella bivia* | https://www.ncbi.nlm.nih.gov/pubmed/22375046 | inguinal bubo |
| *Prevotella bivia* | https://www.ncbi.nlm.nih.gov/pubmed/23001520 | Abdominal wall phlebitis following renal transplant |
| *Prevotella bivia* | https://www.ncbi.nlm.nih.gov/pubmed/24452170 | empyema |
| *Prevotella bivia* | https://www.ncbi.nlm.nih.gov/pubmed/24787738 | Pelvic inflammatory disease |
| *Prevotella bivia* | https://www.ncbi.nlm.nih.gov/pubmed/25114266 | Necrotizing fasciitis |
| *Prevotella bivia* | https://www.ncbi.nlm.nih.gov/pubmed/28008411 | Proctitis |
| *Prevotella bivia* | https://www.ncbi.nlm.nih.gov/pubmed/28903767 | bacterial vaginosis |
| *Prevotella bivia* | https://www.ncbi.nlm.nih.gov/pubmed/28931859 | bacterial vaginosis |
| *Prevotella bivia* | https://www.ncbi.nlm.nih.gov/pubmed/29772525 | bacterial vaginosis |
| *Prevotella bivia* | https://www.ncbi.nlm.nih.gov/pubmed/29860038 | multi-center survey of multi-drug resistant isolates |
| *Prevotella bivia* | https://www.ncbi.nlm.nih.gov/pubmed/8013486 | endocarditis |
| *Prevotella bivia* | https://www.ncbi.nlm.nih.gov/pubmed/8205934 | obstetrics gynecology specimen |
| *Prevotella bivia* | https://www.ncbi.nlm.nih.gov/pubmed/8270797 | association with cervical cancer |
| *Prevotella bivia* | https://www.ncbi.nlm.nih.gov/pubmed/8324131 | bacterial vaginosis in pregnant women |
| *Prevotella bivia* | https://www.ncbi.nlm.nih.gov/pubmed/8677085 | bacteremia after C-section |
| *Prevotella bivia* | https://www.ncbi.nlm.nih.gov/pubmed/8907604 | female genital tract infection |
| *Prevotella bivia* | https://www.ncbi.nlm.nih.gov/pubmed/9003606 | dog/cat bite wound |
| *Prevotella bivia* | https://www.ncbi.nlm.nih.gov/pubmed/9745330 | Periodontal abscesses |
| *Prevotella buccalis* | https://www.ncbi.nlm.nih.gov/pubmed/14662931 | urinary tract infection after renal transplant |
| *Prevotella buccalis* | https://www.ncbi.nlm.nih.gov/pubmed/24565649 | Endodontic infections |
| *Prevotella buccalis* | https://www.ncbi.nlm.nih.gov/pubmed/9266340 | Periodontitis |
| *Prevotella disiens* | https://www.ncbi.nlm.nih.gov/pubmed/15508748 | Periodontitis |
| *Prevotella disiens* | https://www.ncbi.nlm.nih.gov/pubmed/1747864 | bacterial vaginosis |
| *Prevotella disiens* | https://www.ncbi.nlm.nih.gov/pubmed/19161595 | bacterial vaginosis |
| *Prevotella disiens* | https://www.ncbi.nlm.nih.gov/pubmed/24565649 | Endodontic infections |
| *Prevotella disiens* | https://www.ncbi.nlm.nih.gov/pubmed/26183701 | cranioplasty infection |
| *Prevotella disiens* | https://www.ncbi.nlm.nih.gov/pubmed/8205934 | obstetrics gynecology specimen |
| *Prevotella disiens* | https://www.ncbi.nlm.nih.gov/pubmed/8324131 | bacterial vaginosis in pregnant women |
| *Prevotella disiens* | https://www.ncbi.nlm.nih.gov/pubmed/8907604 | female genital tract infection |
| *Prevotella timonensis* | https://www.ncbi.nlm.nih.gov/pubmed/17392225 | breast abscess |
| *Prevotella timonensis* | https://www.ncbi.nlm.nih.gov/pubmed/29307650 | various sites mostly genital and wound |

**Supplementary Figure 1. Correlation Network Analysis.**

**Dataset 2 Cases**

**a**

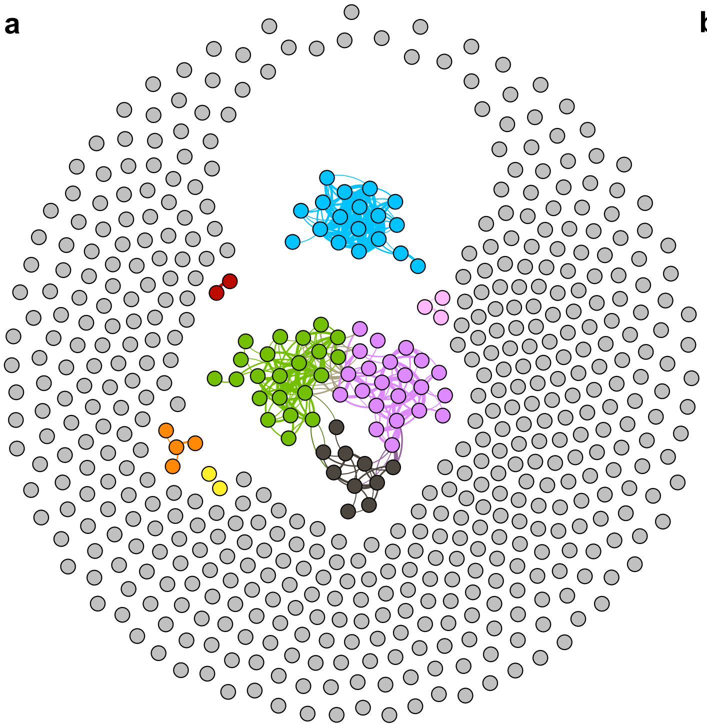

**b**

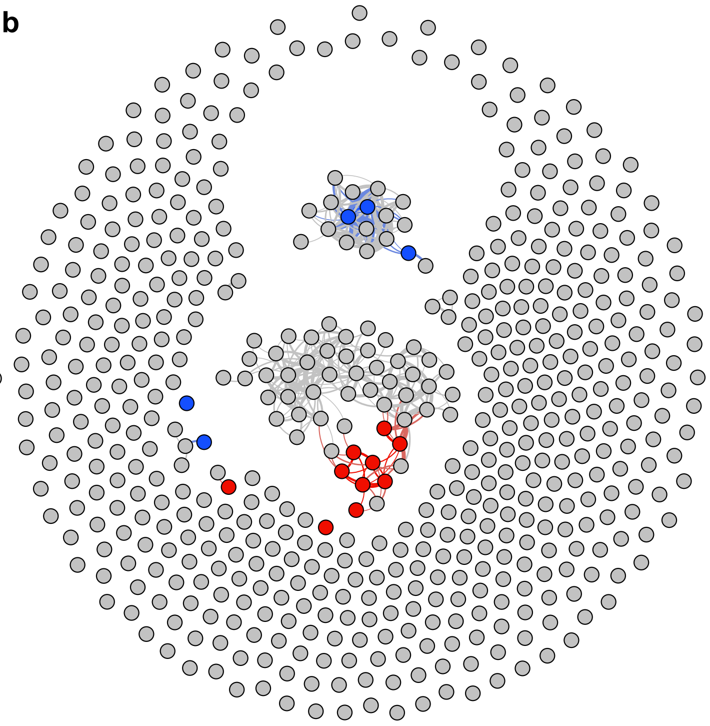

**Dataset 2 Controls**

**a**

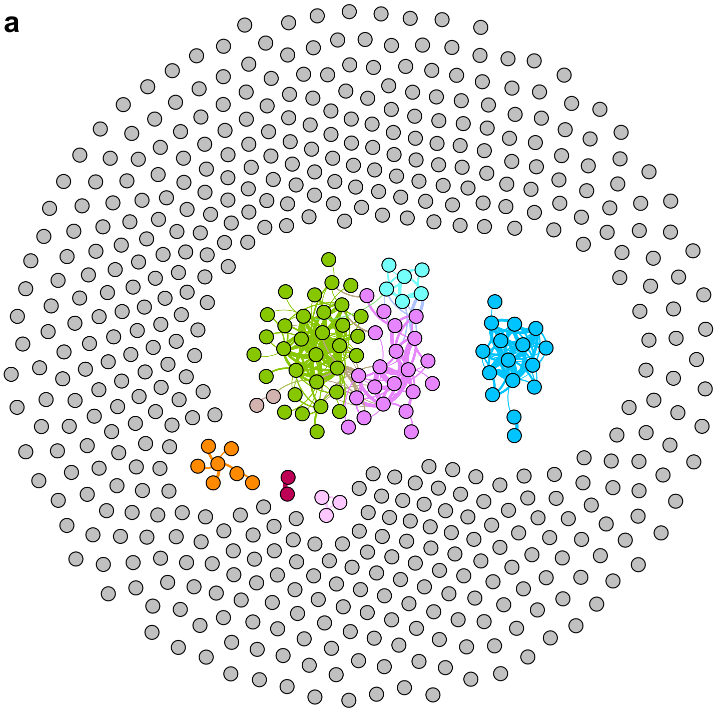

**b**

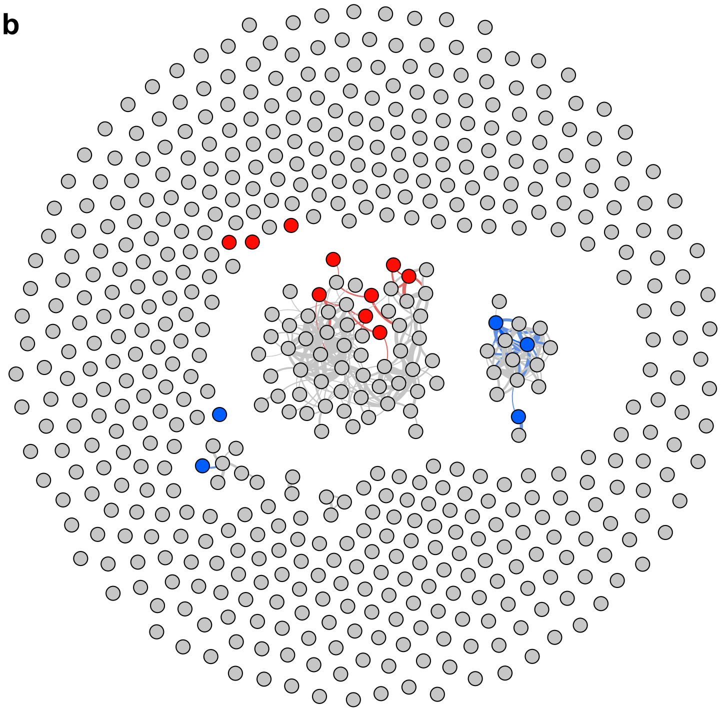

**Dataset 1 Cases**

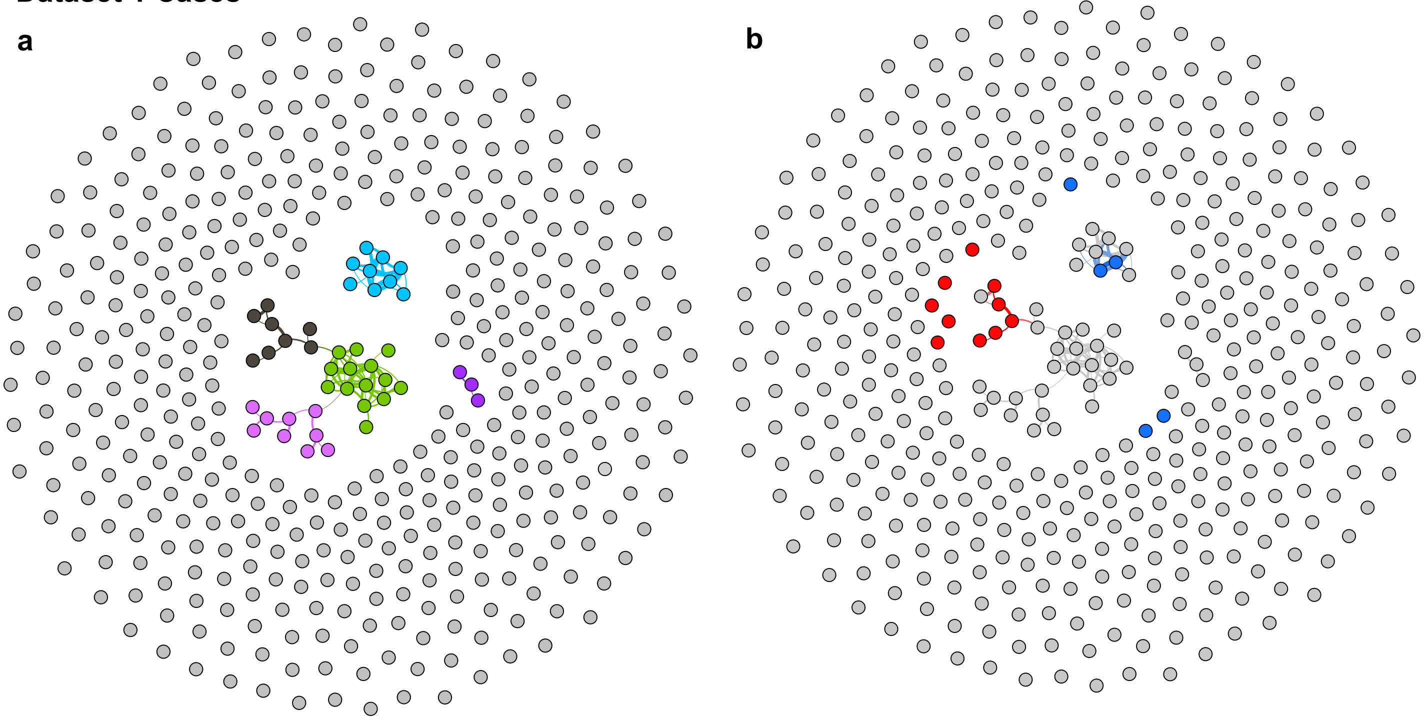

**Dataset 1 Controls**

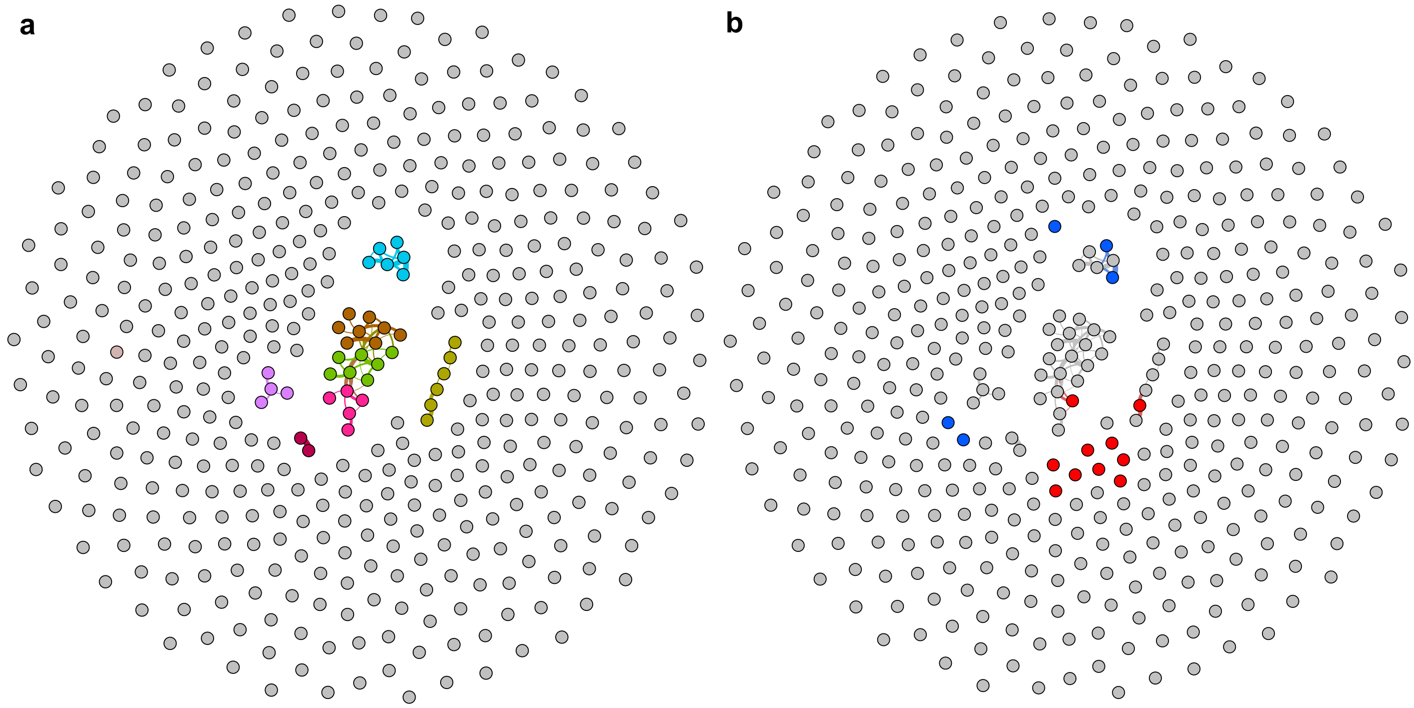

We calculated pairwise correlations in relative abundances for all genera microbiome-wide, for each dataset and in cases and controls separately. To display, we set an arbitrary threshold of correlation coefficient at r≥ |0.4| to connect genera that were correlated. At r≥ |0.4| all correlations were significant at P<3E-4 (the limit for 3,000 permutations). The graphics denoted by “a” display the algorithm-predicted clusters in different colors.

Graphics denoted by “b” are identical to their “a” counterpart except algorithm generated colors are now shown in grey and PD-associated taxa are highlighted in blue (if increased in PD) or red (if decreased in PD).  Dataset 2 has more power due to larger sample size, and has greater resolution due to deeper sequencing, nonetheless, the general patterns are similar in the two datasets. Generally, the 15 PD-associated genera fall in 3 clusters. As best seen in dataset 2 cases, which has the largest sample size and power, *Porphyromonas, Prevotella*, and *Corynebacterium_1* co-occur in cluster 1. Eight of the 10 in cluster 2 also connect at $r \geq |0.4|$, the other two, *Oscillospira* connects to cluster 2 at $r=0.25$ ($P<3E-4$) and *Lachnospiraceae_UCG-004* connects at $r=0.35$ ($P<3E-4$).  *Lactobacillus* and *Bifidobacterium* connect to each other (cluster 3) at $r=0.33$ ($P<3E-4$).
